# Targeted protein degradation via intramolecular bivalent glues

**DOI:** 10.1101/2023.02.14.528511

**Authors:** Oliver Hsia, Matthias Hinterndorfer, Angus D. Cowan, Kentaro Iso, Tasuku Ishida, Ramasubramanian Sundaramoorthy, Mark A. Nakasone, Hana Imrichova, Caroline Schätz, Andrea Rukavina, Koraljka Husnjak, Martin Wegner, Alejandro Correa-Sáez, Conner Craigon, Ryan Casement, Chiara Maniaci, Andrea Testa, Manuel Kaulich, Ivan Dikic, Georg E. Winter, Alessio Ciulli

## Abstract

Targeted protein degradation is a pharmacological modality based on the induced proximity of an E3 ubiquitin ligase and a target protein to promote target ubiquitination and proteasomal degradation. This has been achieved either via bifunctional compounds (PROTACs) composed of two separate warheads that individually bind the target and E3 ligase, or via molecular glues that monovalently bind either the ligase or the target^1–4^. Using orthogonal genetic screening, biophysical characterization, and structural reconstitution, we investigate the mode of action of bifunctional BRD2/4 degraders (IBG1-4) and find that – instead of connecting target and ligase *in trans* as PROTACs do – they simultaneously engage two adjacent domains of the target protein *in cis*. This conformational change glues BRD4 to the E3 ligases DCAF11 or DCAF16, leveraging intrinsic target-ligase affinities which, albeit pre-existing, do not translate to BRD4 degradation in absence of compound. Structural insights into the ternary BRD4:IBG1:DCAF16 complex guided the rational design of improved degraders of low picomolar potency. We thus introduce a new modality in targeted protein degradation, termed intramolecular bivalent glues (IBGs), which work by bridging protein domains to enhance surface complementarity with E3 ligases for productive ubiquitination and degradation.

## Main

### Sulfonamide-based PROTACs degrade BRD4 independently of DCAF15

The cullin-4 RING ligase (CRL4) substrate receptor DCAF15 is responsible for the pharmacologic degradation of the mRNA splicing factor RBM39 via the aryl sulfonamide molecular glues indisulam and E7820^5–9^. Efforts to leverage aryl sulfonamides as E3 binding ligands for PROTACs, including our own, have so far met with limited success (Extended Data Fig. 1a, b)^10–12^. However, a recent patent filing described a PROTAC-like degrader, herein referred to as IBG1 (Fig. 1a), consisting of the BET bromodomain inhibitor JQ1 tethered to E7820. IBG1 shows potent BRD4 degradation (DC_50_ = 0.15 nM) and pronounced growth inhibition in various cancer cell lines^13^. We synthesized IBG1 and confirmed efficient killing of diverse cell lines (Extended Data Fig. 1c) and BET protein degradation that was specific for BRD4 and BRD2 over their paralogue BRD3 (Fig. 1b, c, Extended Data Fig. 1d, e). The proteasome inhibitor MG132 and the neddylation inhibitor MLN4924 blocked BET protein degradation (Fig. 1d, Extended Data Fig. 1f), and MLN4924 also prevented BRD4 ubiquitination (Fig. 1e), indicating that IBG1 works via cullin-RING E3 ubiquitin ligase (CRL)-mediated ubiquitination and proteasomal degradation. Surprisingly, BRD4/BRD2 degradation was unaffected by DCAF15 knockout or knockdown (Fig. 1f, Extended Data Fig. 1g), suggesting an unexpected DCAF15-independent mode of degradation.

**Fig. 1:**
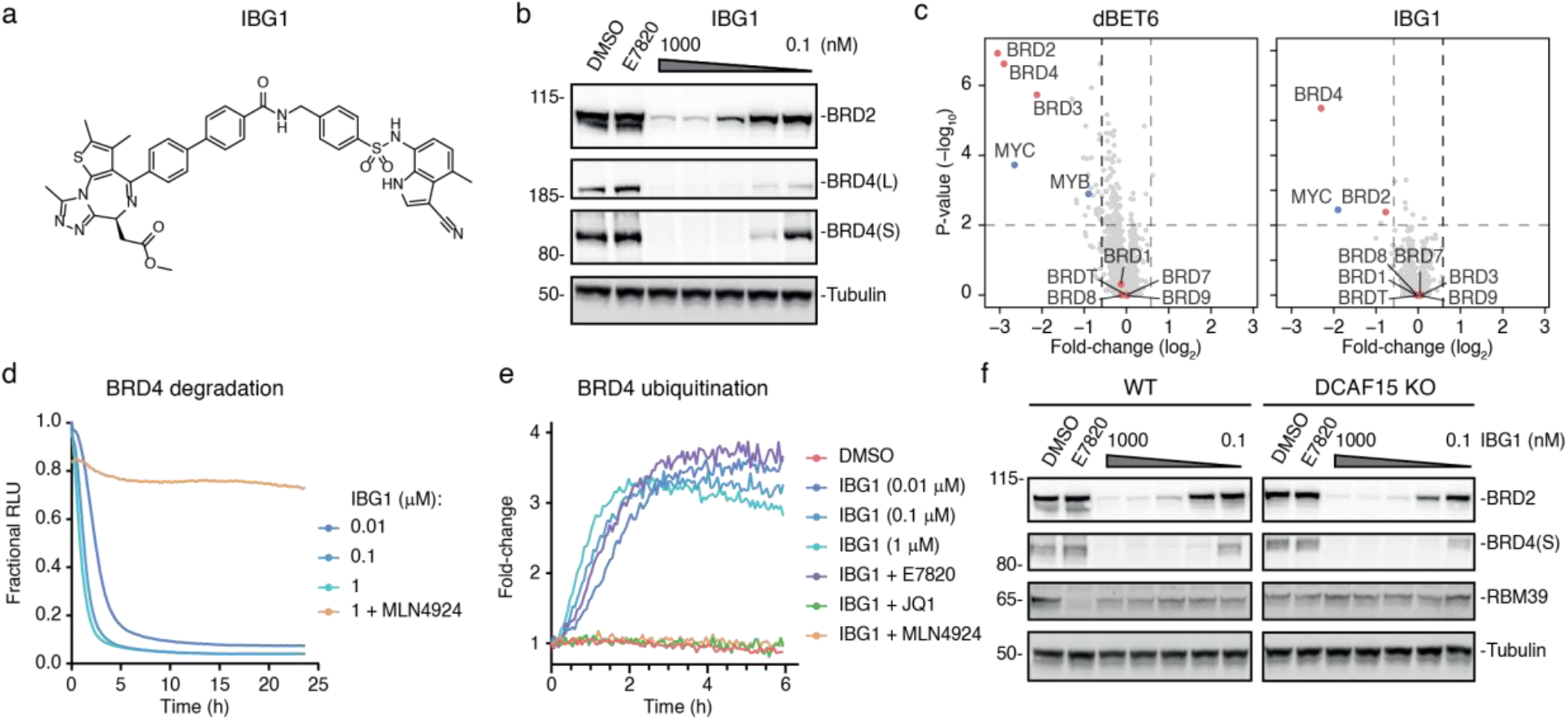
IBG1 degrades BRD2 and BRD4 independent of DCAF15. **a**, Structure of sulfonamide-based PROTAC IBG1. **b**, BET protein degradation activity of IBG1. HEK293 cells were treated for 6 hours with DMSO, E7820 (1000 nM) or increasing concentrations of IBG1 and BET protein levels were assessed via immunoblot. Western blot is representative of n = 3 independent experiments. **c**, Whole proteome changes after degrader treatment. Quantitative proteomics in KBM7 cells was performed after 6-hour treatment with DMSO, IBG1 (1 nM) or dBET6 (10 nM). Volcano plots show fold-change and significance over DMSO. n = 3 biological replicates. **d**, NanoBRET kinetic degradation assay. BromoTag-HiBiT-BRD4 knock-in HEK293 cells were treated with IBG1 at indicated concentrations with or without 1-hour MLN4924 (10 µM) pre-treatment. n = 3 biological replicates. **e**, NanoBRET kinetic ubiquitination assay. LgBiT-transfected HiBiT-BromoTag-BRD4 knock-in HEK293 cells were treated with IBG1 at indicated concentrations or at 10 nM following 1-hour pre-treatment with JQ1, E7820 (both 10 µM) or MLN4924 (1 µM). n = 4 biological replicates. **f**, DCAF15 independent BET protein degradation. HCT-116 WT and DCAF15 KO cells were treated with increasing concentrations of IBG1 for 6 h and BET protein levels were assessed via immunoblot. Data representative of n = 3 independent experiments.

### IBG1 recruits CRL4^DCAF^^16^ for the degradation of BRD4

To systematically identify the factors required for the degradation activity of IBG1, we set up a time-resolved FACS-based CRISPR/Cas9 BRD4 degradation screen (Fig. 2a). We engineered a dual fluorescence BRD4 protein stability reporter, consisting of a BRD4-mTagBFP fusion coupled to mCherry for normalization. We expressed this reporter in KBM7 cells harbouring a doxycycline (DOX)-inducible Cas9 allele^14^, transduced this cell line with a CRL-focused pooled sgRNA library^15^ and induced Cas9 expression with DOX. We then triggered BRD4 degradation via treatment with IBG1 or MZ1^16^ and used FACS to isolate cells with elevated (BRD4^HIGH^) or decreased (BRD4^LOW^) BRD4-BFP levels. In the DMSO control screen, we found 20S proteasome subunits, the COP9 signalosome, as well as the CRL3^SPOP^ complex to potently regulate BRD4 stability, recapitulating the known endogenous BRD4 turnover machinery^17,18^ (Fig. 2b). For MZ1, we identified subunits of the CRL2^VHL^ complex, including the CUL2 backbone, the adapters ELOB and ELOC, and the substrate receptor VHL, consistent with the known engagement of VHL by MZ1^16^ (Fig. 2b). This confirmed that our time-resolved screens can identify genes required for steady-state as well as induced BRD4 degradation, independent of gene essentiality. Next, we focused on the genes required for BRD4 degradation by IBG1. In line with our previous observations, our screens found the compound to work independently of DCAF15. Instead, we identified members of the CRL4^DCAF16^ complex, notably the CUL4A backbone, RBX1, the adapter DDB1 and the substrate receptor DCAF16, to be required for BRD4 degradation by IBG1, as recently reported for the monovalent BET degrader GNE-0011^19–22^ (Extended Data Fig. 2a). We furthermore identified DCAF16 alongside the CUL4-associated ubiquitin-conjugating enzyme UBE2G1^23,24^ as the top hits mediating resistance to IBG1 treatment in an orthogonal viability-based CRISPR screen (Fig. 2c).

**Fig. 2:**
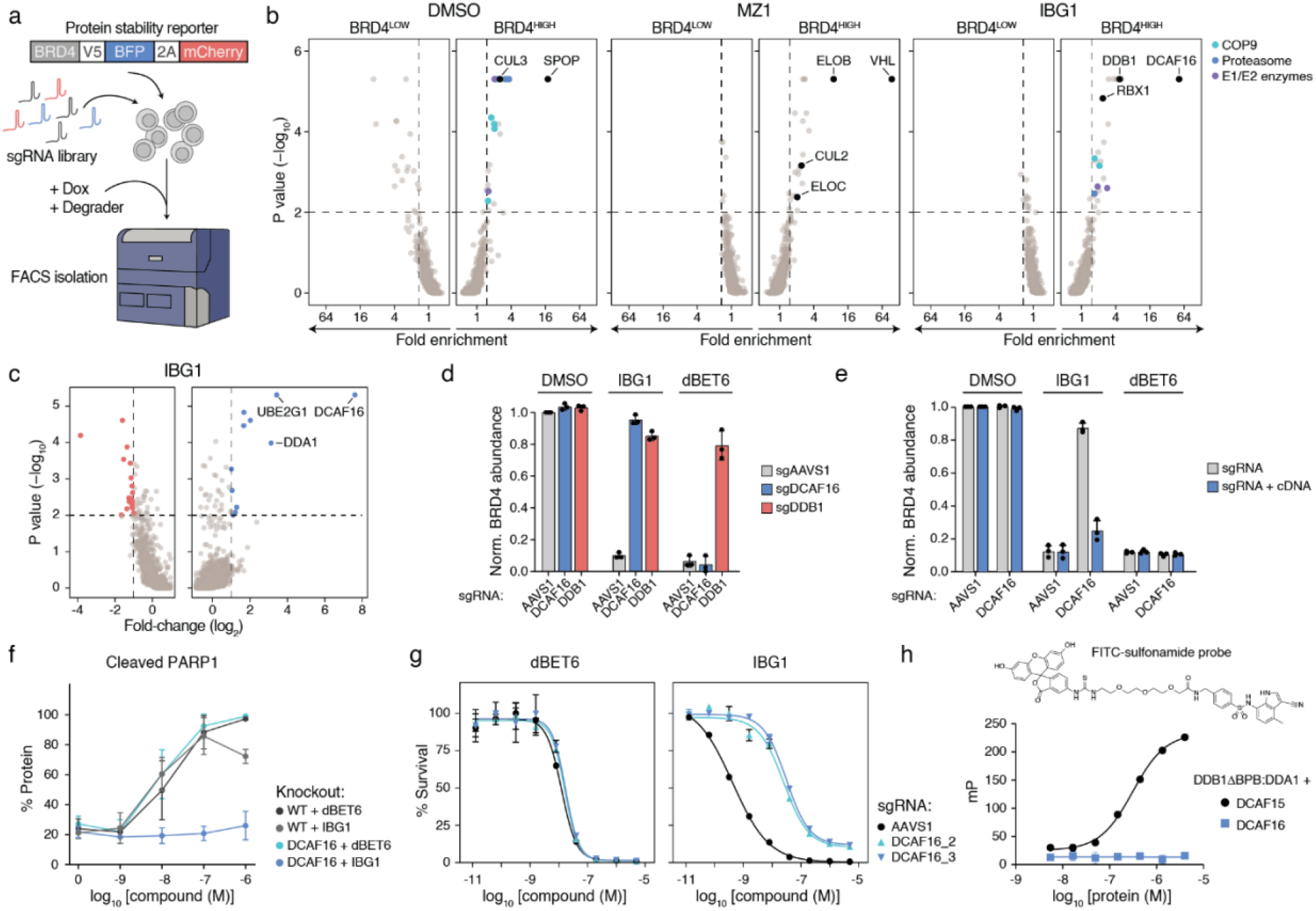
IBG1-induced BRD2/4 degradation is dependent on CRL4^DCAF16^. **a**, Schematic representation of dual fluorescence reporters and FACS-based CRISPR/Cas9 screens. KBM7 iCas9 BRD4-BFP protein stability reporter cells were transduced with a CRL-focused sgRNA library and Cas9 was induced via doxycycline for 3 days before cells were treated with BET degraders and sorted based on BRD4-BFP/mCherry ratios. **b**, FACS-based BRD4 stability CRISPR screens. sgRNA library-expressing KBM7 iCas9 BRD4 reporter cells were treated with DMSO, MZ1 (10 nM) or IBG1 (1 nM) for 6 hours before flow cytometric cell sorting. 20S proteasome subunits (blue), COP9 signalosome subunits (cyan) and E1 or E2 ubiquitin enzymes (purple) inside the scoring window (p-value < 0.01, fold-change > 1.5; dashed lines) are highlighted. **c**, CRISPR/Cas9 viability screen. HCT-116 cells were transduced with Cas9 and a UPS-focused sgRNA library and upon selection treated with IBG1 (58 nM; 4-fold IC_50_) for 6 days. Genes showing > 2-fold enrichment or depletion and p-value < 0.01 are highlighted. **d**, FACS-based CRISPR/Cas9 screen validation. KBM7 iCas9 BRD4-BFP reporter cells were transduced with AAVS1, DCAF16 or DDB1-targeting sgRNAs and treated with DMSO, IBG1 (1 nM) or dBET6 (10 nM) for 6 hours and BRD4-BFP levels were quantified via flow cytometry. **e**, DCAF16 KO/rescue. KBM7 iCas9 BRD4-BFP reporter cells were lentivirally transduced with AAVS1 or DCAF16-targeting sgRNAs, as well as a sgRNA resistant DCAF16 cDNA. After knockout of endogenous DCAF16, cells were treated for 6 hours as above and BRD4-BFP levels were quantified via flow cytometry. **f**, Apoptosis induction. KBM7 WT or DCAF16 KO cells were treated with indicated concentrations of dBET6 or IBG1 for 16 h. Levels of cleaved PARP1 were evaluated via immunoblotting. **g**, Viability assay. KBM7 iCas9 sgAAVS1 control or DCAF16 knockout cells were treated with increasing doses of IBG1or dBET6 for 72 hours and cell viability was evaluated by CellTiterGlo assay. Dose-response curves fitted using non-linear regression. n = 3 biological replicates, mean +/- s.d. **h**, Fluorescence polarization (FP) binary binding assay. FITC-labelled sulfonamide probe (top) was titrated into DCAF15:DDB1ΔBPB:DDA1 or DCAF16:DDB1ΔBPB:DDA1. n = 3 technical replicates. Data in **d**-**f**, n = 3 independent experiments; Data in **d**-**h**, mean +/- s.d.

In validation assays in KBM7 and HCT-116 cells, CRISPR-based knockout and siRNA-mediated knockdown of CRL4^DCAF16^ complex members prevented degradation of BRD4-BFP as well as endogenous BRD2 and BRD4 by IBG1, (Fig. 2d, Extended Data Fig. 2b-e), while ectopic expression of sgRNA-resistant DCAF16 restored degradation (Fig. 2e, Extended Data Fig. 2f). Finally, knockout of DCAF16 prevented the induction of apoptosis by IBG1 (Fig. 2f, Extended Data Fig. 2g) and led to enhanced tolerance of KBM7 cells (Fig. 2g), while IBG1 still induced efficient MYC downregulation, in line with retained DCAF16-independent BET bromodomain inhibition activity of its JQ1 warhead (Extended Data Fig. 2g). Together, these data show that despite the incorporation of a DCAF15-targeting aryl sulfonamide warhead^9^, IBG1 critically depends on the structurally unrelated CRL4 substrate receptor DCAF16 for BET protein degradation and cancer cell line efficacy. We thus investigated a potential affinity of IBG1 to DCAF16. As expected, we observed dose-dependent binding of a FITC-labelled E7820 probe to recombinant DCAF15, whereas it showed no affinity for DCAF16 (Fig. 2h). Additionally, the presence of excess amounts of E7820 or sulfonamide-containing truncations of IBG1 (compounds **1a**-**d**) did not prevent BRD4 ubiquitination or degradation by IBG1 (Fig. 1e, Extended Data Fig. 3a). These results indicated that the sulfonamide warhead is not involved in the recruitment of an E3 ligase in a PROTAC-like manner. However, IBG1 fragments harbouring truncations of the sulfonamide warhead (compounds **1e**-**g)** failed to promote BRD4 degradation despite efficient binding to BRD4 (Extended Data Fig. 3b, c), suggesting that the E7820 moiety is required for IBG1 activity, but in a role outside of direct E3 ligase recruitment.

### IBG1 enhances an intrinsic affinity between BRD4 and DCAF16

To further investigate the mechanism of action of IBG1, we sought to characterise the possible interactions between DCAF16, BRD4 and IBG1 *in vitro*. Using isothermal titration calorimetry (ITC), we observed the formation of a ternary complex between IBG1, DCAF16 and BRD4^Tandem^, a BRD4 construct containing both bromodomains (BD1 and BD2) connected by the native linker (K_D_ = 567 nM; Fig. 3a). Consistent with this, a time-resolved fluorescence transfer (TR-FRET) complex formation assay showed a dose-dependent ternary complex formed between DCAF16 and BRD4^Tandem^ in the presence of IBG1 (EC_50_ = 44 nM, Fig. 3b). A complementary TR-FRET based complex stabilization assay confirmed an interaction between DCAF16 and BRD4^Tandem^ in the presence of IBG1 (K_D_ = 712 nM), however, unexpectedly we also observed an intrinsic affinity of DCAF16 to BRD4^Tandem^ in the absence of IBG1 both in TR-FRET (K_D_ = 1 µM, Fig. 3c) and ITC (K_D_ = 4 µM, Extended Data Fig. 3d). Interestingly, no such intrinsic affinity was observed with isolated BRD4^BD1^. These observations were corroborated by size exclusion chromatography (SEC), where DCAF16 and BRD4^Tandem^ co-eluted in the absence of compound and this interaction was stabilised by IBG1, while again no interaction was observed with isolated BD1 and BD2 (Fig. 3d, e). In alphaLISA displacement assays, finally, we found significantly enhanced affinity of IBG1 to BRD4^Tandem^ in the presence of DCAF16 (IC_50_ = 12.8 nM) as compared to IBG1 and BRD4^Tandem^ alone (IC_50_ = 462 nM; cooperativity (α) = 36, Extended Data Fig. 3e), further supporting a role of IBG1 in the formation of a high-affinity BRD4:IBG1:DCAF16 ternary complex. Again, we observed no influence of DCAF16 on the binding of IBG1 to isolated BRD4^BD1^, corroborating that both bromodomains are required for complex formation. Together, these orthogonal assays establish an intrinsic affinity between BRD4 and DCAF16, which is stabilized by IBG1 and requires the presence of both bromodomains tethered together.

**Fig. 3:**
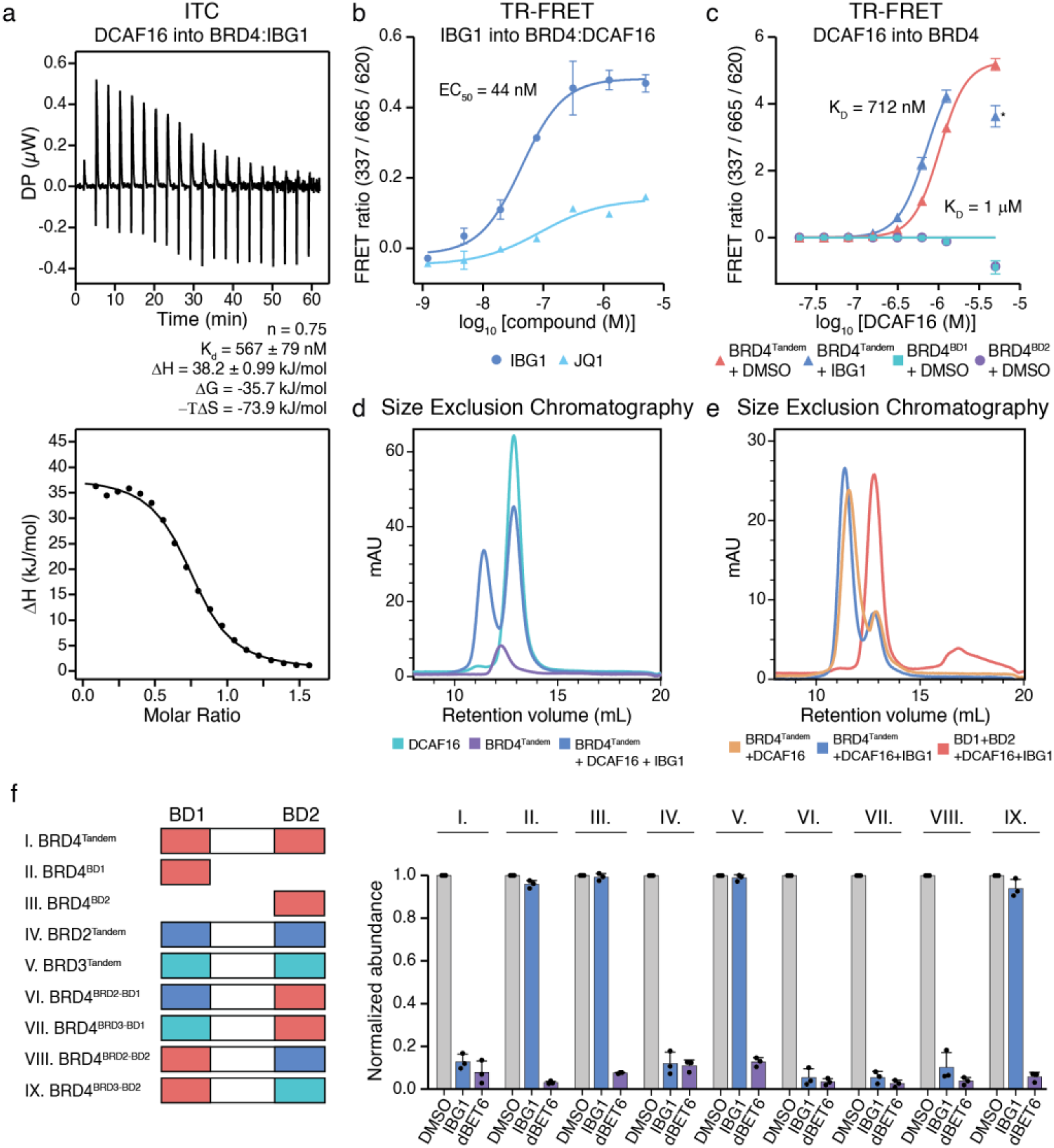
IBG1 enhances the intrinsic interaction between BRD4 tandem bromodomain region and DCAF16. **a**, ITC measurement of DCAF16:DDB1ΔBPB:DDA1 binding to pre-incubated BRD4^Tandem^:IBG1 complex (1:1.1 molar ratio). **b**, TR-FRET ternary complex formation assay. Anti-His-europium antibody bound to BRD4^Tandem^ was incubated with equimolar Cy5-labelled DCAF16:DDB1ΔBPB:DDA1 and increasing concentrations of IBG1 or JQ1. n = 3 technical replicates, mean +/- s.d. **c**, TR-FRET complex stabilization assay. 200 nM His-tagged BRD4^Tandem^ or BRD4^BD^^1^, bound to anti-His-europium antibody, were incubated with increasing concentrations of Cy5-labelled DCAF16:DDB1ΔBPB:DDA1 in the presence or absence of 1 µM IBG1. n = 2 independent experiments each with 2 technical replicates, mean +/- s.d. **d**, **e**, Size exclusion chromatography (SEC) UV chromatograms. DCAF16:DDB1ΔBPB:DDA1 and BRD4^Tandem^ alone or mixed at a 2:1 molar ratio in the presence of excess IBG1 (**d**), DCAF16:DDB1ΔBPB:DDA1 and BRD4^Tandem^ mixed at a 1:1 molar ratio in the absence or presence of excess IBG1 (**e**), as well as DCAF16:DDB1ΔBPB:DDA1 mixed with BRD4^BD^^1^ and BRD4^BD^^2^ at a molar ratio of 1:1:1 with excess IBG1 (**e**) were run on an S200 10/300 column. **f**, BET protein stability reporter assay. BRD2, BRD3 and BRD4 bromodomain tandems, as well as isolated BRD4 bromodomains and chimeras fused to mTagBFP were ectopically expressed in KBM7 cells and protein stability after 6-hour treatment with DMSO, IBG1 (1 nM) or dBET6 (10 nM) was quantified via flow cytometry. BD, bromodomain. n = 3 independent experiments, mean +/- s.d.

To further explore differential behaviour of individual bromodomains and BRD4^Tandem^, we focused on cellular assays based on a dual fluorescence BRD4 reporter. We generated a panel of KBM7 cell lines stably expressing either WT or truncated reporters (Fig. 3f, Extended Data Fig. 3f) and assessed the degradation of these constructs after treatment with IBG1 or dBET6 via flow cytometry. As expected, we observed potent degradation of WT BRD4(S) by both degraders. Serial deletion of the NPS, BID, ET and SEED domains did not affect degradation (Extended Data Fig. 3f) and the presence of BRD4^Tandem^ was sufficient for degradation (Fig. 3f). While the isolated BD1 and BD2 bromodomains were potently degraded by dBET6, we observed no degradation by IBG1 (Fig. 3f). Additionally, disruption of the JQ1 binding sites within the acetyl-lysine binding pockets in either bromodomain via single asparagine to phenylalanine changes (N140F or N433F, respectively) was sufficient to prevent degradation by IBG1, whereas simultaneous mutation of both bromodomains was required to disrupt dBET6-based degradation (Extended Data Fig. 3f). We furthermore utilized the BromoTag system, a ‘bump&hole’ L387A mutant BRD4^BD2^ degron tag^25^, to evaluate the degradation of a BromoTag-MCM4 fusion and observed potent degradation by the ‘bumped’ VHL-based PROTAC AGB1, while a similarly ‘bumped’ derivative of IBG1 (bIBG1) failed to induce any degradation (Extended Data Fig. 3g, h). Together, these data confirm that, unlike for most classical BET PROTACs, a single BRD4 bromodomain is not sufficient to trigger degradation by IBG1. Instead, it requires the simultaneous engagement of both acetyl-lysine binding pockets to induce degradation.

Another feature that distinguishes IBG1 from many other BET protein degraders is its specificity for BRD2 and BRD4 over BRD3 (Fig. 1c, Extended Data Fig. 1b, c). We therefore employed the FACS-based protein stability assay to identify the features determining this specificity. Consistent with their effects on endogenous proteins, we observed potent degradation of BRD2, BRD3 and BRD4 tandem constructs by dBET6, whereas IBG1 selectively degraded BRD2^Tandem^ and BRD4^Tandem^ while not affecting BRD3^Tandem^ (Fig. 3f). When we exchanged the linker from BRD4^Tandem^ with the corresponding regions in BRD2 and BRD3, we observed no influence on degradation (Extended Data Fig. 3g). We also ruled out a role of the known SPOP degron within the linker region in BRD4 (Extended Fig. 3g). Next, we swapped either BD1 or BD2 from BRD4^Tandem^ with the corresponding domain from BRD2 or BRD3. While exchange of the first bromodomain had minimal influence on protein degradation, for the second bromodomain only a swap with BRD2 was tolerated. In contrast, replacement by the BRD3^BD2^ fully disrupted degradation by IBG1 (Fig. 3f). Thus, the second bromodomain determines the selectivity of IBG1 for BRD2 and BRD4 over BRD3.

### IBG1 bivalently binds both bromodomains to glue BRD4 to DCAF16

To gain molecular insights into the mechanism underpinning IBG1-induced BRD4 degradation, we solved the structure of the ternary complex formed between BRD4^Tandem^, IBG1 and DCAF16:DDB1ΔBPB:DDA1 by cryo-electron microscopy at ∼3.77 Å resolution (Fig. 4a, Extended Data Fig. 4). DCAF16 adopts a unique fold consisting of 8 helices, several loops and a structural zinc ion coordinated by residues C100 and C103 in the loop between α3 and α4 and C177 and C179 of α8 (Extended Data Fig. 5a). Helices 4-6 bind the central cleft between β-propeller A (BPA) and C (BPC) of DDB1 in a binding mode distinct from other DCAF proteins and other CRL4 substrate receptors^7,26–28^ (Extended Data Fig. 5b). Helices 1, 3, 7 and 8 fold into a helical bundle that sits on the outer surface of BPC blades 5 and 6, as well as the loop between strands c and d of blade 7. Consistent with its role as CRL substrate receptor, this helical bundle of DCAF16 bridges the DDB1 adaptor protein with the neosubstrate BRD4. Remarkably, we found both bromodomains simultaneously bound to DCAF16 with a single continuous piece of density representing one molecule of IBG1 sitting between DCAF16, BD1 and BD2 (Fig. 4b). While the JQ1 moiety of IBG1 binds canonically to the acetyl-lysine binding pocket of BD2, the E7820 moiety unexpectedly binds to the equivalent pocket of BD1. The binding mode of the E7820 portion of IBG1 overlays well with other sulfonamide-containing BET inhibitors that have been co-crystallized with BD1^29,30^, with the nitrogen atom of the cyano group taking a position that is consistently occupied by a conserved water molecule in BET bromodomain crystal structures^31^ (Extended Data Fig. 5c). In line with these observations, we found that E7820 and other aryl-sulfonamide derivatives exhibit weak binding affinity to BRD4^Tandem^ as well as isolated bromodomains (Extended Data Fig. 5d, e). In SEC we observed increased retention of IBG1-bound BRD4^Tandem^ compared to apo- or JQ1-bound BRD4^Tandem^, indicating a decrease in hydrodynamic radius consistent with compaction of BRD4 through intramolecular dimerization of bromodomains induced by IBG1 (Fig. 4c). Thus, both BRD4 bromodomains are simultaneously engaged and bridged by the opposing ends of a single IBG1 molecule.

**Fig. 4:**
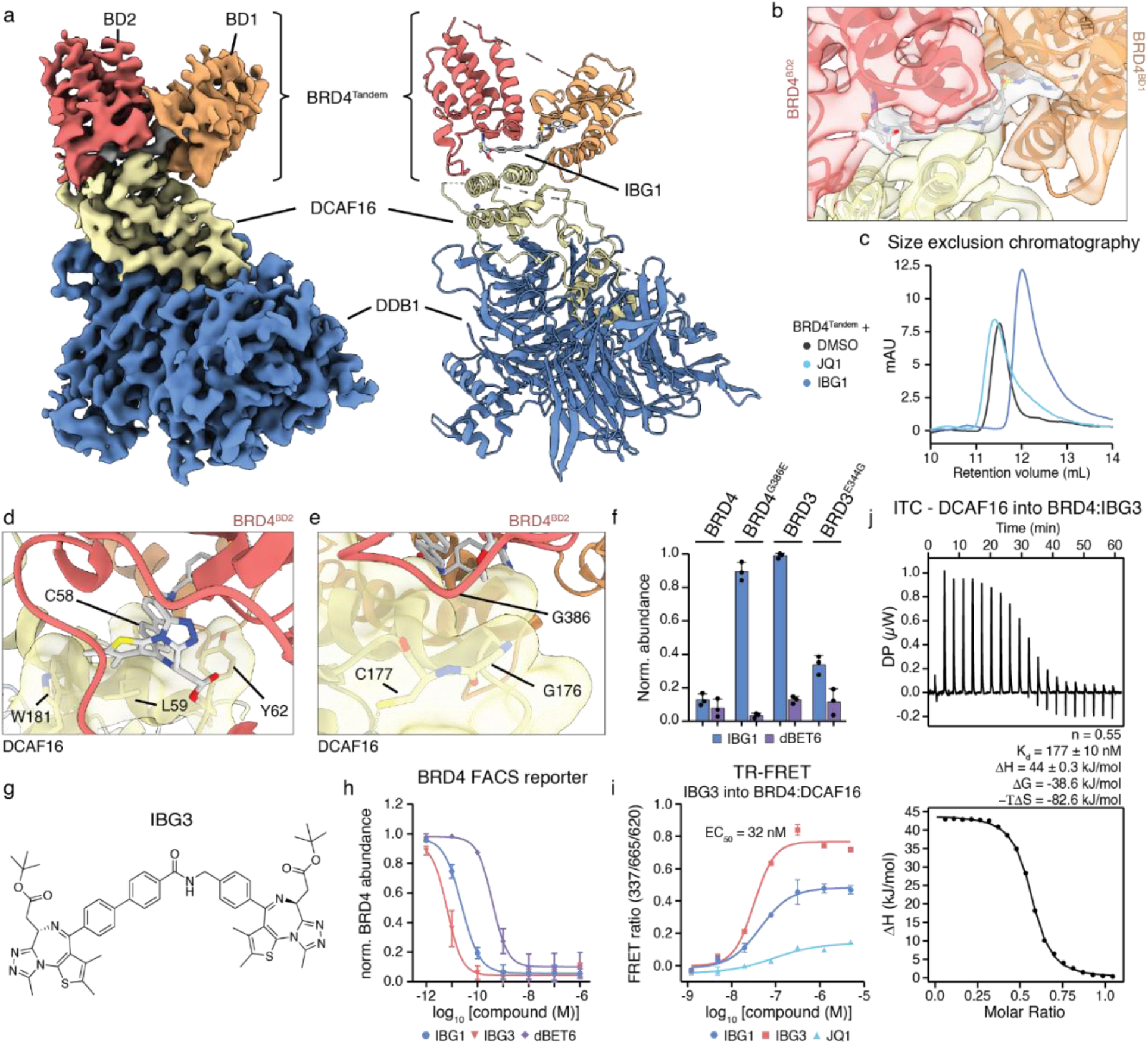
IBG1 intramolecularly engages both BRD4 bromodomains simultaneously and glues BRD4 to DCAF16. **a**, Electron density (left) and model (right) for the complex formed between DCAF16 (yellow), DDB1ΔBPB (blue), BRD4^Tandem^ (BD1 orange, BD2 red) and IBG1 (grey). **b**, Electron density at the DCAF16:IBG1:BRD4 interface. The JQ1 moiety binds to BD2 while the sulfonamide engages BD1. **c**, A hydrophobic cage formed by DCAF16 residues C58, L59, Y62 and W181 encloses the JQ1 moiety and linker phenyl ring of IBG1. **d**, Selectivity determining residue G386 of BD2 at the interface with DCAF16. For **b-d**, colours as in **a**. **e**, FACS reporter assay. WT BRD3 and BRD4 as well as single point mutant bromodomain tandem dual fluorescence reporter KBM7 cells were treated with IBG1 (1 nM) or dBET6 (10 nM) for 6 hours and BET protein stability was evaluated via flow cytometry. n = 3 independent experiments, mean +/- s.d. **f**-**j**, Structure (**f**) and mechanistic characterization of dual-JQ1 containing intramolecular glue BRD4 degrader IBG3. **h**, BRD4 degradation. BRD4^Tandem^ dual fluorescence KBM7 reporter cells were treated for 6 hours with increasing concentrations of IBG1, IBG3 or dBET6 and BRD4 protein stability was assessed via flow cytometry. n = 3 independent experiments, mean +/- s.d. **i**, TR-FRET ternary complex formation assay. Anti-His-europium antibody bound to BRD4^Tandem^ was incubated with equimolar Cy5-labelled DCAF16:DDB1ΔBPB:DDA1 and increasing concentrations of IBG1, IBG3 or JQ1. Data for JQ1, IBG1 as in Fig. 3b. n = 3 technical replicates, mean +/- s.d. **j**, ITC measurements of DCAF16:DDB1ΔBPB:DDA1 complex binding to pre-incubated BRD4^Tandem^:IBG3 (1:1.1 molar ratio).

At the ternary interface, DCAF16 encloses the hydrophobic dimethylthiophene and phenyl groups of JQ1 as well as the linker phenyl, shielding them from solvent (Fig. 4d). DCAF16 also contacts BD1 through residue W54, which binds into a hydrophobic pocket on the surface of BD1 (Extended Data Fig. 5f). The ternary complex is furthermore stabilized by intramolecular contacts between the two bromodomains, including the sandwiching of M442 between W81 and P375 in the WPF shelves of BD1 and BD2, respectively (Extended Data Fig. 5g). G386 of BD2 is positioned at a crucial interface in close contact with DCAF16, with only limited space available for the amino acid side chain (Fig. 4e). Interestingly, the corresponding residue in BRD2 is also a glycine (G382), while in BRD3 it is a glutamate (E344), suggesting a role of this residue in determining the selectivity of IBG1 for BRD2 and BRD4. Indeed, engineering a G386E mutation in the BRD4^Tandem^ FACS reporter completely abrogated degradation by IBG1, likely due to steric clashes with DCAF16, while the reciprocal E344G mutation in BRD3 sensitized it to degradation by IBG1 (Fig. 4f).

Based on this structurally elucidated mode of action, we hypothesized that bifunctional compounds harbouring two high-affinity bromodomain ligands should stabilize this conformation even more efficiently, potentially enabling the generation of more effective DCAF16-based intramolecular bivalent glue degraders. We thus synthesized a series of compounds in which we replaced E7820 by a second JQ1 moiety while keeping the linker architecture of IBG1 intact (compounds IBG2 and IBG3, Fig 4g, Extended Fig. 6a) and found that BRD4 and BRD2 degradation efficiencies indeed exceeded those of the original compound IBG1, with the most potent compound IBG3 showing degradation in a low picomolar range (DC50 = 6.7 and 8.6 pM, respectively, Fig. 4h, Extended Fig. 6b, c). Compared to IBG1, IBG3 also showed improved gluing of the BRD4:DCAF16 complex in a TR-FRET ternary complex formation assay (EC_50_ = 32 nM, Fig. 4i), increased affinity of DCAF16 for BRD4:IBG3 in isothermal titration colorimetry (ITC) experiments (Fig. 4j) and more pronounced compaction of tandem bromodomains in SEC (Extended Data Fig. 6d). Like its parental compound, degradation by IBG3 was specific for BRD4 and BRD2 over BRD3 (Extended Data Fig. 6b), selective for bromodomain tandems over isolated BRD4 bromodomains (Extended Data Fig. 6e), and mediated by DCAF16 (Ext. Data Fig. 6f, g), indicating degradation via the conserved intramolecular glue mechanism. BRD4 constructs harbouring two copies of either BD1 or BD2 were fully resistant to degradation by both IBG1 and IBG3 (Extended Data Fig. 6h), supporting the importance of the explicit relative arrangement of the bromodomains for ternary complex architecture. This also likely explains the functional difference to previously published bivalent bromodomain-targeting compounds, such as MT1 and MS645, that work as potent inhibitors without inducing BET protein degradation^32–34^ (Extended Data Fig. 6i, j). Thus, based on mechanistic and structural insights into the mode of action of IBG1, we rationally designed IBG3 as an improved intramolecular bivalent glue degrader with higher efficacy than any other degrader reported to date to our knowledge.

### IBG4 is a CRL4^DCAF11^-dependent intramolecular bivalent glue degrader

As bridging of two domains on BRD4 induced a potent *gain of function* outcome through stabilizing interactions to DCAF16, we surmised that protein surface adaptation via bivalent domain engagement might harbour the potential to induce or stabilize a spectrum of additional PPIs with different functional outcomes and mechanistic manifestations. Thus, we sought to generalize the concept of intramolecular bivalent glue degraders. To this end, a recently disclosed BRD4 degrader consisting of a pyrazolo pyrimidine warhead connected to JQ1 via a short rigid linker^35^ (herein termed IBG4, Fig. 5a) caught our attention since it, akin to IBG1, showed efficient degradation of BRD4^Tandem^, while sparing isolated bromodomains and BRD4 acetyl-lysine binding pocket mutants N140F and N433F (Fig. 5b, Ext. Data Fig. 7a). In size exclusion chromatography, IBG4 induced a similar compaction of BRD4^Tandem^ as IBG1 (Fig. 5c). In NanoBRET conformational biosensor assays^36^ both compounds induced comparable levels of intramolecular bromodomain interactions (Extended Data Fig. 7b), together indicating that IBG4 induces bromodomain dimerization *in cis,* similar to IBG1. Finally, the pyrazolo pyrimidine warhead of IBG4 showed comparable affinity to BRD4 bromodomains as the E7820 moiety in IBG1 (Extended Data Fig. 7c). Thus, despite being structurally differentiated, IBG4 phenotypically mimics the cellular mode of action of IBG1, suggesting that both compounds may work via a similar intramolecular glue-like mode of action. Unlike IBG1, however, IBG4 showed high specificity for BRD4 and did not efficiently degrade BRD2 (Ext. Data Fig 7d), pointing towards different structural requirements of a potential ternary BRD4:IBG4:E3 ligase complex. Indeed, while degradation was blocked by the neddylation inhibitor MLN4924 (Extended Data Fig. 7e), DCAF16 knockout had no effect on IBG4-mediated BRD4 degradation (Extended Data Fig. 7). We thus performed a CRISPR/Cas9 BRD4 degradation screen and besides the endogenous BRD4 turnover factor SPOP identified the CRL4^DCAF11^ complex as strongest hit mediating resistance to IBG4-induced degradation (Fig. 5d, e). As observed for DCAF16 and despite no predicted structural similarity, DCAF11 showed measurable intrinsic affinity for BRD4 in TR-FRET (Fig. 5f). Again, this interaction was significantly enhanced in the presence of IBG4 (Figure 5f, g). Finally, in line with stabilization of the ternary complex, the addition of IBG4 induced co-elution of BRD4 with DCAF11 in SEC (Extended Data Fig. 7g, h). In conclusion, IBG4 recapitulates all cellular and biophysical properties of the above-described intramolecular glue degraders, but extends the mechanistic scope to another, structurally unrelated, E3 ligase. Collectively, our data establish intramolecular dimerization of protein domains as a novel and generally applicable strategy for targeted protein degradation that can be rationally engineered following principles of structure-based drug design.

**Fig. 5:**
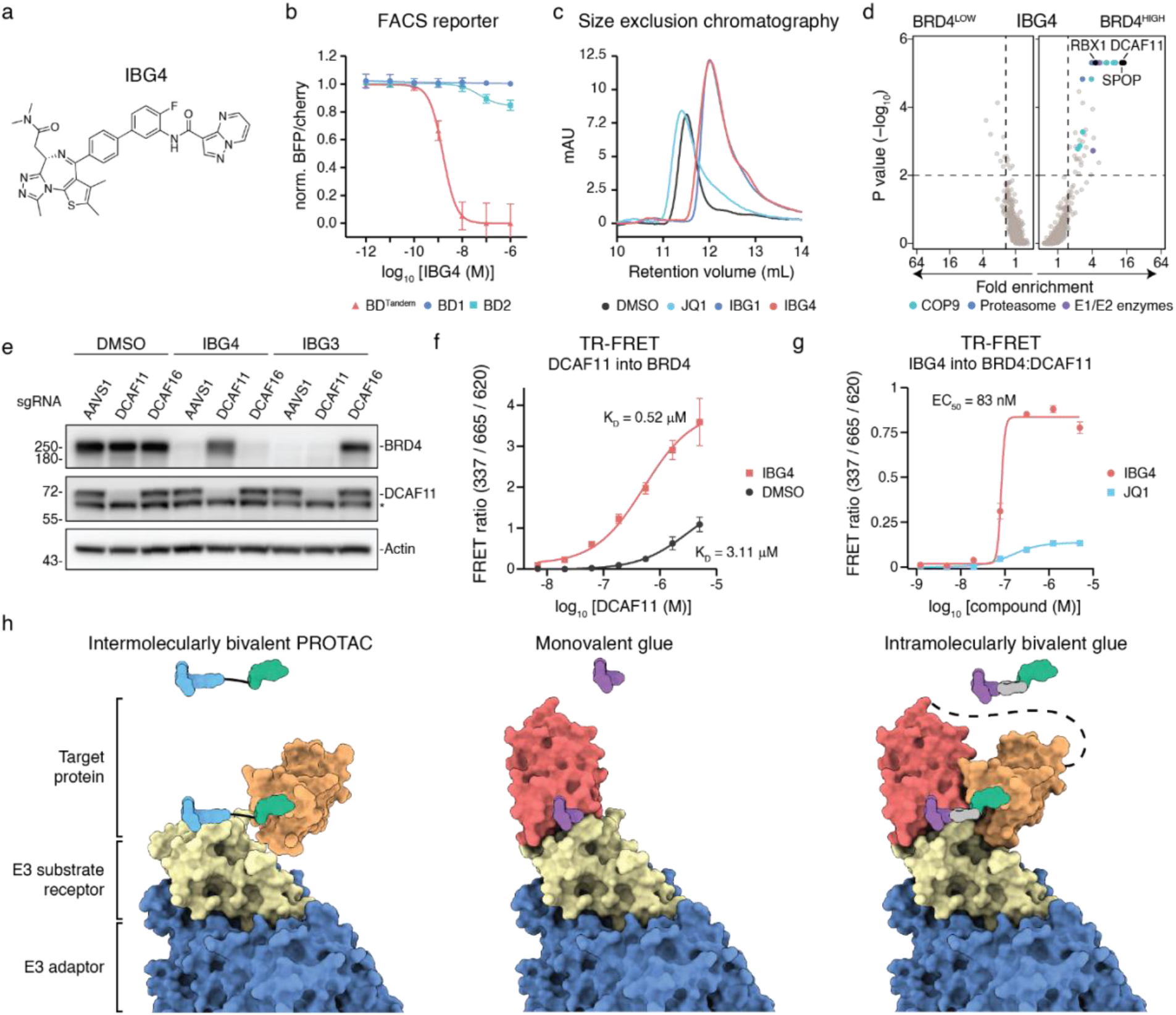
IBG4 is a DCAF11-dependent intramolecular bivalent glue degrader. **a**, Structure of BET degrader IBG4. **b**, Bromodomain tandem requirement of IBG4. BRD4^Tandem^, BD1 or BD2 dual fluorescence reporter KBM7 cells were treated for 6 hours with increasing concentrations of IBG4 and protein degradation was evaluated via flow cytometry. n = 3 independent experiments, mean +/- s.d. **c**, Size exclusion chromatograms of BRD4^Tandem^ incubated with DMSO, JQ1, IBG1 or IBG4. Data for DMSO, JQ1 and IBG1 as in Fig 4c. **d**, BRD4 stability CRISPR screen. KBM7 iCas9 BRD4 dual fluorescence reporter cells expressing a CRL-focused sgRNA library were treated with IBG4 (100 nM) for 6 hours before flow cytometric cell sorting into BRD4^LOW^, BRD4^MID^ and BRD4^HIGH^ fractions as in Fig. 2b. 20S proteasome subunits (blue), COP9 signalosome subunits (cyan) and E1 or E2 ubiquitin enzymes (purple) inside the scoring window (p-value < 0.01, fold-change > 1.5; dashed lines) are highlighted. **e**, Immunoblot-based screen validation. AAVS1 control, DCAF11 or DCAF16 knockout KBM7 cells were treated with DMSO, IBG4 (100 nM) or IBG3 (0.1 nM) for 6 hours and BRD4 levels were analysed via immunoblotting. * denotes unspecific band. Data are representative for n = 3 independent experiments. **f**, TR-FRET complex stabilization assay. 100 nM His-tagged BRD4^Tandem^ bound to anti-His europium antibody, was incubated with increasing concentrations of Cy5-labelled DCAF11:DDB1ΔBPB:DDA1 and 500 nM IBG4 or DMSO. Data shown are from n = 2 independent experiments, each with 3 technical replicates, mean +/- s.d. **g**, TR-FRET complex formation assay. His-tagged BRD4^Tandem^ bound to anti-His europium antibody was incubated with equimolar Cy5-labelled DCAF11:DDB1ΔBPB:DDA1 and increasing concentrations of IBG4 or JQ1. Data shown are from n = 3 technical replicates, mean +/- s.d. **h**, Schematic model of the different molecular recognition with traditional monovalent glues and bivalent PROTACs vs. intramolecularly bivalent glue as revealed in this work.

## Discussion

Most molecular glue degraders reported to date preferentially engage the E3 ligase at a binary level, and subsequently recruit target proteins. This notion underpins physiological regulatory circuits, such as the auxin-induced degradation of the IAA transcriptional repressor proteins in plant development^37^. It is also leveraged therapeutically by monovalent molecular glue degraders such as lenalidomide and the related immunomodulatory drugs (“IMiDs”) that induce the degradation of C2H2 zinc finger transcription factors (TFs) by binding to the E3 ligase CRBN and triggering ligase-TF interactions^28,38–41^. Such a mechanism, however, means that only targets that can be productively paired to a chemically accessible ligase can be actioned via this strategy. In contrast, very few glues to date have been developed from a given target protein binding compound^42,43^. Here, we define the mechanism of chemically distinct BET protein degraders as simultaneously engaging two separate sites on the target protein to nucleate formation of stable ternary complexes and induce target protein degradation. Hence, we reveal a new strategy distinct from conventional bivalent PROTACs and monovalent glues, termed ‘intramolecular bivalent gluing’, that enables the development of potent and target-selective degraders (Fig. 5g). Based on our mechanistic and structural insights, we rationally improved the first-generation intramolecular bivalent glue degrader IBG1 by enhancing its affinity to tandem bromodomains and glueing to DCAF16. This resulted in the second-generation IBG3, that showed half-maximal degradation at single digit picomolar concentrations, demonstrating that this novel class of degraders can reach efficiencies higher than any PROTAC reported to date^44^.

Around 60-80% of all human proteins feature at least two distinct domains and are hence potentially accessible to targeted degradation via intramolecular bivalent gluing^45,46^. Remarkably, both IBG1 and IBG4 feature only a single high-affinity BET ligand, while the second warhead shows only low affinity for its respective target protein domain. Nevertheless, both compounds trigger rapid and profound degradation at nanomolar concentrations, suggesting that these glues can efficiently degrade target proteins even when utilizing suboptimal secondary ligands. Even though the intramolecular bivalent glue degraders discovered here are currently focused on a single family of target proteins, these relatively lenient requirements for target binding suggest that this approach might be applicable for a much broader range of targets. Conversely, our work also highlights the challenges of using sub-specific or weak-affinity ligands, such as E7820, as E3 binding “handles” for conventional PROTAC mechanism, thus offering a cautionary tale as the field expands to E3 ligases beyond CRBN and VHL.

Despite showing that IBG1 and IBG4 work via highly similar modes of action, we find that they utilize two structurally unrelated E3 ligases to induce protein ubiquitination and degradation: while IBG1 functions via CRL4^DCAF16^, IBG4 relies on CRL4^DCAF11^. We identified intrinsic affinities between BRD4 and either of the two E3 ligases even in the absence of ligands. This reinforces the emerging concept that molecular glue degraders often stabilize pre-existing, albeit functionally inconsequential E3-target interactions^43,47,48^ and further suggests that these affinities may be essential for degradation via intramolecular glue degraders. The exclusive requirement of IBG1 and IBG4 on DCAF16 and DCAF11, respectively, suggests that the varying warhead arrangement and linker architecture align the BRD4 bromodomains in different orientations relative to each other, generating distinct protein-ligand surfaces that are selectively recognized by the two ligases. The structural studies undertaken herein ålso highlight the importance of linker rigidity and lipophilicity for the gluing between the target and the ligase and represent both a key focal point for further discovery and optimisation of intramolecular bivalent glues. Together, our work supports a model in which both DCAF11 and DCAF16 are primed for BET bromodomain recognition and that relatively mild modifications of the BET protein interaction surface could be sufficient to trigger productive complex stabilization and ubiquitination. This apparent affinity of BRD4 to various E3 ligases might be a potential explanation for the eminent accessibility of BET proteins for chemically induced protein degradation^49^.

In conclusion, we elucidated the mode of action of structurally distinct BET protein degraders that converge on a shared novel mode of action: intramolecular dimerization of two protein domains to potently induce ubiquitination and proteasomal degradation. This concept of modulating the surface of a target protein by an intramolecular, chemical bridging of two binding sites *in cis* could outline a generalizable strategy to pharmacologically induce proximity with E3 ligases or other cellular effector proteins with intrinsic affinity potential, thus paving the way for the innovation of chemical strategies to modulate or rewire cellular circuits.

## Acknowledgements

We are grateful to all members of the Ciulli and Winter laboratories and to Juraj Konc for experimental advice and helpful discussions. From the Ciulli lab, we particularly want to thank Sarath Ramachandran, Andre Wijaya, Zoe Rutter, Thomas Webb, and Valentina Spiteri for assistance with biophysical assay troubleshooting and productive discussions. We acknowledge the European Synchrotron Radiation Facility (ESRF) for provision of synchrotron radiation facilities, and we would like to thank Max Nanao for assistance and support in using beamline ID23-2. We thank Johannes Zuber and members of the Zuber laboratory at the Research Institute of Molecular Pathology for sharing iCas9 cell lines, reagents and plasmids. In particular, we thank Florian Andersch for help with analysing flow cytometry based CRISPR screens. We want to thank the Core Facility Flow Cytometry of the Medical University of Vienna for access to flow cytometry instruments and assistance with flow cytometric cell sorting; the CeMM Biomedical Sequencing Facility for NGS sample processing, sequencing and data curation; and Thomas Hannich and the Proteomics Facility of the CeMM Molecular Discovery Platform for access to instruments. We thank Diane Cassidy from CeTPD for the assistance with the maintenance of cell lines and support with DNA prep and sequencing. We also want to acknowledge the MRC PPU DNA sequencing and services facility for additional cloning and sequencing support. We thank Nicholas Larsen at Foghorn Therapeutics for sharing plasmids and Kristin Riching at Promega for the gift of the bromodomain conformational sensor plasmid.

## Contributions

O.H., M.H. and A.D.C. contributed equally and will be putting their name first on the citation in their CVs. O.H., M.H., A.D.C, G.E.W. and A.C. conceived and planned this project. O.H., M.H. and A.D.C. designed and conducted experiments with help from K.I., T.I. and A.R. O.H., M.H., A.D.C., G.E.W. and A.C. analysed and interpreted original data. K.I., T.I., R.C., C.M. and A.T. designed and synthesized compounds. H.I. and C.S. analysed FACS-based CRISPR screens and K.H. and M.W. performed and analysed viability-based CRISPR screens with input from M.K. and I.D., A.R. performed and analysed quantitative expression proteomics. A.D.C. performed Cryo-EM imaging, data processing and 3D reconstruction with help from R.S. and M.A.N.; M.A.N., A.C-S., C.C., C.M. and A.T. established critical reagents and methodology. O.H., M.H., A.D.C., G.E.W. and A.C. co-wrote the manuscript with input from all co-authors.

## Data availability

Source data for Fig. 1c, Fig. 2b, c, Fig. 5d, Extended Data Fig. 2a and Extended Data Fig. 6f are included in the Supplementary information files of the manuscript. Cryo-EM density maps are deposited in the EMDB with the accession code EMD-17172 and will be released upon publication. The atomic model is deposited under Protein Data Bank ID 8OV6. Raw micrographs and particle stacks will be available in the EMPIAR-database. Quantitative proteomics data have been deposited to the ProteomeXchange Consortium PRIDE repository^50^ with the accession ID PXD040570.

## Extended Data

**Extended Data Fig. 1:**
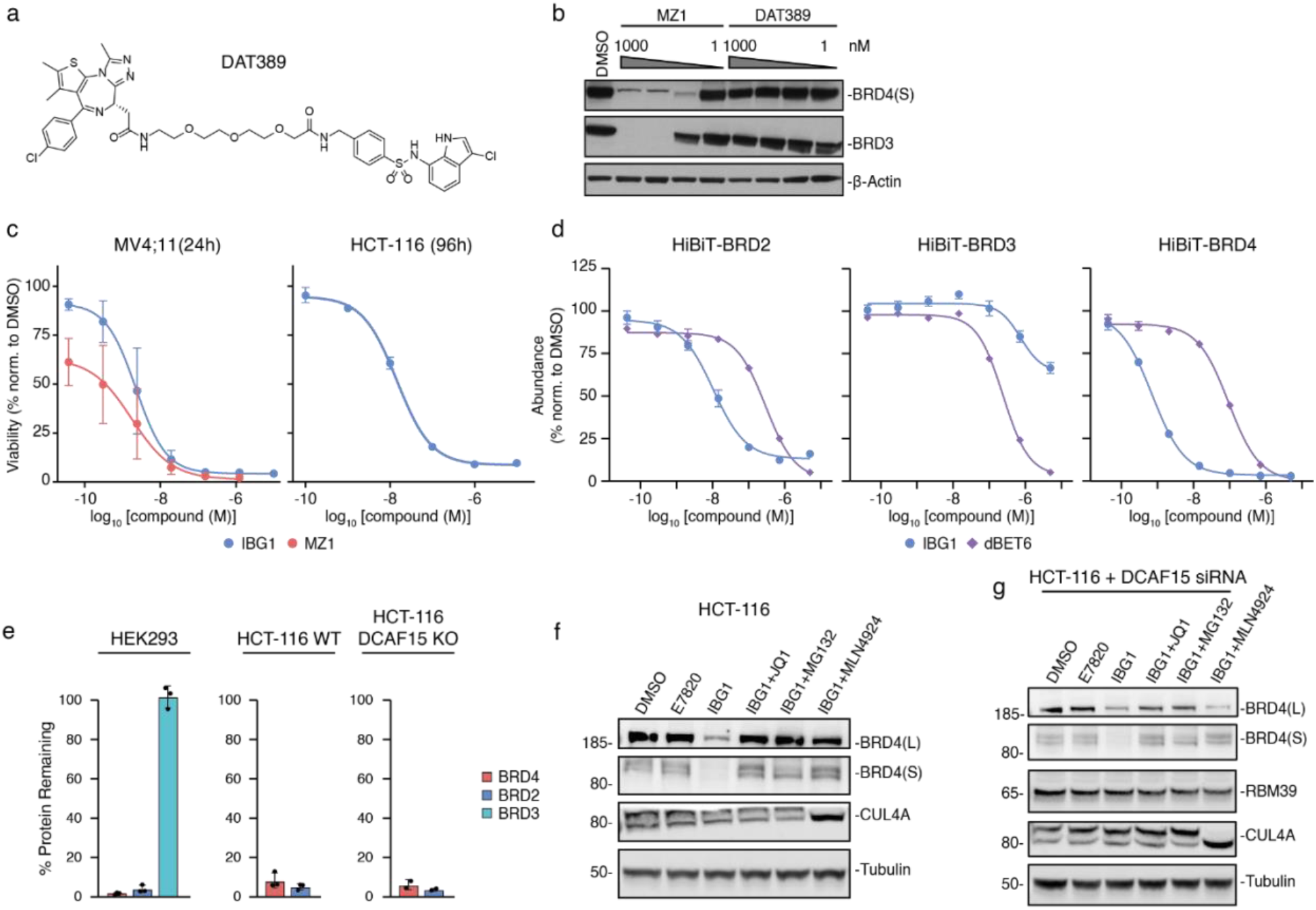
IBG1 degrades BRD2 and BRD4 independent of DCAF15. **a**, **b**, Structure (**a**) and BET protein degradation (**b**) of sulfonamide-based PROTAC DAT389. HeLa cells were treated with increasing concentrations of MZ1 or DAT389 for 16 hours and BET protein levels were analysed by immunoblot. **c**, Cytotoxicity of IBG1 and VHL-based PROTAC MZ1. MV4;11 and HCT-116 cells were treated with increasing concentrations of compounds for 24 or 96 hours, respectively, and cell viability was assessed via CellTiterGlo assay. Dose-response curves were fitted using non-linear regression. n = 2 biological repeats, mean +/- s.d. **d**, End-point HiBiT protein degradation. BRD2, BRD3 or BRD4 HiBiT knock-in HEK293 cells were treated with the indicated compounds for 5 hours and levels of HiBiT-tagged proteins were quantified via the HiBiT lytic detection system. Dose-response curves were fitted using non-linear regression. n = 3 independent repeats, mean +/- s.d. **e**, Degradation activities of IBG1. BET protein levels were quantified (from n = 3 independent experiments) based on immunoblotting after compound treatment in HEK293, HCT-116 WT and DCAF15 KO cells. Source data, Supplementary Fig. 1a. **f**, **g**, In-cell mechanistic evaluation of IBG1. HCT-116 WT (**f**) or DCAF15 knockdown (**g**) cells were treated for 2 hours with E7820 (1 µM) or IBG1 (10 nM) alone, or after 1 hour pre-treatment with JQ1 (10 µM), MG132 (50 µM) or MLN4924 (3 µM). Western blot representative of 3 (**f**) or 2 (**g**) independent experiments.

**Extended Data Fig. 2:**
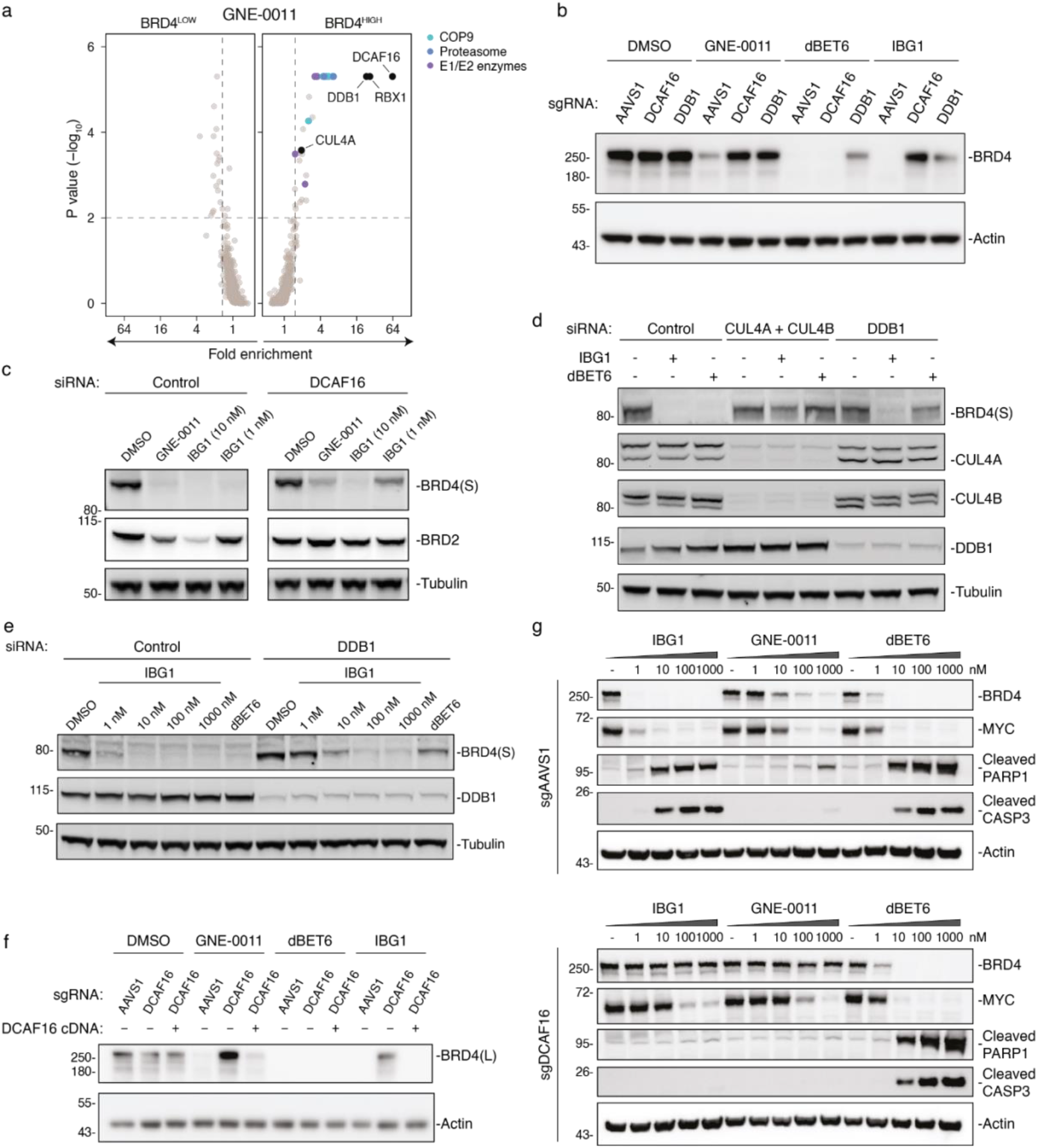
IBG1 degrades BRD2/4 via CRL4DCAF16. **a**, BRD4 stability CRISPR screen. KBM7 iCas9 BRD4 dual fluorescence reporter cells expressing a CRL-focused sgRNA library were treated with GNE-0011 (1 µM) for 6 hours before flow cytometric cell sorting into BRD4^LOW^, BRD4^MID^ and BRD4^HIGH^ fractions as in Fig. 2b. 20S proteasome subunits (blue), COP9 signalosome subunits (cyan) and E1 or E2 ubiquitin enzymes (purple) inside the scoring window (p-value< 0.01, fold-change > 1.5; dashed lines) are highlighted. **b**-**f**, Immunoblot-based CRISPR/Cas9 screen validation. **b**, KBM7 iCas9 cells were lentivirally transduced with sgRNAs targeting AAVS1, DCAF16 or DDB1 and 3 days after Cas9 induction, cells were treated with GNE-0011 (1 µM), dBET6 (10 nM) or IBG1 (1 nM) for 6 hours and BRD4 levels were analysed via immunoblot. Data is representative for n = 2 independent experiments. **c**-**e**, HCT-116 cells were transfected with siRNA pools targeting the indicated genes and treated with DMSO, IBG1, GNE-0011 or dBET6 for 2 h at the indicated concentrations and BET protein levels were analysed via immunoblotting. Data are representative of n = 2 independent experiments**. f**, DCAF16 knockout/rescue. KBM7 iCas9 cells were lentivirally transduced with the indicated DCAF16-targeting or AAVS1 control sgRNAs, as well as a DCAF16 cDNA in which the sgRNA target sites were removed by synonymous mutations. After knockout of endogenous DCAF16 and compound treatment for 6 hours as above, BRD4 expression levels were assessed via immunoblotting. **g**, Induction of apoptosis. KBM7 iCas9 WT or DCAF16 knockout cells were treated with increasing concentrations of IBG1, GNE-0011 or dBET6 for 16 h and levels of BRD4, MYC, cleaved PARP1 and cleaved caspase 3 were analysed via immunoblotting as in Fig. 2g. Data are representative of n = 3 independent experiments.

**Extended Data Fig. 3:**
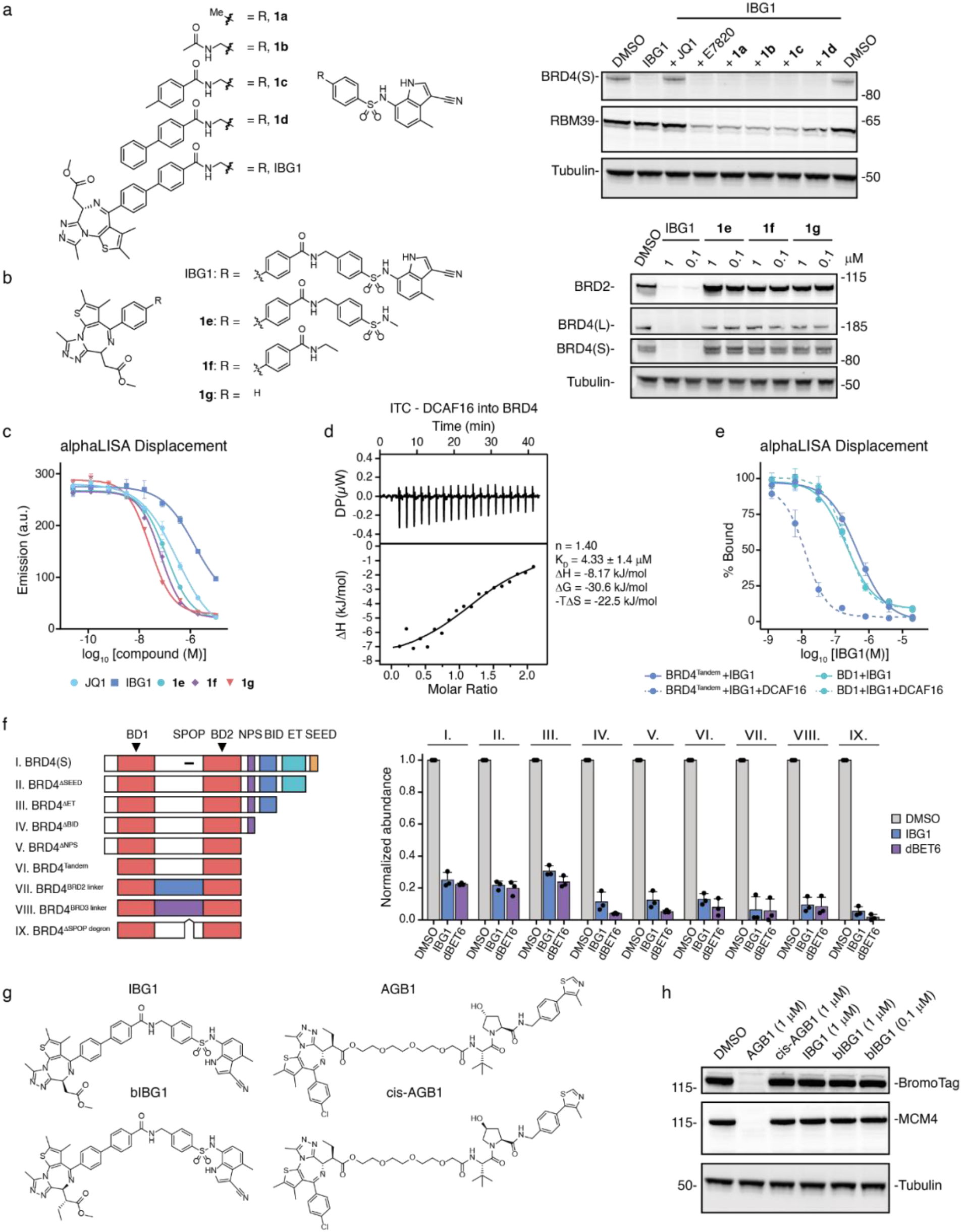
Mechanistic evaluation of IBG1 mode of action. **a**, Competitive degradation assay. HCT-116 cells were pre-treated for 1 hour with 10 µM of sulfonamide-containing truncations of IBG1 (compounds **1a**-**d**), followed by 2-hour treatment with IBG1 (10 nM) and immunoblot analysis. Data is representative of n = 2 independent experiments. **b**, Degradation activities of JQ1-containing truncations of IBG1. HCT-116 were cells treated with indicated concentrations of JQ1-containing truncations of IBG1 (compounds 1**e**-**g**) for 6 hours and analysed by immunoblotting. Data is representative of n = 2 independent experiments. **c**, alphaLISA displacement assay. His-BRD4^BD2^ preincubated with a biotinylated JQ1 probe was titrated against increasing concentrations of IBG1 or truncated compounds **1e**-**g**. n = 3 technical replicates, mean +/- s.d. **d**, Isothermal titration calorimetry measurement of DCAF16:DDB1ΔBPB:DDA1 binding to BRD4^Tandem^. **e**, alphaLISA displacement assay. His-BRD4^Tandem^ or His-BRD4^BD1^ were preincubated with a biotinylated JQ1 probe and titrated against increasing concentrations of IBG1 in the presence or absence of DCAF16. n = 2 independent experiments each with 3 technical replicates, mean +/- s.d. **f**, Protein stability reporter assay. WT or truncated forms of BRD4 fused to mTagBFP (left) were stably expressed in KBM7 cells and protein stability after 6-hour treatment with DMSO, IBG1 (1 nM) or dBET6 (10 nM) was quantified via flow cytometric evaluation of the mTagBFP/mCherry ratio (right). BD, bromodomain; NPS, N-terminal phosphorylation sites; BID, basic residue-enriched interaction domain; ET, extraterminal domain; SEED, Serine/Glutamic acid/Aspartic acid-rich region. n = 3 independent experiments, mean +/- s.d. **g**, **h**, BromoTag degradation. HEK293 cells stably expressing BromoTag-MCM4 were treated for 5 hours with DMSO, BromoTag degrader AGB1 and non-degrader cis-AGB1, IBG1, or ‘bumped’ IBG1 analogue bIBG1 (**g**) and BromoTag-MCM4 levels were analysed by immunoblotting (**h**). Data representative of n = 2 independent experiments.

**Extended Data Fig. 4:**
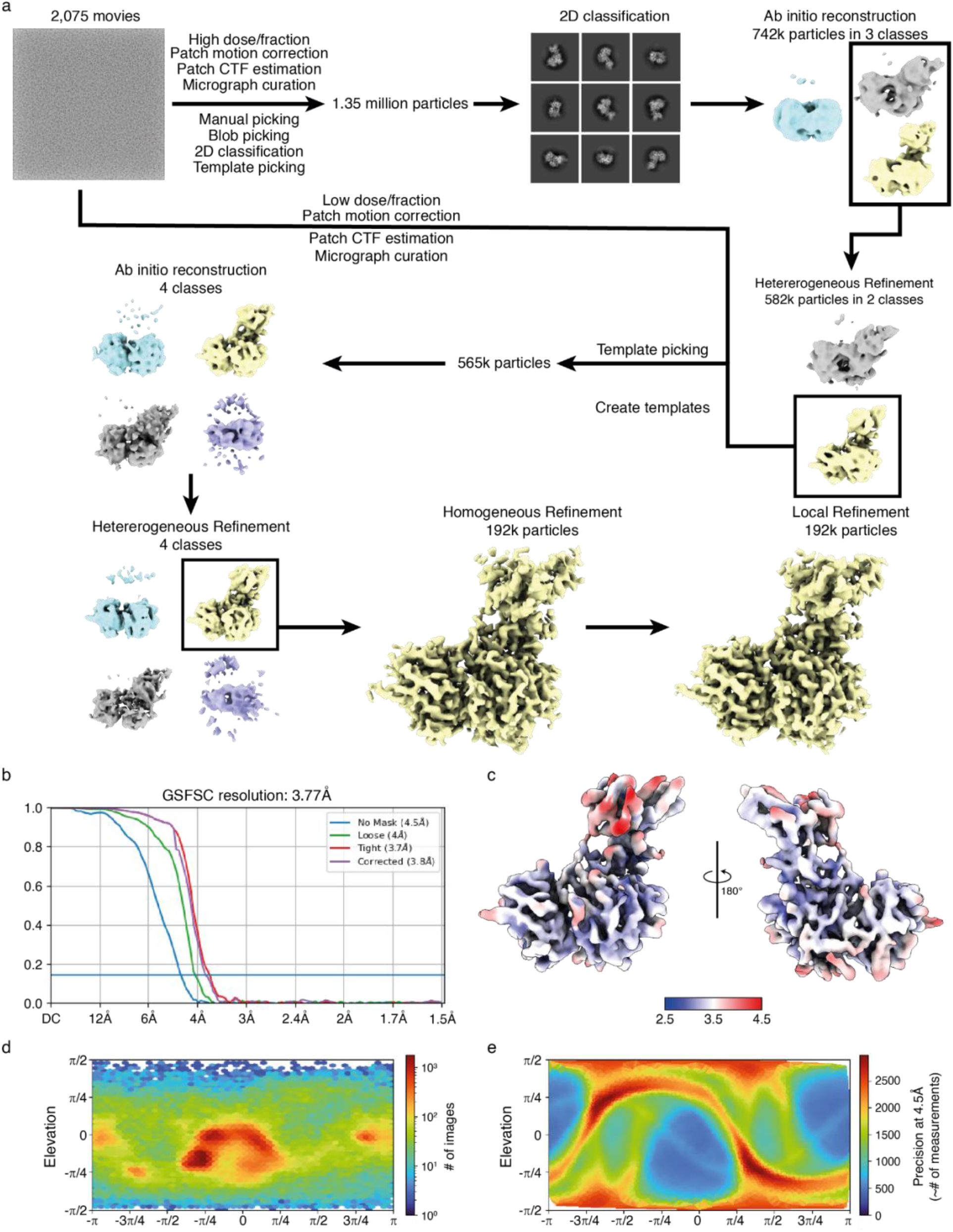
Cryo-EM data processing. **a**, Workflow for Cryo-EM data processing. **b**, Gold-standard Fourier shell correlation at a cut-off of 0.143. **c**, Local resolution estimation on the unsharpened map. **d**, **e**, Angular distribution plot (**d**) and posterior position directional distribution plot (**e**) for the final local refinement.

**Extended Data Fig. 5:**
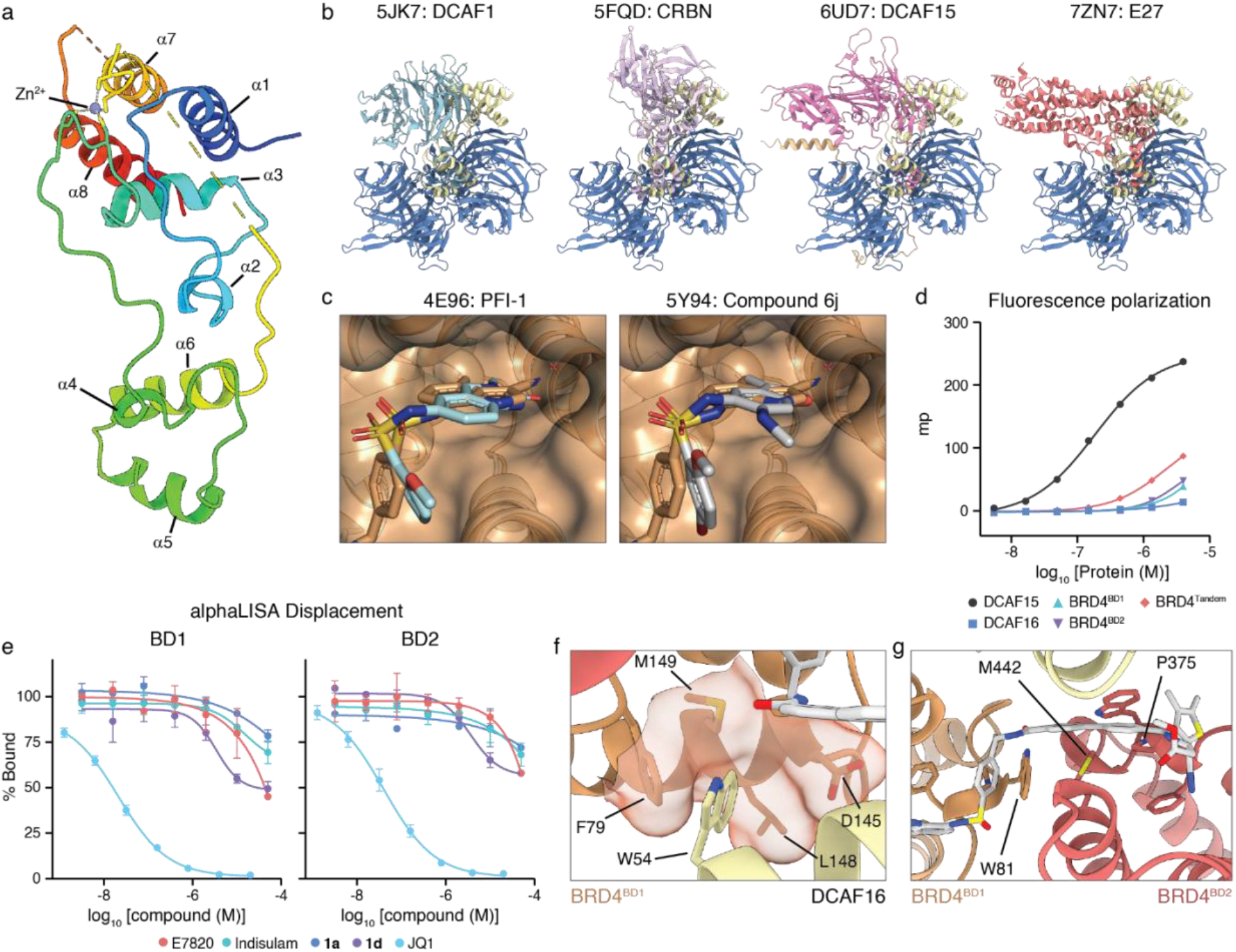
Structure-based characterization of the ternary BRD4:IBG1:DCAF16 complex. **a**, Structure of DCAF16 coloured rainbow from N-to C-terminus. **b**, Superposition of DCAF16 (yellow) vs. known substrate receptors bound to DDB1 (blue). **c**, Comparison of binding mode in the acetyl-lysine pocket of BRD4^BD1^ (orange surface and cartoon) between IBG1 (orange) and known sulfonamide BET inhibitors PFI-1 (left, light blue) and compound 6j (right, grey). The cyano group of IBG1 overlays close to a conserved water molecule found in both crystal structures and many other published BD1 structures. **d**, Fluorescence polarization binary binding assay. Indicated proteins were titrated into 20 nM FITC-sulfonamide probe (from Fig. 2g). n = 3 technical replicates, mean +/- s.d. **e**, alphaLISA displacement assay. Competition of a biotinylated-JQ1 probe following titration of compounds **1a**, **1d**, E7820, Indisulam, or JQ1 into His-BRD4^BD1^ (left) or His-BRD4^BD2^ (right). n = 2 technical replicates, mean +/- s.d. **f**, Detailed view of DCAF16:BD1 interface. Residue W54 of DCAF16 bound to a hydrophobic pocket on the surface of BD1. **g**, Detail view of BD1:BD2 interface. Residue M442 of BD2 is sandwiched between residues W81 and P375 of the BD1 and BD2 WPF shelves, respectively, as well as the linker of IBG1.

**Extended Data Fig. 6:**
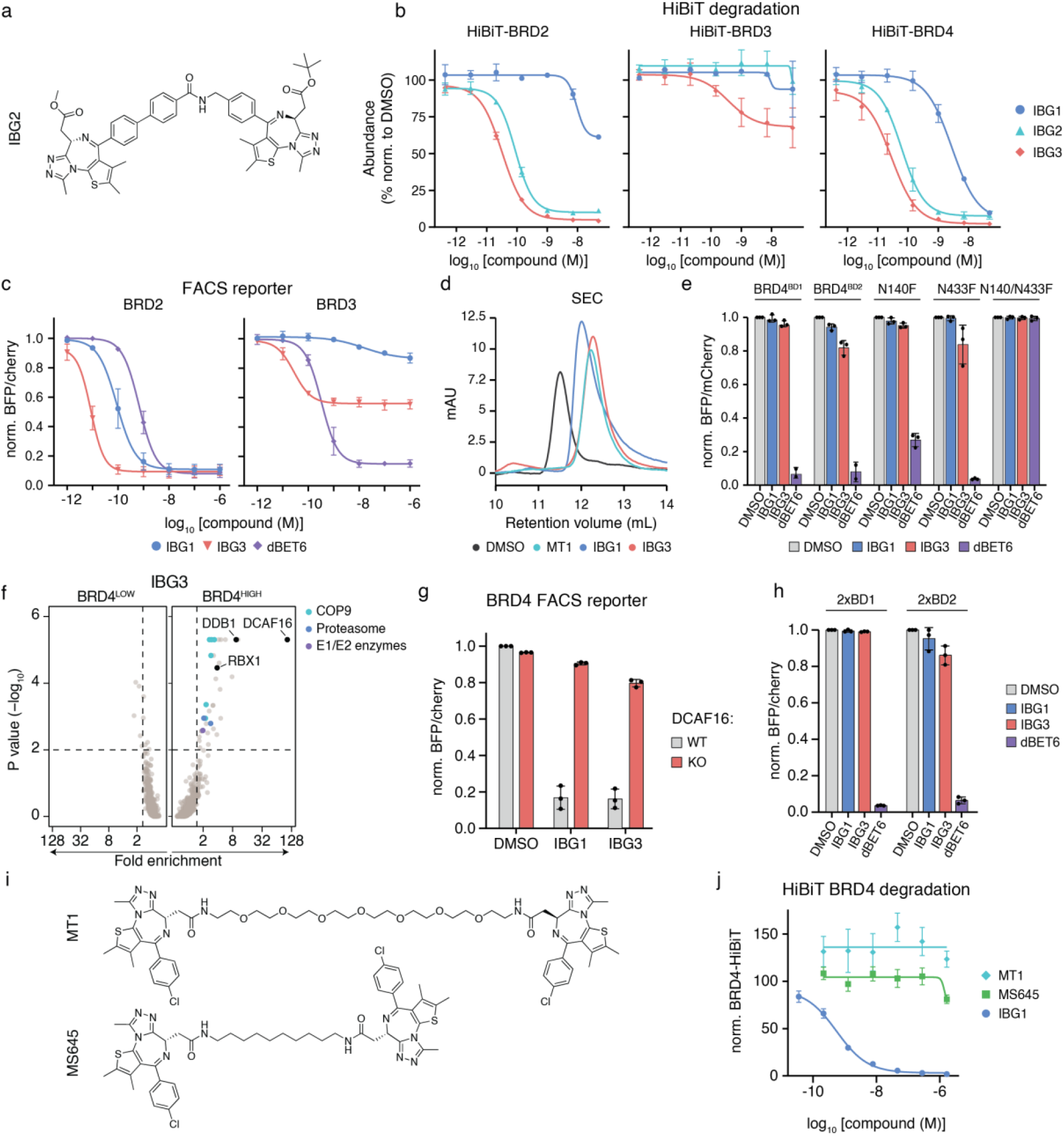
Rational design of improved intramolecular bivalent glue BET degraders. **a**, Structure of double JQ1 containing intramolecular bivalent glue degrader IBG2. **b**, HiBiT degradation assay. HEK293 HiBiT knock-in cells were treated with the indicated compounds for 5 h and levels of BRD2-, BRD3- and BRD4-HiBiT proteins were quantified via luminescence as in Extended Data Fig. 1b. Data shown are from n = 3 independent experiments, mean +/- s.d. **c**, BET protein specificity. KBM7 cells expressing BRD2^Tandem^ or BRD3^Tandem^ dual fluorescence reporters were treated with increasing concentrations of IBG1, IBG3 or dBET6 for 6 hours and BET protein levels were quantified via flow cytometry. **d**, Size exclusion chromatograms of BRD4^Tandem^ incubated with DMSO, MT1, IBG1 or IBG3. Data for DMSO and IBG1 as in Fig. 4c. **e**, Bromodomain tandem selectivity. KBM7 cells expressing isolated BRD4 bromodomains or mutated BRD4^Tandem^ constructs were treated with IBG1 (1 nM), IBG3 (0.1 nM) or dBET6 (10 nM) for 6h and protein levels were evaluated via flow cytometry. **f**, BRD4 stability CRISPR screen. KBM7 iCas9 BRD4 dual fluorescence reporter cells expressing a CRL-focused sgRNA library were treated with IBG3 (0.1 nM) for 6 hours before flow cytometric cell sorting as in Fig. 2b. 20S proteasome subunits (blue), COP9 signalosome subunits (cyan) and E1 or E2 ubiquitin enzymes (purple) inside the scoring window (p-value < 0.01, fold-change > 1.5; dashed lines) are highlighted. **g**, DCAF16 dependency. BRD4(S) dual fluorescence reporter KBM7 iCas9 cells were lentivirally transduced with a DCAF16-targeting sgRNA and 3 days post Cas9 induction cells were treated with DMSO, IBG1 (1nM) or IBG3 (0.1 nM) for 6 hours before FACS-based quantification of BRD4 levels. **h**, Bromodomain arrangement. KBM7 cells expressing dual fluorescence reporters harbouring tandems of either BD1 or BD2 of BRD4 were treated with DMSO, IBG1 (1 nM), IBG3 (0.1 nM) or dBET6 (10 nM) for 6 hours and analysed by flow cytometry. **i**, **j**, Structures (**i**) and HiBiT-BRD4 degradation activity (**j**) of bivalent BET inhibitors MT1 and MS645 after treatment for 24 hours. Data for **c**, **e**, **g**, **h** are from n = 3 independent experiments, mean +/- s.d. Data in j is for n = 2 independent experiments, mean +/- s.d.

**Extended Data Fig. 7:**
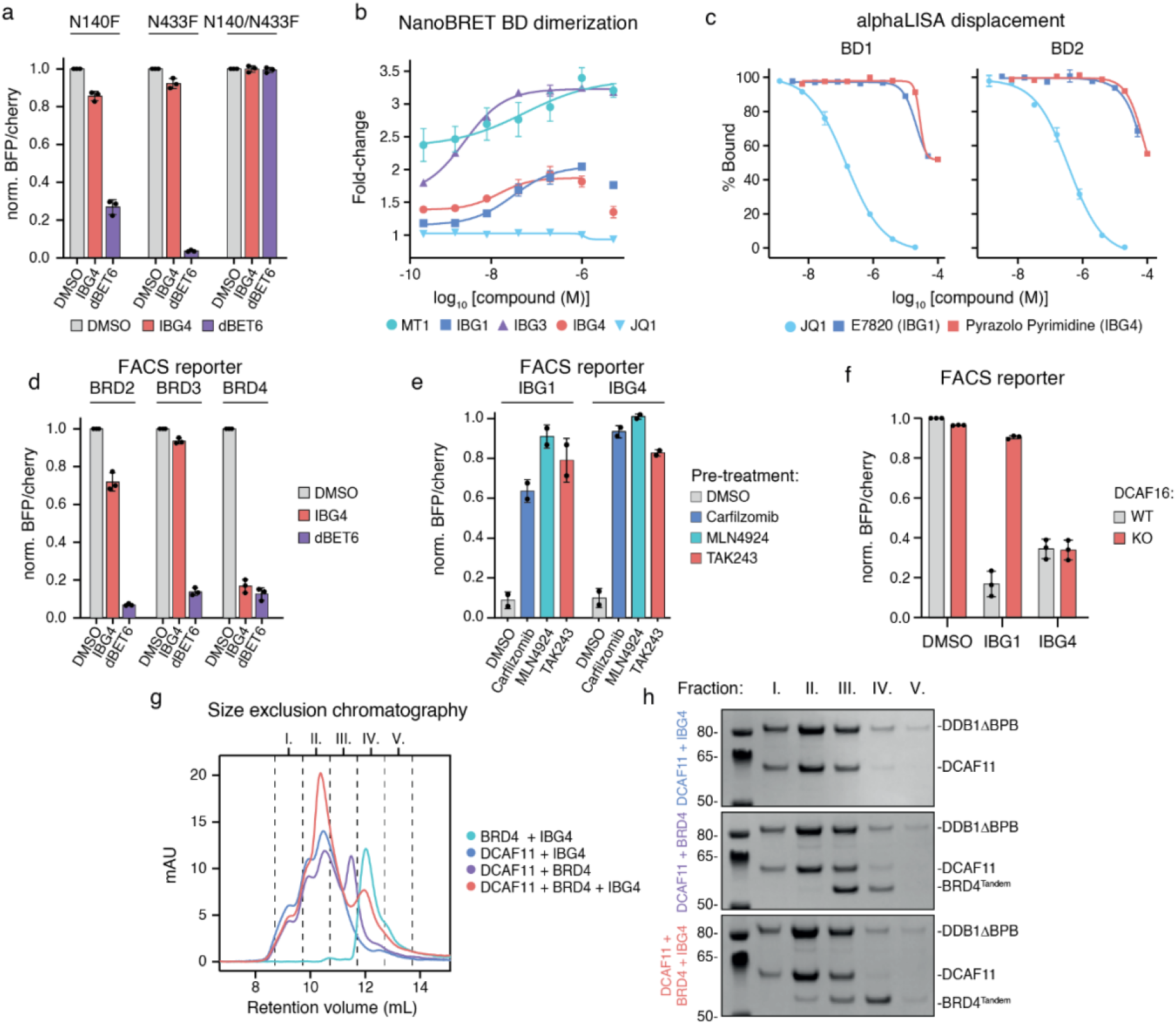
IBG4 is a DCAF11-dependent intramolecular bivalent glue degrader. **a**, Bromodomain tandem specificity. KBM7 cells expressing bromodomain mutant BRD4^Tandem^ dual fluorescence reporters were treated with DMSO, IBG4 (100 nM) or dBET6 (10 nM) for 6 hours and analysed by flow cytometry. **b**, NanoBRET bromodomain dimerization assay. Indicated compounds were titrated into transiently expressed BRD4^Nluc-Tandem-HaloTag^ in HEK293 cells. **c**, alphaLISA displacement assay. Increasing concentrations of JQ1, E7820 or the pyrazolo pyrimidine warhead of IBG4 were titrated against His-tagged BRD4 bromodomains and biotinylated JQ1 probe. n = 3 technical replicates, mean +/- s.d. **d**, BET protein selectivity. Bromodomain tandem BRD2, BRD3 or BRD4 dual fluorescence reporter KBM7 cells were treated with DMSO, IBG4 (100 nM) or dBET6 (10 nM) for 6 hours and analysed by flow cytometry. **e**, Mechanistic FACS reporter assay. KBM7 BRD4 dual fluorescence reporter cells were co-treated with IBG1 (1 nM) or IBG4 (100 nM) and Carfilzomib (1 µM), MLN4924 (1 µM) or TAK243 (0.5 µM) for 6 hours and BRD4 levels were analysed via flow cytometry. **f**, DCAF16-independence of c IBG4. KBM7 iCas9 WT or DCAF16 knockout cells expressing BRD4(S) dual fluorescence reporter were treated with DMSO, IBG1 (1 nM) or IBG4 (100 nM) for 6h and BRD4 degradation was assessed via flow cytometry. **g**, **h**, Size exclusion chromatograms of different combinations of DCAF11, BRD4^Tandem^ and IBG4 (**g**) and corresponding peak fractions run on SDS-PAGE (**h**). Data for **a**, **d**, **f**, n = 3 independent experiments, data for **b**, **e**, n = 2 independent experiments, mean +/- s.d.

**Extended Data Table 1:**
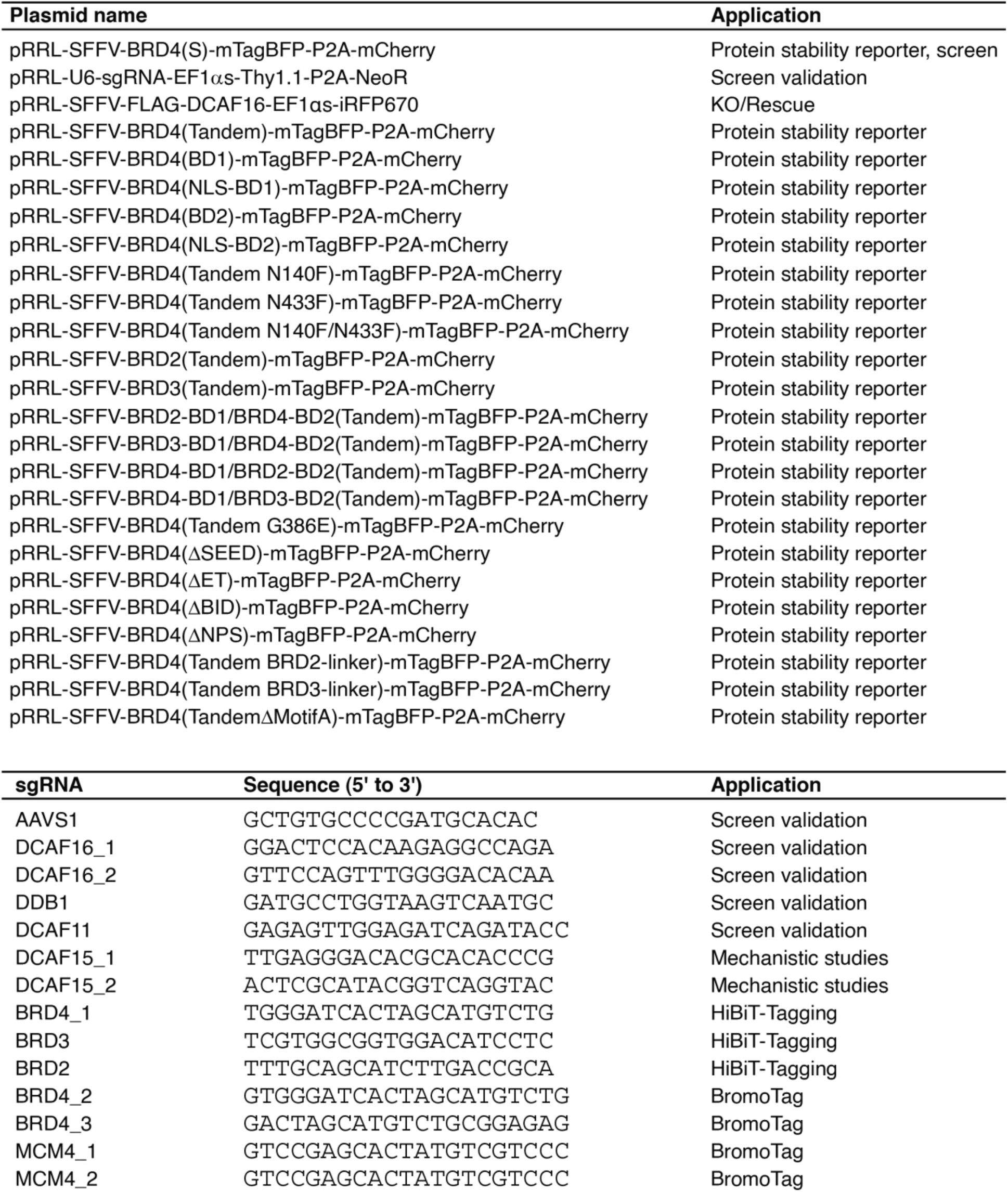
Plasmids and sgRNAs used in this study.

**Extended Data Table 2:** Summary of cryo-EM data collection conditions and image processing.

|  |  |
| --- | --- |
| **Data collection** |  |
| Magnification | 190,000 |
| Voltage (kV) | 200 |
| Detector | Falcon 4i |
| Electron total dose (e-/Å<sup>2</sup>) | 12.7 |
| Data Format | EER |
| Defocus range (μm) | -(1.7-3.2) |
| Pixel size (Å) | 0.74 |
| Exposure time (Sec) | 2 |
| Total fractions | 18 |
| Dose per fraction (e-/Å<sup>2</sup>) | 0.69 |
| **Reconstruction** |  |
| Symmetry imposed | C1 |
| Initial particle images (no.) | 1,466,607 |
| Final particle images (no.) | 530,879 |
| Map resolution (Å) | 4 |
| FSC threshold | 0.143 |
| Map resolution range (Å) | 4-8 |
| Sharpening B-factor (Å<sup>2</sup>) | -210.5 |

## Methods

### 1. Chemistry

Chemicals that are commercially available were purchased from Apollo Scientific, Sigma-Aldrich, Fluorochem, CombiBlocks, TCI, and Enamine and were used without further purification. Liquid chromatography–mass spectrometry (LC-MS) was carried out on a Shimadzu HPLC/MS 2020 equipped with a Hypersil Gold column (1.9 μm, 50 × 2.1 mm^2^), a photodiode array detector, and an electrospray ionization (ESI) detector. The samples were eluted with a 3 min gradient of 5–95% acetonitrile in water containing 0.1% formic acid at a flow rate of 0.8 mL/min. Flash column chromatography was performed on a Teledyne ISCO Combiflash Companion installed with disposable normal phase RediSep Rf columns (230–400 mesh, 40–63 mm; SiliCycle). Preparative HPLC purification was performed on a Gilson preparative HPLC system equipped with a Waters X-Select C18 column (100 mm × 19 mm and 5 μm particle size) using a gradient from 5 to 95% of acetonitrile in water containing 0.1% formic acid over 10 min at a flow rate of 25 mL/min. Compound characterization using NMR was performed either on a Bruker 500 Ultra shield or on a Bruker Ascend 400 spectrometer. The ^1^H NMR and ^13^C NMR reference solvents used are CDCl3*-d1* (δH = 7.26 ppm/δC = 77.16 ppm), CD3OD (δH = 3.34 ppm/δC = 49.86 ppm), DMSO*-d6* (δH = 2.50 ppm/δC = 39.52 ppm), or acetone*-d6* (δH = 2.05 ppm/δC = 29.84 ppm). Signal patterns are described as singlet (s), doublet (d), triplet (t), quartet (q), multiplet (m), broad singlet (bs), or a combination of the listed splitting patterns. The coupling constants (*J*) are measured in hertz (Hz).

#### Synthesis of IBG1

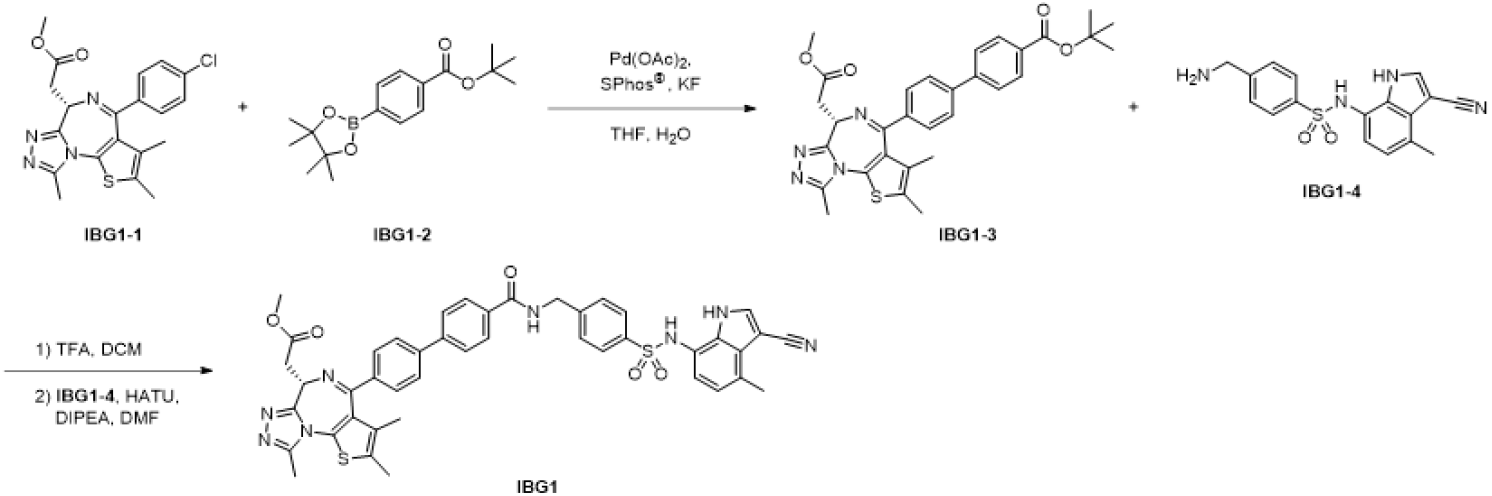

##### *tert*-Butyl (S)-4’-(6-(2-methoxy-2-oxoethyl)-2,3,9-trimethyl-6H-thieno[3,2-f][1,2,4]triazolo[4,3-a][1,4]diazepin-4-yl)-[1,1’-biphenyl]-4-carboxylate (IBG1-3)

A mixture of methyl (R)-2-(4-(4-chlorophenyl)-2,3,9-trimethyl-6H-thieno[3,2-f][1,2,4]triazolo[4,3-a]azepin-6-yl)acetate (**IBG1-1**) (126 mg, 0.30 mmol), 4-(*tert*-utoxycarbonyl)phenylboronic acid pinacol ester (**IBG1-2**) (101 mg, 0.33 mmol, 1.1 eq.), potassium fluoride (52.8 mg, 0.91 mmol, 3.0 eq.), SPhos^®^ (12.4 mg, 0.03 mmol, 0.1 eq.), palladium(II) acetate (6.8 mg, 0.03 mmol, 0.1 eq.), and water (20 µL) in THF (1.0 mL) was purged with nitrogen atmosphere then the mixture was stirred at reflux temperature overnight. The resulted mixture was diluted with excess amount of EtOAc, filtered through a celite pad, washed with EtOAc, concentrated in vacuo, and then purified by silica gel column chromatography (DCM-MeOH) to afford **IBG1-3** (193 mg, quantitative yield.). ^1^H NMR (500 MHz, CDCl_3_) δ 8.05 (d, *J* = 8.4 Hz, 2H), 7.63-7.59 (m, 4H), 7.56-7.54 (m, 2H), 4.66 (dd, *J* = 7.8, 6.3 Hz, 1H), 3.79 (s, 3H), 3.72-3.62 (m, 2H), 2.69 (s, 3H), 2.43 (s, 3H), 1.75 (s, 3H), 1.61 (s, 9H). ^13^C NMR (101 MHz, CD_3_OD) δ 174.08, 167.99, 167.91, 157.75, 153.06, 146.32, 144.46, 139.95, 134.31, 134.10, 133.41, 133.22, 133.14, 131.89, 131.33, 129.23, 128.93, 83.33, 55.82, 53.31, 38.11, 29.34, 15.29, 13.83, 12.48. LC-MS, ESI^+^, *m/z* 557.4 [M+H]^+^.

##### Methyl (S)-2-(4-(4’-((4-(*N*-(3-cyano-4-methyl-1H-indol-7-yl)sulfamoyl)benzyl)carbamoyl)-[1,1’-biphenyl]-4-yl)-2,3,9-trimethyl-6H-thieno[3,2-f][1,2,4]triazolo[4,3-a][1,4]diazepin-6-yl)acetate (IBG1)

To a solution of **IBG1-3** (57.0 mg, 90 µmol) in DCM (0.5 mL) was added trifluoroacetic acid (0.5 mL) and then the mixture was stirred at room temperature for 4 hours. The resulted mixture was concentrated in vacuo, and then toluene was added thereto. After concentrated in vacuo again, the obtained crude compound, 4-(aminomethyl)-*N*-(3-cyano-4-methyl-1H-indol-7-yl)benzenesulfonamide (**IBG1-4**)^6^ (30.8 mg, 90.5 µmol, 1.0 eq.), and *N,N*-diisopropylethylamine (78.8 µL, 453 µmol, 5.0 eq.) were mixed in *N*,*N*-dimethylformamide (1.0 mL) and then HATU (61.9 mg, 163 µmol, 1.8 eq.) was added to it. After the mixture was stirred at room temperature overnight, the resulted mixture was purified with preparative HPLC (ODS, H_2_O-MeCN with 0.1% HCOOH) to afford **IBG1** (26.2 mg, 35% yield). ^1^H NMR (500 MHz, DMSO-*d6*) δ 11.86 (d, *J* = 2.7 Hz, 1H), 9.90 (s, 1H), 9.14 (t, J = 6.0 Hz, 1H), 8.15 (d, *J* = 3.1 Hz, 1H), 8.00 (d, J = 8.4 Hz, 2H), 7.83-7.79 (m, 4H), 7.69 (d, *J* = 8.4 Hz, 2H), 7.53 (d, *J* = 8.3 Hz, 2H), 7.45 (d, *J* = 8.3 Hz, 2H), 6.77 (d, *J* = 7.8 Hz, 1H), 6.62 (d, 7.8 Hz, 1H), 4.55-4.52 (m, 3H), 3.69 (s, 3H), 3.54-3.43 (m, 2H), 2.62 (s, 3H), 2.56 (s, 3H), 2.43 (s, 3H), 1.69 (s, 3H). ^13^C NMR (126 MHz, DMSO-*d6*) δ 171.07, 165.87, 163.87, 154.72, 149.89, 144.91, 141.80, 140.90, 137.67, 137.33, 135.14, 133.26, 132.03, 130.71, 130.36, 130.04, 129.84, 129.00, 127.99, 127.54, 127.26, 127.02, 126.76, 126.63, 126.44, 122.40, 120.55, 118.01, 117.29, 84.30, 53.42, 51.51, 42.24, 36.28, 17.56, 14.04, 12.65, 11.21. HRMS, ESI^+^, *m/z* calcd for C44H39N8O5S2 [M+H]^+^ 823.2485; found 823.2505.

#### Synthesis of compound 1a

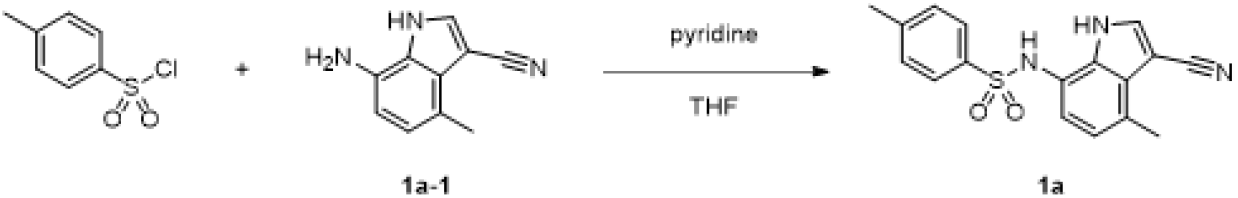

##### *N*-(3-Cyano-4-methyl-1H-indol-7-yl)-4-methylbenzenesulfonamide (1a)

To a solution of 7-amino-4-methyl-1H-indole-3-carbonitrile (**1a-1**) (30 mg, 0.18 mmol) and pyridine (42.5 µL, 0.53 mmol, 3.0 eq.) in tetrahydrofuran (1.0 mL) was added 4-toluenesulfonyl chloride (50.1 mg, 0.26 mmol, 1.5 eq.) and then the mixture was stirred at room temperature overnight. The resulted mixture was diluted with DMSO (1.0 mL), filtered through a membrane filter (0.43 µm), and then purified by preparative HPLC (ODS, H_2_O-MeCN with 0.1% HCOOH) to afford **1a** (39.3 mg, 0.117 mmol in 71% yield). ^1^H NMR (400 MHz, DMSO-*d6*) δ 11.85 (bs, 1H), 9.82 (bs, 1H), 8.15 (s, 1H), 7.58 (d, *J* = 8.3 Hz, 2H), 7.31 (d, *J* = 8.1 Hz, 2H), 6.77 (d, J = 7.7 Hz, 1H), 6.60 (d, *J* = 7.7 Hz, 1H), 2.56 (s, 3H), 2.34 (s, 3H). ^13^C NMR (101 MHz, DMSO-*d4*) δ 144.87, 138.10, 136.81, 132.13, 131.18, 128.89, 128.59, 128.11, 124.09, 122.37, 119.79, 119.01, 85.98, 22.60, 19.26. LC-MS, ESI^+^, *m/z* 323.8 [M+H]^+^.

#### Synthesis of compound 1b

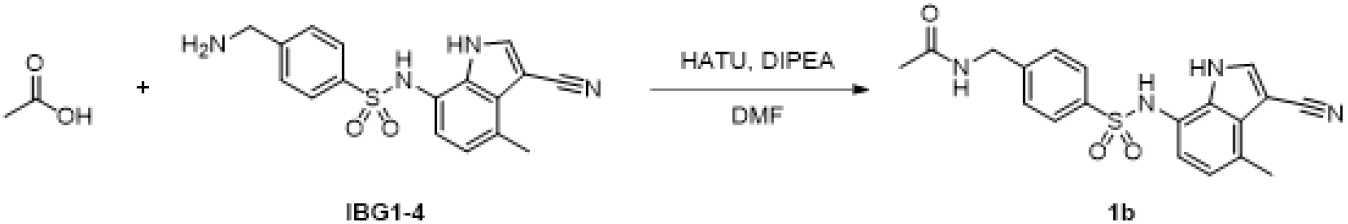

##### *N*-(4-(*N*-(3-Cyano-4-methyl-1H-indol-7-yl)sulfamoyl)benzyl)acetamide (1b)

To a solution of 4-(aminomethyl)-*N*-(3-cyano-4-methyl-1H-indol-7-yl)benzenesulfonamide (**IBG1-4**) (20 mg, 58.8 µmol, 1.0 eq.), acetic acid (5.0 µL, 456 µmol, 5.0 eq.), and *N,N*-diisopropylethylamine (51.2 µL, 294 µmol, 5.0 eq.) in *N,N*-dimethylformamide (0.5 mL) was added HATU (40.2 mg, 106 µmol, 1.8 eq.) and then the mixture was stirred at room temperature for 6 hours. The resulted mixture was diluted with 0.5 mL of DMSO, filtered, and purified with preparative HPLC (ODS, H_2_O-MeCN with 0.1% HCOOH) to afford **1b** (3.4 mg, 15% yield). ^1^H NMR (400 MHz, acetone-*d6*) δ 8.05 (s, 1H), 7.62 (d, *J* = 8.3 Hz, 2H), 7.39 (d, *J* = 8.3 Hz, 2H), 6.78 (d, *J* = 7.7 Hz, 1H), 6.63 (d, *J* = 7.7 Hz, 1H), 4.41 (s, 2H), 2.64 (s, 3H), 1.95 (s, 3H). ^13^C NMR (101 MHz, acetone-*d6*) δ 171.07, 146.14, 138.80, 135.07, 132.41, 129.81, 128.64, 128.35, 127.96, 123.61, 121.37, 120.96, 100.21, 86.66, 43.08, 22.71, 18.20. LC-MS, ESI^+^, *m/z* 382.9 [M+H]^+^.

#### Synthesis of compound 1c

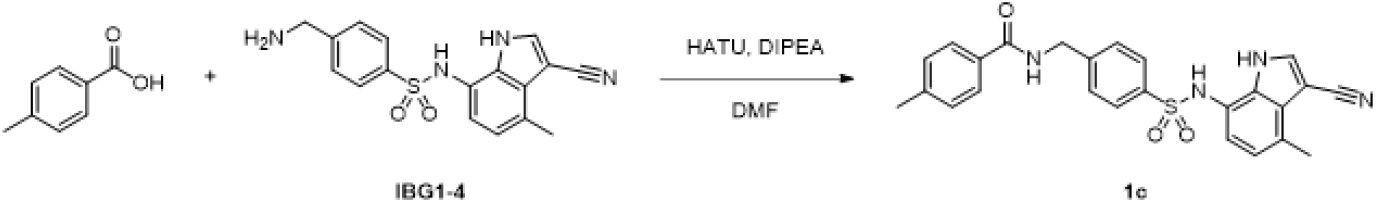

##### *N*-(4-(*N*-(3-Cyano-4-methyl-1H-indol-7-yl)sulfamoyl)benzyl)-4-methylbenzamide (1c)

To a solution of 4-(aminomethyl)-*N*-(3-cyano-4-methyl-1H-indol-7-yl)benzenesulfonamide (**IBG1-4**) (30.0 mg, 79.6 µmol), *p*-toluic acid (13.0 mg, 95.5 µmol, 5.0 eq.), and *N,N*-diisopropylethylamine (69.3 µL, 398 µmol, 5.0 eq.) in *N,N*-dimethylformamide (0.5 mL) was added HATU (45.4 mg, 119 µmol, 1.5 eq.) and then the mixture was stirred at room temperature for 5 hours. The resulted mixture was purified with silica gel column chromatography (ODS, H_2_O-MeCN with 0.1% HCOOH). The obtained compound was triturated with DCM. The solid was collected by filtration, washed with DCM, and then dried in vacuo to afford **1c** (4.8 mg, 13% yield). ^1^H NMR (500 MHz, acetone-*d6*) δ 11.09 (bs, 1H), 8.87 (bs, 1H), 8.28 (bs, 1H), 8.09 (s, 1H), 7.83 (d, *J* = 8.2 Hz, 2H), 7.65 (d, *J* = 8.4 Hz, 2H), 7.48 (d, *J* = 8.4 Hz, 2H), 7.28 (d, *J* = 8.0 Hz, 2H), 6.80 (dd, *J* = 7.1, 0.7 Hz, 1H), 6.67 (d, *J* = 7.7 Hz, 1H), 4.63 (d, *J* = 6.0 Hz, 2H), 2.64 (s, 3H), 2.37 (s, 3H). ^13^C NMR (126 MHz, acetone-*d6*) δ 167.36, 146.43, 142.49, 138.64, 135.15, 132.71, 132.44, 129.83, 129.75, 128.67, 128.30, 128.13, 127.90, 123.57, 121.37, 120.96, 117.56, 86.73, 43.43, 21.34, 18.21. LC-MS, ESI^+^, *m/z* 459.1 [M+H]^+^.

#### Synthesis of compound 1d

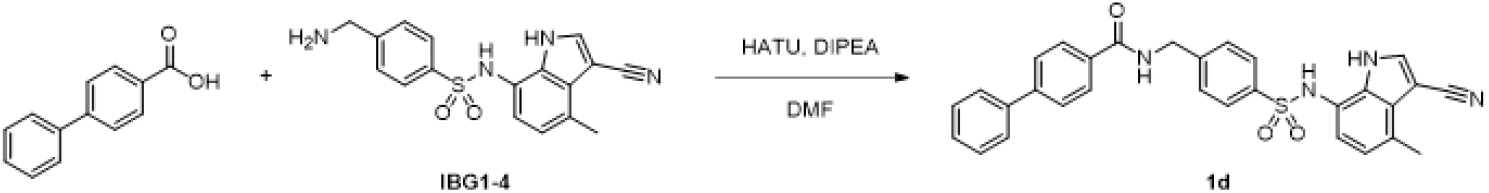

##### *N*-(4-(*N*-(3-Cyano-4-methyl-1H-indol-7-yl)sulfamoyl)benzyl)-[1,1’-biphenyl]-4-carboxamide (1d)

To a solution of 4-(aminomethyl)-*N*-(3-cyano-4-methyl-1H-indol-7-yl)benzenesulfonamide hydrochloride (**IBG1-4**) (30.0 mg, 79.6 µmol), 4-phenylbenzoic acid (18.9 mg, 95.5 µmol, 5.0 eq.), and *N,N*-diisopropylethylamine (69.3 µL, 398 µmol, 5.0 eq.) in *N,N*-dimethylformamide (0.5 mL) was added HATU (45.4 mg, 119 µmol, 1.5 eq.) and then the mixture was stirred at room temperature for 5 hours. The resulted mixture was purified with silica gel column chromatography (ODS, H_2_O-MeCN with 0.1% HCOOH). The obtained compound was triturated with DCM. The solid was collected by filtration, washed with DCM, and then dried in vacuo to afford **1d** (3.69 mg, 9% yield). ^1^H NMR (500 MHz, DMSO-*d6*) δ 11.91 (s, 1H), 9.94 (s, 1H), 9.16 (t, *J* = 6.0 Hz, 1H), 8.16 (s, 1H), 7.99 (d, *J* = 8.5 Hz, 2H), 7.79 (d, *J* = 8.4 Hz, 2H), 7.75-7.73 (m, 2H), 7.68 (d, *J* = 8.4 Hz, 2H), 7.51-7.48 (m, 2H), 7.45 (d, *J* = 8.3 Hz, 2H), 7.41 (t, *J* = 7.4 Hz, 1H), 6.77 (d, *J* = 7.8 Hz, 1H), 6.59 (d, *J* = 7.8 Hz, 1H), 4.55 (d, *J* = 4.6 Hz, 2H), 2.55 (s, 3H). ^13^C NMR (126 MHz, DMSO-*d6*) δ 165.98, 144.96, 142.93, 139.13, 135.21, 132.79, 130.49, 129.03, 128.07, 127.96, 127.52, 127.06, 126.87, 126.58, 126.47, 122.43, 118.11, 117.38, 84.28, 42.23, 17.64. LC-MS, ESI^+^, *m/z* 521.1 [M+H]^+^.

#### Synthesis of compound 1e

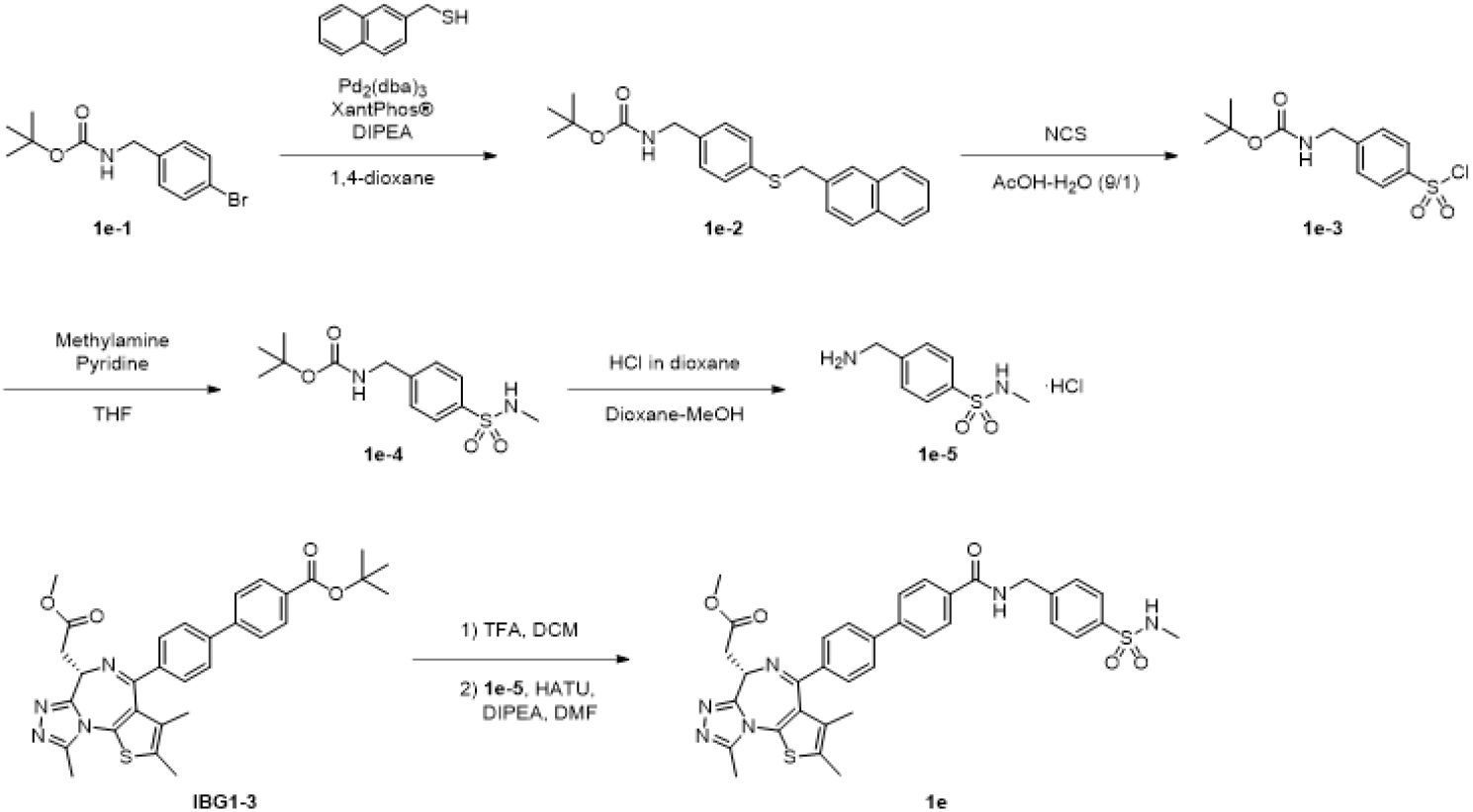

##### *tert*-Butyl (4-((naphthalen-2-ylmethyl)thio)benzyl)carbamate (1e-2)

To a mixture of *tert*-butyl bromobenzylcarbamate (**1e-1**) (500 mg, 1.75 mmol), naphthalen-2-ylmethanethiol (335 mg, 1.92 mmol, 1.1 eq.), tris(dibenzylideneacetone)dipalladium(0) (80.0 mg, 0.09 mmol, 0.05 eq.), XantPhos^®^ (101 mg, 0.17 mmol, 0.1 eq.) was added DIPEA (609 µL, 3.49 mmol, 2.0 eq.) and 1,4-dioxane (10 mL) under nitrogen and then the mixture was stirred at 100 °C for 8 hours. The resulted mixture was concentrated in vacuo and then purified by silica gel column chromatography (heptane-EtOAc) to afford **1e-2** (584 mg, 88% yield). ^1^H NMR (400 MHz, CDCl_3_) δ 7.81-7.73 (m, 3H), 7.67 (s, 1H), 7.47-7.44 (m, 3H), 7.28 (d, *J* = 8.2 Hz, 2H), 7.15 (d, *J* = 7.9 Hz, 2H), 4.76 (bs, 1H), 4.25 (s, 4H), 1.45 (s, 9H). ^13^C NMR (101 MHz, CDCl_3_) δ 155.96, 137.56, 135.24, 135.04, 133.46, 132.75, 130.55, 128.44, 128.11, 127.83, 127.79, 127.53, 127.08, 126.28, 125.96, 79.70, 44.40, 39.72, 28.54. LC-MS, ESI^-^, *m/z* 377.9 [M-H]^-^.

##### *tert*-Butyl (4-(chlorosulfonyl)benzyl)carbamate (1e-3)

To a solution of **1e-2** (250 mg, 659 µmol) in acetic acid (9.0 mL) and water (1.0 mL) was added *N*-chlorosuccinimide (440 mg, 3.29 mmol, 5.0 eq.) and then the mixture was stirred at room temperature for 1.5 hours. The resulted mixture was added to water and then the organic was extracted with toluene. The organic layer was washed with water and brine, dried over MgSO_4_, and then purified by silica gel column chromatography (heptane-EtOAc) to afford **1e-3** (57.6 mg, 29% yield). ^1^H NMR (400 MHz, CDCl_3_) δ 7.98 (d, *J* = 8.3 Hz, 2H), 7.52 (d, *J* = 8.3 Hz, 2H), 5.08 (bs, 1H), 4.42-4.41 (m, 2H), 1.46 (s, 9H). ^13^C NMR (101 MHz, CDCl_3_) δ 155.99, 147.73, 143.36, 128.30, 127.53, 80.43, 44.25, 28.50. LCMS was only detected as sulfonic acid form: LC-MS, ESI^-^, *m/z* 285.8 [M-Cl +O]^-^.

##### *tert*-Butyl (4-(*N*-methylsulfamoyl)benzyl)carbamate (1e-4)

To a solution of **1e-3** (22.1 mg, 72.3 µmol) and pyridine (17.5 µL, 217 µmol, 3.0 eq.) in tetrahydrofuran (1.0 mL) was added 2.0 M methylamine in THF (72.3 µL, 145 µmol, 2.0 eq.) and then the mixture was stirred at room temperature overnight. After the addition of 2.0 M methylamine in THF (72.3 µL, 145 µmol, 2.0 eq.), the mixture was stirred for 2 hours. The resulted mixture was added to water and then the organic was extracted with toluene. The organic layer was washed with water and brine, dried over MgSO_4_, and then purified by silica gel column chromatography (heptane-EtOAc) to afford **1e-4** (17.6 mg, 81% yield). ^1^H NMR (400 MHz, CD_3_OD) δ 7.78 (d, *J* = 6.4 Hz, 2H), 7.46 (d, *J* = 6.4 Hz, 2H), 4.29 (s, 2H), 2.5 (s, 3H), 1.42 (s, 9H). ^13^C NMR (101 MHz, CDCl_3_) δ 155.83, 147.62, 143.12, 128.12, 127.31, 80.23, 44.04, 28.31. LC-MS, ESI^+^, *m/z* 299.6 [M+H]^+^.

##### 4-(Aminomethyl)-N-methylbenzenesulfonamide hydrochloride (1e-5)

To a solution of **1e-4** (17.6 mg, 58.6 µmol) in 1,4-dioxane (293 µL) was added 4 M hydrogen chloride in 1,4-dioxane (293 µL, 1.17 mmol, 20 eq.), and then the mixture was stirred at room temperature for 3.5 hours. Additional 4 M hydrogen chloride in 1,4-dioxane (293 µL, 1.17 mmol, 20 eq.) and MeOH (293 µL) were added, and the mixture was stirred for 2 hours. The resulted mixture was concentrated in vacuo and then used for next reaction without further purification.

##### Methyl (S)-2-(4-(4’-(ethylcarbamoyl)-[1,1’-biphenyl]-4-yl)-2,3,9-trimethyl-6H-thieno[3,2-f][1,2,4]triazolo[4,3-a][1,4]diazepin-6-yl)acetate (1e)

To a solution of **IBG1-3** (18.2 mg, 36.4 µmol) in DCM (0.5 mL) was added trifluoroacetic acid (0.5 mL) and then the mixture was stirred at room temperature for 2 hours. The resulted mixture was concentrated in vacuo, and then toluene was added thereto. After concentrated in vacuo again to afford the corresponding carboxylic acid. To a mixture of the carboxylic acid, 4-(aminomethyl)-*N*-methylbenzenesulfonamide hydrochloride (**1e-5**) (13.8 mg, 58.2 µmol, 1.6 eq.), and *N,N*-diisopropylethylamine (31.7 µL, 182 µmol, 5.0 eq.) in *N,N*-dimethylformamide (1.0 mL) was added HATU (24.9 mg, 65.4 µmol, 1.5 eq.) and then the mixture was stirred at room temperature overnight. After the addition of HATU (24.9 mg, 65.4 µmol, 1.5 eq.) and *N,N*-diisopropylethylamine (31.7 µL, 182 µmol, 5.0 eq.), the mixture was stirred for 2 hours. The resulted mixture was added to ammonium chloride aqueous solution and then the organic was extracted with EtOAc. The obtained organic extract was washed with brine, dried over MgSO_4_, concentrated in vacuo, and then purified by silica gel column chromatography (DCM-MeOH). The obtained compound was purified again by preparative HPLC (ODS, H_2_O-MeCN with 0.1% HCOOH) to afford **1e** (6.8 mg, 27% yield). ^1^H NMR (400 MHz, CD_3_OD) δ 8.00-7.98 (m, 2H), 7.84-7.79 (m, 4H), 7.77-7.75 (m, 2H), 7.59-7.58 (m, 4H), 4.70 (s, 2H), 4.66 (t, J = 7.2 Hz, 1H), 4.52 (s, 1H), 3.79 (s, 3H), 3.58 (d, *J* = 7.2 Hz, 2H), 2.73 (s, 3H), 2.53 (s, 3H), 2.49 (s, 3H), 1.76 (s, 3H). ^13^C NMR (101 MHz, CD_3_OD) δ 171.78, 168.40, 165.71, 155.44, 150.81, 144.09, 143.18, 142.16, 137.96, 137.56, 133.22, 132.02, 131.84, 130.90, 130.84, 129.07, 127.72, 127.68, 127.07, 126.89, 126.84, 53.48, 51.03, 42.71, 35.75, 27.80, 13.01, 11.53, 10.19. HRMS, ESI^+^, *m/z* calcd for C35H35N6O5S2 [M+H]^+^, 683.2110; found 683.215.

#### Synthesis of compound 1f

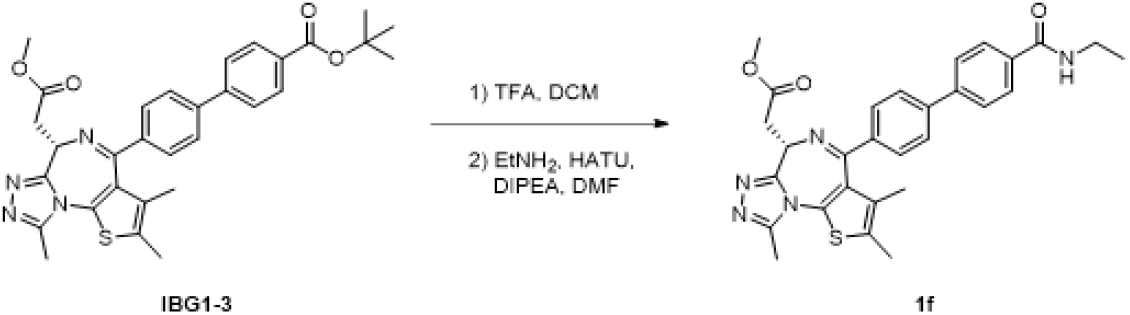

##### Methyl (S)-2-(4-(4’-(ethylcarbamoyl)-[1,1’-biphenyl]-4-yl)-2,3,9-trimethyl-6H-thieno[3,2-f][1,2,4]triazolo[4,3-a][1,4]diazepin-6-yl)acetate (1f)

To a solution of *tert*-butyl (S)-4’-(6-(2-methoxy-2-oxoethyl)-2,3,9-trimethyl-6H-thieno[3,2-f][1,2,4] triazolo[4,3-a][1,4]diazepin-4-yl)-[1,1’-biphenyl]-4-carboxylate (**IBG1-3**) (20.0 mg, 0.03 mmol) in DCM (0.5 mL) was added trifluoroacetic acid (0.5 mL) and then the mixture was stirred at room temperature for 2 hours. The resulted mixture was concentrated in vacuo, and then toluene was added thereto. After concentrated in vacuo again to afford the corresponding carboxylic acid. To the mixture of the carboxylic acid, 2 M ethylamine in THF (49.9 µL, 100 µmol, 2.0 eq.), and *N,N*-diisopropylethylamine (43.5 µL, 250 µmol, 5.0 eq.) in *N,N*-dimethylformamide (1.0 mL) was added HATU (28.5 mg, 74.9 µmol, 1.5 eq.) and then the mixture was stirred at room temperature overnight. The resulted mixture was purified directly by preparative HPLC (ODS, H_2_O-MeCN with 0.1% HCOOH) to afford **1f** (4.9 mg, 19% yield) as a white solid. ^1^H NMR (400 MHz, CD_3_OD) δ 7.91 (d, *J* = 8.4, 2H), 7.77-7.72 (m, 4H), 7.56 (d, *J* = 8.2 Hz, 2H), 4.64 (t, *J* = 7.2 Hz, 1H), 4.50 (s, 1H), 3.78 (s, 3H), 3.56 (d, *J* = 7.2 Hz, 2H), 3.43 (q, *J* = 7.2 Hz, 2H), 2.71 (s, 3H), 2.47 (s, 3H), 1.75 (s, 3H), 1.24 (t, *J* = 7.2 Hz, 3H). ^13^C NMR (101 MHz, CD_3_OD) δ 173.20, 169.60, 167.08, 156.87, 152.17, 144.24, 143.66, 138.90, 135.20, 133.40, 133.22, 132.35, 132.26, 130.43, 128.93, 128.25, 128.11, 54.91, 52.41, 37.20, 35.87, 14.90, 14.38, 12.92, 11.57. HRMS, ESI^+^, *m/z* calcd for C29H30N5O3S [M+H]^+^, 528.2069; found 528.2066.

#### Synthesis of compound 1g

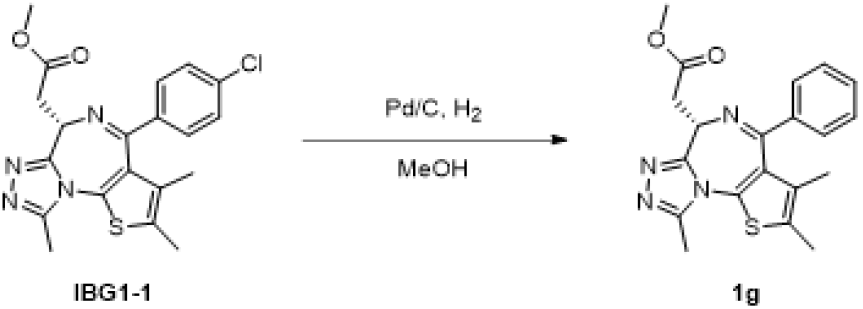

##### Methyl (S)-2-(2,3,9-trimethyl-4-phenyl-6H-thieno[3,2-f][1,2,4]triazolo[4,3-a][1,4]diazepin-6-yl)acetate (1g)

To a solution of methyl (R)-2-(4-(4-chlorophenyl)-2,3,9-trimethyl-6H-thieno[3,2-f][1,2,4]triazolo[4,3-a]azepin-6-yl)acetate (**IBG1-1**) (29.3 mg, 70.6 µmol) in MeOH (1.0 mL) was added palladium on carbon (10wt%, 7.5 mg, 7.1 µmol, 0.1 eq.) under nitrogen and then the mixture was stirred under hydrogen at room temperature overnight. The resulted mixture was diluted with excess amount of EtOAc and then stirred under air at room temperature for 1 hour. The resulted mixture was filtered, washed with EtOAc, and then concentrated by nitrogen blow. The crude mixture was suspended in EtOAc and small amount of DMSO, washed with saturated ammonium chloride aqueous solution, water, and brine, dried over MgSO_4_, concentrated in vacuo, and then purified by silica gel column chromatography (DCM-MeOH). The obtained fraction which had the desired product was concentrated in vacuo and then purified again by preparative HPLC (ODS, H_2_O-MeCN with 0.1% HCOOH) to afford **1g** (2.6 mg, 10%) as a white solid. ^1^H NMR (500 MHz, CDCl_3_) δ 7.48-7.42 (m, 3H), 7.38-7.35 (m, 2H), 4.66-4.64 (m, 1H), 3.80 (s, 3H), 3.69-3.66 (m, 2H), 2.70 (s, 3H), 2.43 (s, 3H), 1.69 (s, 3H), ^13^C NMR (126 MHz, CDCl_3_) δ 172.18, 165.02, 155.48, 149.84, 138.28, 132.10, 131.23, 130.87, 130.51, 130.35, 128.48, 128.43, 53.83, 51.83, 36.80, 14.26, 13.05, 11.83. HRMS, ESI^+^, *m/z* calcd for C20H21N4O2S [M+H]^+^, 381.1385; found 381.1389.

#### Synthesis of bIBG1

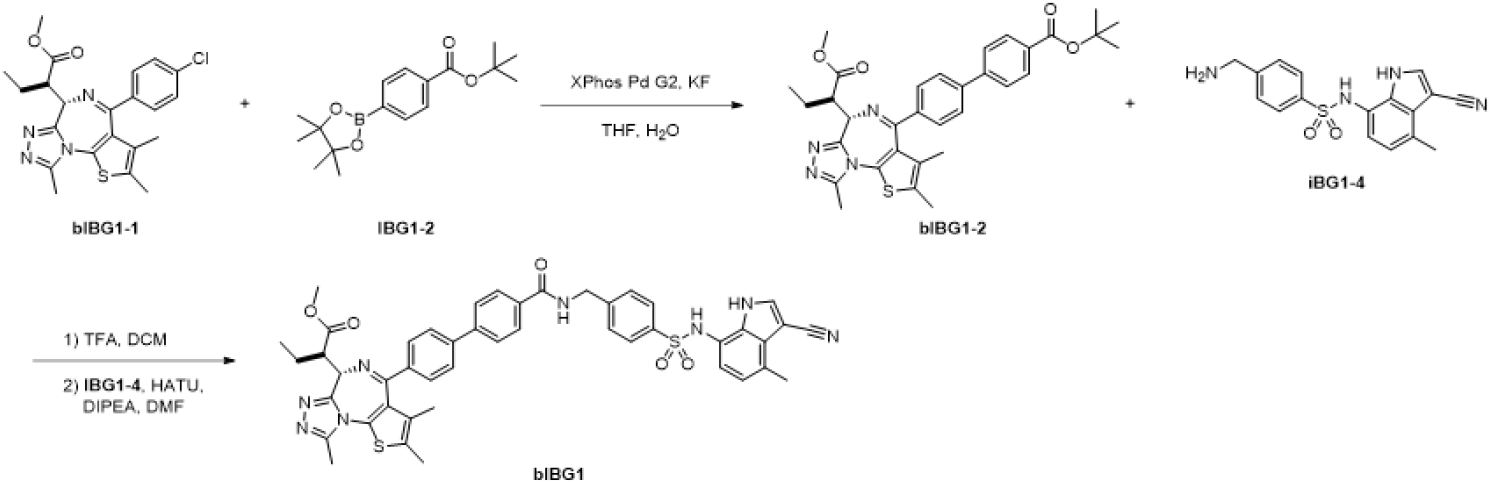

##### tert-Butyl 4’-((S)-6-((R)-1-methoxy-1-oxobutan-2-yl)-2,3,9-trimethyl-6H-thieno[3,2-f][1,2,4]triazolo[4,3-a][1,4]diazepin-4-yl)-[1,1’-biphenyl]-4-carboxylate (bIBG1-2)

A solution of methyl (R)-2-((S)-4-(4-chlorophenyl)-2,3,9-trimethyl-6H-thieno[3,2-f][1,2,4]triazolo[4,3-a][1,4]diazepin-6-yl)butanoate (**bIBG1-1**) (20 mg, 0.027 mmol), potassium fluoride (16 mg, 0.27 mmol, 10 eq.), 4-(tert-butoxycarbonyl)phenylboronic acid pinacol ester (24 mg, 0.080 mmol, 3.0 eq.) and [2-(2-aminophenyl)phenyl]-hydroxy-oxo-palladium [2-chloro-6-(2,4,6-triisopropylphenyl)phenyl]-dicyclohexylphosphane (6.6 mg, 0.0080 mmol, 0.30 eq.) in DMF (3.0 mL) and water (0.15 mL) was heated to 100 °C. After being stirred for 15 hours, the mixture was filtered and washed with water and saturated aqueous NaCl. The organic layer was extracted, dried over Na_2_SO_4_, filtered, and concentrated. Column chromatography of the residue (DCM-MeOH) gave the impure product, which was purified again with preparative HPLC (ODS, H_2_O-MeCN with 0.1% HCOOH) to give **bIBG1-2** (9.0 mg, 0.015 mmol, 58% yield).^1^H NMR (400 MHz, CD_3_OD) δ 8.05 (d, *J* = 8.4 Hz, 2H), 7.79-7.71 (m, 4H), 7.51 (d, *J* = 8.3 Hz, 2H), 4.29 (d, *J* = 11.0 Hz, 1H), 3.90 (s, 3H), 3.88 (m, 1H), 2.72 (s, 3H), 2.49 (s, 3H), 2.09 (m, 1H), 1.75 (s, 3H), 1.71 (m, 1H), 1.63 (s, 9H), 1.06 (t, *J* = 7.3 Hz, 3H). ^13^C NMR (101 MHz, CD_3_OD) δ 177.28, 167.91, 167.11, 156.70, 152.97, 146.24, 144.43, 139.68, 134.23, 134.09, 133.35, 133.18, 132.95, 131.83, 131.22, 129.18, 128.88, 83.27, 61.38, 53.10, 51.93, 29.30, 25.06, 15.26, 13.80, 12.68, 12.41. LC-MS, ESI^+^, *m/z* 442.9 [M+H]^+^.

##### methyl (R)-2-((S)-4-(4’-((4-(N-(3-Cyano-4-methyl-1H-indol-7-yl)sulfamoyl)benzyl)carbamoyl)-[1,1’-biphenyl]-4-yl)-2,3,9-trimethyl-6H-thieno[3,2-f][1,2,4]triazolo[4,3-a][1,4]diazepin-6-yl)butanoate (bIBG1)

To a solution of **bIBG1-2** (9.0 mg, 0.0154 mmol) in DCM (0.5 mL) was added trifluoroacetic acid (0.50 mL, 6.53 mmol). After being stirred for 3 hours, the mixture was concentrated. The residue was azeotropically dried three times with toluene. To a solution of the resultant residue and **IBG1-4** (15 mg, 0.023 mmol, 1.5 eq.) in DMF (0.8 mL) was added HATU (12 mg, 0.030 mmol, 2.0 eq.) and *N,N*-dimethylformamide (0.013 mL, 0.076 mmol, 5.0 eq.). After being stirred for 2 hours, the mixture was purified with preparative HPLC (ODS, H_2_O-MeCN with 0.1% HCOOH) to give **bIBG1** (3.9 mg, 0.0046 mmol, 30% yield).^1^H NMR (500 MHz, CD_3_OD) δ 8.00-7.95 (m, 2H), 7.92 (s, 1H), 7.83-7.34 (m, 4H), 7.69-7.62 (m, 2H), 7.56-7.45 (m, 4H), 6.75 (m, 1H), 6.52 (m, 1H), 4.66 (s, 2H), 4.30 (d, *J* = 11.0 Hz, 1H), 3.90 (s, 3H), 3.87 (dd, *J* = 10.9, 3.7 Hz, 1H), 2.73 (s, 3H), 2.66 (s, 3H), 2.50 (s, 3H), 2.10 (m, 1H), 1.77 (s, 3H), 1.71 (m, 1H), 1.06 (t, *J* = 7.45 Hz, 3H) ^13^C NMR (126 MHz, CD_3_OD) δ 177.31, 170.62, 167.24, 156.76, 153.00, 146.76, 145.43, 144.43, 139.95, 139.61, 136.36, 135.44, 134.27, 134.13, 133.17, 132.96, 131.27, 131.17, 129.96, 129.66, 129.63, 129.14, 129.10, 124.58, 122.52, 122.44, 119.37, 86.94, 61.37, 53.14, 51.92, 44.90, 25.08, 19.13, 15.28, 13.80, 12.69, 12.42. HRMS, ESI^+^, *m/z* calcd for C46H43N8O5S2 [M+H]^+^, 851.2798; found 851.2834.

#### Synthesis of IBG2

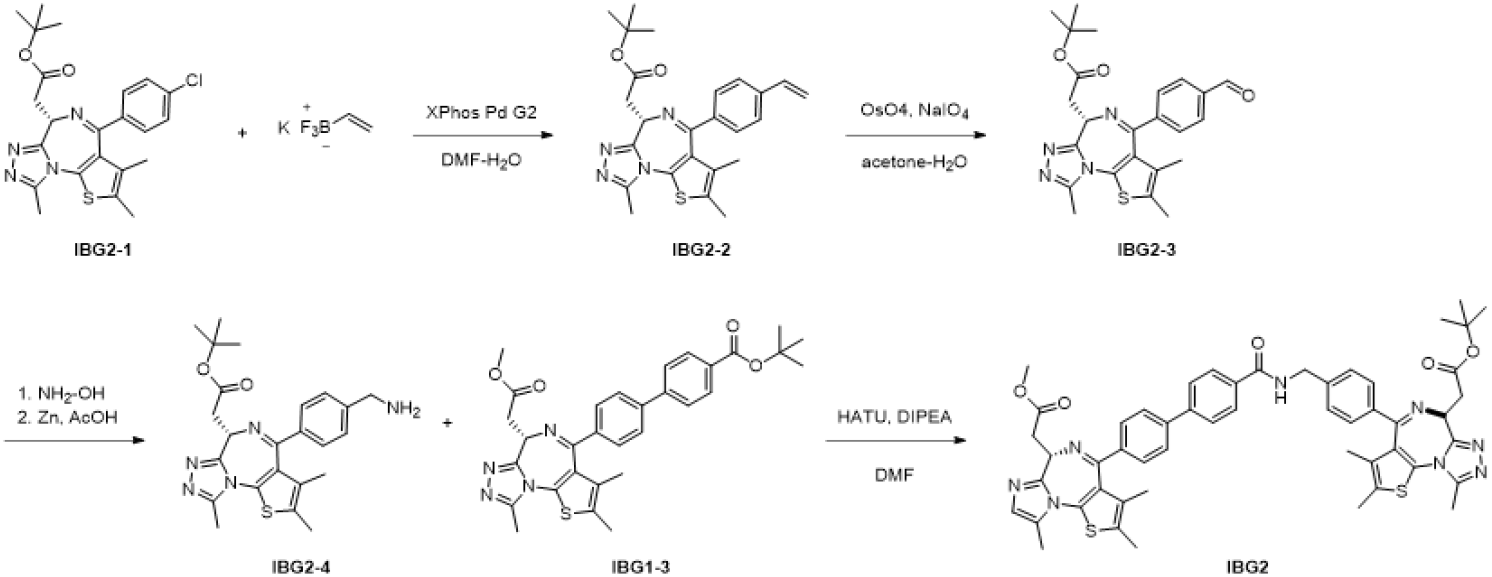

##### *tert-*Butyl (S)-2-(2,3,9-trimethyl-4-(4-vinylphenyl)-6H-thieno[3,2-f][1,2,4]triazolo[4,3-a][1,4]diazepin-6-yl)acetate (IBG2-2)

A solution of JQ1 (**IBG2-1**) (240 mg, 0.525 mmol), potassium vinyltrifluoroborate (211 mg, 1.58 mmol, 3.0 eq.), [2-(2-aminophenyl)phenyl]-chloro-palladium dicyclohexyl-[2-(2,4,6-triisopropylphenyl)phenyl]phosphane (124 mg, 0.158 mmol, 0.30 eq.) and *N,N*-diisopropylethylamine (0.46 mL, 2.63 mmol, 5.0 eq.) in DMF (3.0 mL) and water (0.3 mL) was heated to 130 °C. After being stirred for 15 hours, the mixture was filtered and then washed with water and saturated aqueous NaCl. The organic layer was extracted, dried over Na_2_SO_4_, filtered, and concentrated. Column chromatography of the residue (DCM-MeOH) gave **IBG2-2** (180 mg, 0.40 mmol, 76% yield). ^1^H NMR (500 MHz, CDCl_3_) δ 7.42 (d, *J* = 8.3 Hz, 2H), 7.38 (d, *J* = 8.6 Hz, 2H), 6.71 (dd, *J* = 17.6, 10.9 Hz, 1H), 5.80 (d, *J* = 17.6 Hz, 1H), 5.31 (d, *J* = 11.0 Hz, 1H), 4.56 (dd, *J* = 7.2, 6.9 Hz, 1H), 3.55 (d, *J* = 6.9 Hz, 2H), 2.67 (s, 3H), 2.40 (s, 3H), 1.69 (s, 3H), 1.50 (s, 9H). ^13^C NMR (126 MHz, CDCl_3_) δ 171.00, 164.41, 155.77, 149.85, 139.71, 137.69, 136.25, 132.15, 131.21, 130.94, 130.43, 128.82, 126.34, 115.46, 80.94, 54.00, 38.05, 28.30, 14.48, 13.20, 12.00. LC-MS, ESI^-^, *m/z* 449.1 [M+H]^+^.

##### *tert-*Butyl (S)-2-(4-(4-formylphenyl)-2,3,9-trimethyl-6H-thieno[3,2-f][1,2,4]triazolo[4,3-a][1,4]diazepin-6-yl)acetate (IBG2-3)

To a solution of **IBG2-2** (200 mg, 0.446 mmol, 1.0 eq.) and sodium periodate (286 mg, 1.34 mmol, 3.0 eq.) in acetone (5.0 mL) and water (1.0 mL) was added osmium tetroxide (0.14 mL as a 4% aqueous solution, 0.0223 mmol, 0.050 eq.). The reaction was stirred for 2 hours, and then diluted with EtOAc. The organic layer was washed with water and saturated aqueous NaCl, dried over Na_2_SO_4_, filtered, and concentrated. Column chromatography of the residue (DCM-MeOH) to give **IBG2-3** (180 mg,0.40 mmol, 90% yield). ^1^H NMR (500 MHz, CDCl_3_) δ 10.04 (s, 1H), 7.87 (d, *J* = 8.4, 2H), 7.63 (d, *J* = 8.1 Hz, 2H), 4.61 (dd, *J* = 7.0, 7.0 Hz, 1H), 3.57 (d, *J* = 7.0 Hz, 2H), 2.69 (s, 3H), 2.41 (s, 3H), 1.66 (s, 3H), 1.51 (s, 9H). ^13^C NMR (126 MHz, CDCl_3_) δ 191.78, 170.87, 163.96, 155.35, 149.99, 143.68, 137.51, 132.63, 131.00, 130.72, 130.22, 129.82, 129.24, 81.12, 54.30, 37.91, 28.29, 14.48, 13.22, 12.01. LC-MS, ESI^-^, *m/z* 451.2 [M+H]^+^.

##### *tert-*Butyl (S)-2-(4-(4-(aminomethyl)phenyl)-2,3,9-trimethyl-6H-thieno[3,2-f][1,2,4]triazolo[4,3-a][1,4]diazepin-6-yl)acetate (IBG2-4)

To a solution of **IBG2-3** (30 mg, 0.0666 mmol) in ethanol (1.0 mL) was added hydroxylamine hydrochloride (16 mg, 0.226 mmol, 3.4 eq.) and sodium acetate (22 mg, 0.266 mmol, 4.0 eq.). The reaction mixture was stirred at 70 °C for 3 hours. The mixture was cooled to room temperature and diluted with EtOAc, and then quenched with water. The organic layer was washed with saturated aqueous NaCl, dried over Na_2_SO_4_, filtered, and concentrated. The crude was used in the next reaction without further purifications. LC-MS, ESI^-^, *m/z* 446.2 [M+H]^+^.

A mixture of the resultant crude and zinc (8.4 mg, 0.129 mmol) were dissolved in acetic acid (1.0 mL) and stirred at 40 °C. After being stirred for 20 hours, another 5 mg of zinc was added. The reaction was stirred for 2 hours, and then added another 1 mg of zinc. After being stirred for additional 3 hours, the reaction was diluted with DCM and filtered. The filtrate was concentrated and purified with preparative HPLC (ODS, H_2_O-MeCN with 0.1% HCOOH) to give **IBG2-4** (13 mg, 0.0288 mmol, 45% yield). ^1^H NMR (500 MHz, CD_3_OD) δ 7.46 (d, *J* = 8.3 Hz, 2H), 7.41 (d, *J* = 8.4 Hz, 2H), 4.58 (dd, *J* = 8.7, 5.8 Hz, 1H), 3.88 (brs, 2H), 3.53-3.38 (m, 2H), 2.73 (s, 3H), 2.48 (s, 3H), 1.72 (s, 3H), 1.53 (s, 9H). ^13^C NMR (126 MHz, CD_3_OD) δ 172.73, 168.02, 157.79, 152.90, 147.09, 138.92, 134.13, 133.95, 133.28, 133.04, 130.79, 129.50, 83.19, 55.85, 47.01, 39.34, 29.26, 15.17, 13.78, 12.42. LCMS was only detected as sulfonic acid form: LC-MS, ESI^-^, *m/z* 452.10 [M+H]^+^.

##### *tert-*Butyl 2-((S)-4-(4-((4’-((S)-6-(2-(tert-butoxy)-2-oxoethyl)-2,3,9-trimethyl-6H-thieno[3,2-f][1,2,4]triazolo[4,3-a][1,4]diazepin-4-yl)-[1,1’-biphenyl]-4-carboxamido)methyl)phenyl)-2,3,9-trimethyl-6H-thieno[3,2-f][1,2,4]triazolo[4,3-a][1,4]diazepin-6-yl)acetate (**IBG2**)

To a solution of **IBG1-3** (10.0 mg, 0.018 mmol) in DCM (0.5 mL) was added trifluoroacetic acid (0.5 mL), and then the mixture was stirred at room temperature for 5 hours. The reaction mixture was concentrated, and the residue was azeotropically dried three times with toluene. To a solution of the resultant crude and **IBG2-4** (5.4 mg, 0.012 mmol) in DMF (0.5 mL) was added HATU (9.1 mg, 0.0240 mmol, 2.0 eq.) and *N,N*-diisopropylethylamine (0.010 mL, 0.0599 mmol, 5.0 eq.). After being stirred for 3 hours, the mixture was diluted with EtOAc, and then quenched with aqueous NaHCO_3_. The mixture was extracted with EtOAc, and the organic layer was washed with saturated aqueous NaCl, dried over Na_2_SO_4_, filtered, and concentrated. The residue was purified with preparative HPLC (ODS, H_2_O-MeCN with 0.1% HCOOH) to give **IBG2** (6.0 mg, 0.00642 mmol, 54% yield). ^1^H NMR (500 MHz, CD_3_OD) δ 7.97 (d, *J* = 8.4 Hz, 2H), 7.79 (d, *J* = 8.4 Hz, 2H), 7.75 (d, *J* = 8.5 Hz, 2H), 7.58 (d, *J* = 8.1 Hz, 2H), 7.49-7.41 (m, 4H), 4.70-464 (m, 3H), 4.57 (dd, *J* = 8.6, 5.9 Hz, 1H), 3.80 (s, 3H), 3.59 (d, *J* = 7.5 Hz, 2H), 3.44 (m, 2H), 2.73 (s, 3H), 2.71 (s, 3H), 2.49 (s, 3H), 2.46 (s, 3H), 1.76 (s, 3H), 1.70 (s, 3H), 1.51 (s. 9H). ^13^C NMR (126 MHz, CD_3_OD) δ 174.02, 172.75, 170.52, 168.00, 167.91, 157.76, 157.67, 153.04, 152.92, 145.30, 144.37, 144.28, 139.76, 139.08, 135.57, 134.24, 134.14, 134.07, 133.98, 133.22, 133.12, 133.07, 133.04, 131.30, 130.79, 129.95, 129.47, 129.11, 129.04, 83.23, 55.86, 55.72, 53.31, 45.04, 39.34, 38.02, 29.26, 15.29, 15.22, 13.81, 13.79, 12.47, 12.43. HRMS, ESI+, m/z calcd for C51H52N9O5S2 [M+H]^+^, 934.353; found 934.359.

#### Synthesis of IBG3

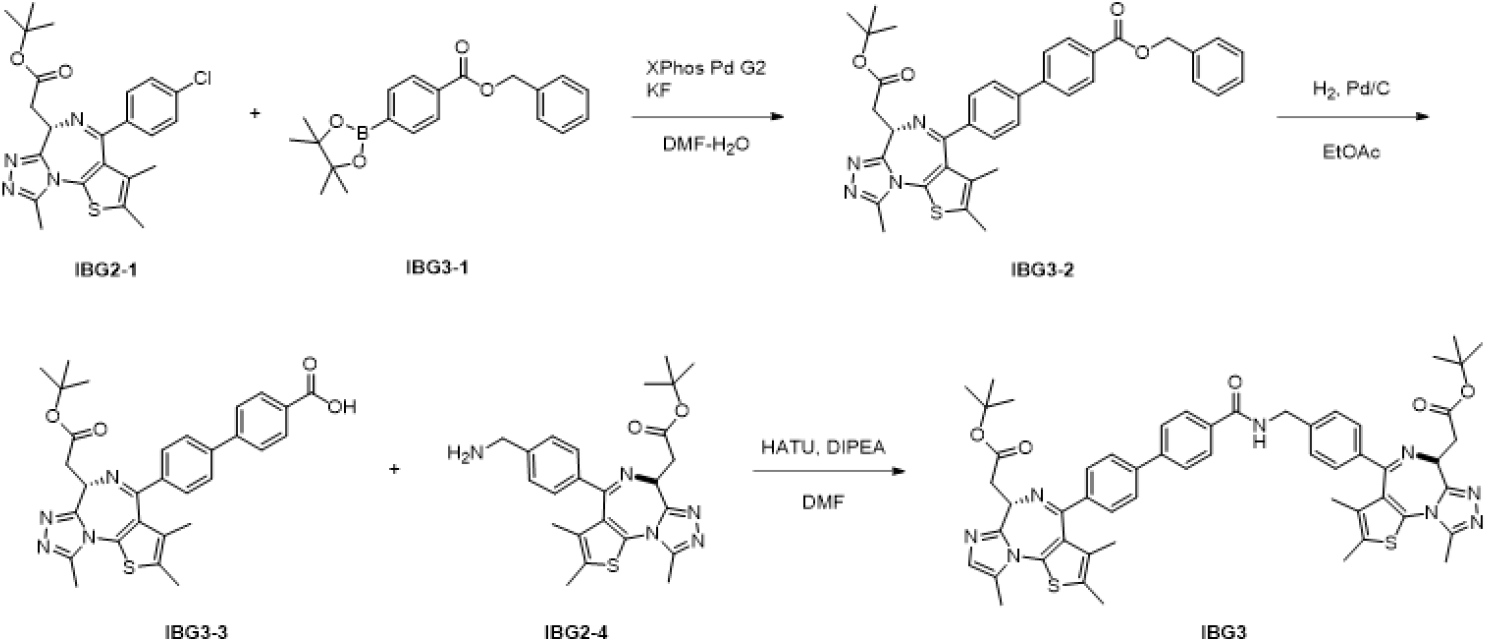

##### Benzyl (S)-4’-(6-(2-(tert-butoxy)-2-oxoethyl)-2,3,9-trimethyl-6H-thieno[3,2-f][1,2,4]triazolo[4,3-a][1,4]diazepin-4-yl)-[1,1’-biphenyl]-4-carboxylate (**IBG3-2**)

A solution of tert-butyl 2-[rac-(9S)-7-(4-chlorophenyl)-4,5,13-trimethyl-3-thia-1,8,11,12-tetrazatricyclo[8.3.0.02,6]trideca-2(6),4,7,10,12-pentaen-9-yl]acetate JQ1 (**IBG2-1**) (50 mg, 0.109 mmol, 1.0 eq.), benzyl 4-(4,4,5,5-tetramethyl-1,3,2-dioxaborolan-2-yl)benzoate (**IBG3-1**) (66.6 mg, 0.197 mmol, 1.8 eq.), [2-(2-aminophenyl)phenyl]-hydroxy-oxo-palladium [2-chloro-6-(2,4,6-triisopropylphenyl)phenyl]-dicyclohexylphosphane (27 mg, 0.0328 mmol, 0.3 eq.), and potassium fluoride (32 mg, 0.547 mmol, 5.0 eq.) in DMF (3.0 mL) and water (0.15 mL) was heated to 130 °C. After being stirred for 15 hours, the mixture was filtered and then, washed with water and saturated aqueous NaCl. The organic layer was extracted, dried over Na_2_SO_4_, filtered, and concentrated. Column chromatography of the residue (DCM-MeOH) gave impure product, which was purified with preparative HPLC (ODS, H_2_O-MeCN with 0.1% HCOOH) gave **IBG3-2** (38 mg, 0.060 mmol, 55% yield). ^1^H NMR (500 MHz, CDCl_3_) δ 8.11 (m, 2H), 7.75 (m, 4H), 7.57 (d, *J* = 7.6 Hz, 2H), 7.48 (d, *J* = 7.8 Hz, 2H), 7.43-7.32 (m, 3H), 5.38 (s, 2H), 4.59 (dd, *J* = 8.7, 5.8 Hz, 1H), 3.54-3.40 (m, 2H), 2.72 (s, 3H), 2.46 (s, 3H), 1.74 (s, 3H), 1.53 (s, 9H). ^13^C NMR (126 MHz, CDCl_3_) δ 170.51, 166.10, 165.37, 155.47, 150.67, 144.44, 141.94, 137.65, 136.17, 131.98, 131.81, 130.85, 130.71, 129.85, 129.26, 129.00, 128.24, 128.12, 128.02, 127.93, 127.85, 126.96, 126.82, 80.98, 53.72, 37.16, 13.05, 11.58, 10.22. LC-MS, ESI^-^, *m/z* 633.2 [M+H]^+^.

##### (S)-4’-(6-(2-(tert-butoxy)-2-oxoethyl)-2,3,9-trimethyl-6H-thieno[3,2-f][1,2,4]triazolo[4,3-a][1,4]diazepin-4-yl)-[1,1’-biphenyl]-4-carboxylic acid (**IBG3-3**)

To a solution of **IBG3-2** (38 mg, 0.0601 mmol) in EtOAc (1.0 mL) was added palladium on carbon (5%, 6.4 mg, 0.05 eq.), and the mixture was stirred under hydrogen atmosphere at room temperature for 5 hours. The suspension was filtered through a pad of silica gel with 20% MeOH in DCM. Without further purifications the crude was used in the next reaction. LC-MS, ESI^-^, *m/z* 543.2 [M+H]^+^.

##### tert-Butyl 2-((S)-4-(4-((4’-((S)-6-(2-(tert-butoxy)-2-oxoethyl)-2,3,9-trimethyl-6H-thieno[3,2-f][1,2,4]triazolo[4,3-a][1,4]diazepin-4-yl)-[1,1’-biphenyl]-4-carboxamido)methyl)phenyl)-2,3,9-trimethyl-6H-thieno[3,2-f][1,2,4]triazolo[4,3-a][1,4]diazepin-6-yl)acetate (IBG3)

To a solution of **IBG2-4** (13 mg, 0.0288 mmol) and **IBG3-3** (17 mg, 0.0313 mmol, 1.1 eq.) in DMF (0.5 mL) was added HATU (22 mg, 0.0576 mmol, 2.0 eq.) and *N,N*-diisopropylethylamine (0.025 mL, 0.144 mmol, 5.0 eq.). After being stirred for 3 hours, the mixture was diluted with EtOAc, and then quenched with aqueous NaHCO_3_. The mixture was extracted with EtOAc, and the organic layer was washed with saturated aqueous NH4Cl and NaCl, dried over Na_2_SO_4_, filtered, and concentrated. The residue was purified with preparative HPLC (ODS, H_2_O-MeCN with 0.1% HCOOH) to give **IBG3** (10.0 mg, 0.0102 mmol, 36% yield). ^1^H NMR (500 MHz, CDl_3_OD) δ 8.00-7.94 (m, 2H), 7.82-7.72 (m, 4H), 7.62-7.56 (m, 2H), 7.49-7.40 (m, 4H), 4.66 (brs, 2H), 4.61 (dd, *J* = 8.6, 5.8 Hz, 1H), 4.56 (dd, *J* = 8.6, 5.9 Hz, 1H), 3.67-3.55 (m, 4H), 2.73 (s, 3H), 2.71 (s, 3H), 2.49 (s, 3H), 2.46 (s, 3H), 1.76, (s, 3H), 1.70 (s, 3H), 1.54 (s, 9H), 1.51 (s, 9H). ^13^C NMR (126 MHz, CD_3_OD) δ 172.74, 172.72, 170.50, 167.95, 167.70, 157.77, 152.94, 145.28, 144.38, 144.28, 139.74, 139.09, 135.58, 134.25, 134.14, 134.06, 133.95, 133.23, 133.16, 133.04, 132.99, 131.25, 130.79, 129.94, 129.45, 129.11, 129.04, 83.20, 83.18, 55.97, 55.86, 45.04, 39.40, 39.35, 29.28, 29.26, 15.28, 15.22, 13.81, 13.78, 12.45, 12.42. HRMS, ESI+, m/z calcd for C54H58N9O5S2 [M+H]^+^, 976.4002; found 976.4047.

#### Synthesis of IBG4

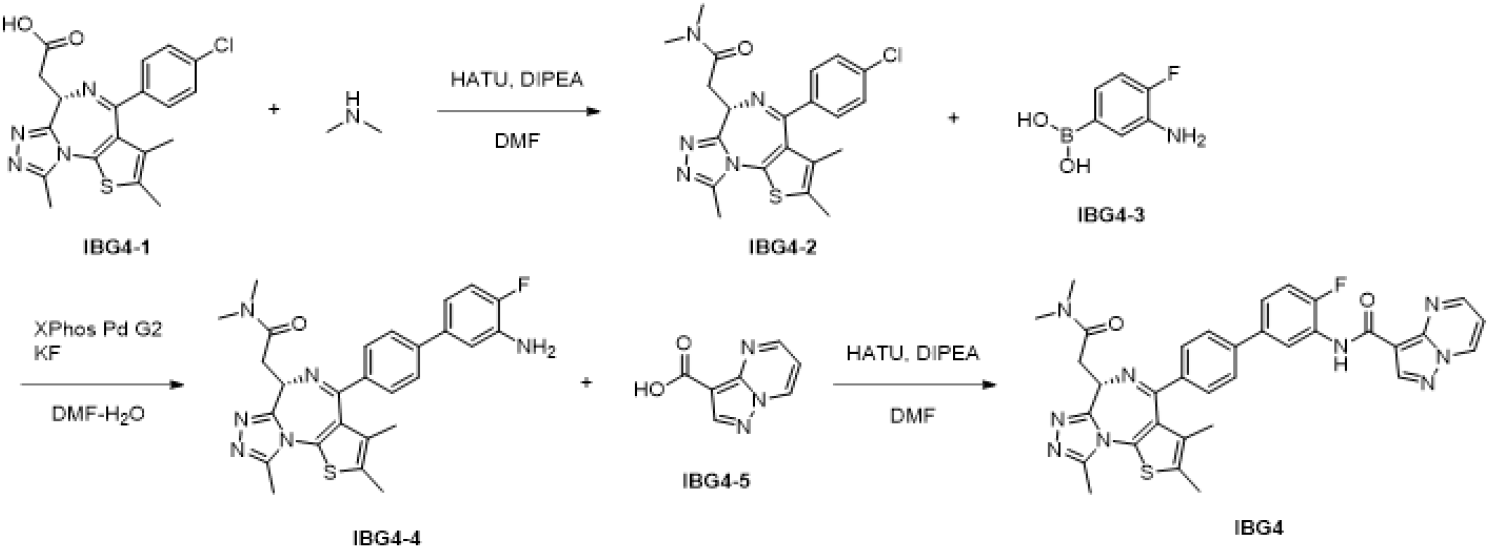

##### (S)-2-(4-(4-Chlorophenyl)-2,3,9-trimethyl-6H-thieno[3,2-f][1,2,4]triazolo[4,3-a][1,4]diazepin-6-yl)-N,N-dimethylacetamide (IBG4-2)

To a solution of (S)-2-(4-(4-chlorophenyl)-2,3,9-trimethyl-6H-thieno[3,2-f][1,2,4]triazolo[4,3-a][1,4]diazepin-6-yl)acetic acid (**IBG4-1**) (87 mg, 0.218 mmol) in DMF (0.5 mL) was added *N,N*-diisopropylethylamine (0.19 mL, 1.09 mmol, 5.0 eq.), dimethylamine (2 M in THF, 1.1 mL, 2.18 mmol, 10 eq.), and HATU (166 mg, 0.436 mmol, 2.0 eq.). After being stirred at room temperature for 3 hours, the mixture was diluted with EtOAc, and then quenched with aqueous NaHCO_3_. The mixture was extracted with EtOAc, and the organic layer was washed with saturated aqueous NaCl, dried over Na_2_SO_4_, filtered, and concentrated. Preparative HPLC purification (ODS, H_2_O-MeCN with 0.1% HCOOH) of the residue gave **IBG4-2** (75 mg, 0.175 mmol, 80% yield). ^1^H NMR (500 MHz, CDCl_3_) δ 7.39 (d, *J* = 8.5 Hz, 2H), 7.31 (d, *J* = 8.7 Hz, 2H), 4.79 (dd, *J* = 7.4, 6.0 Hz, 1H), 3.66 (dd, *J =* 16.1, 5.1 Hz, 1H), 3.58 (dd, *J* = 16.1, 7.5 Hz, 1H), 3.24 (s, 3H), 3.00 (s, 3H), 2.66 (s, 3H), 2.38 (s, 3H), 1.66 (s, 3H). ^13^C NMR (126 MHz, CDCl_3_) δ 170.60, 163.70, 156.08, 149.88, 136.98, 136.68, 132.31, 131.03, 130.69, 129.94, 128.76, 54.62, 37.67, 35.69, 35.63, 14.48, 13.19, 11.96. LC-MS, ESI^-^, *m/z* 428.1 [M+H]^+^.

##### (S)-2-(4-(3’-amino-4’-fluoro-[1,1’-biphenyl]-4-yl)-2,3,9-trimethyl-6H-thieno[3,2-f][1,2,4]triazolo[4,3-a][1,4]diazepin-6-yl)-N,N-dimethylacetamide (IBG4-4)

A mixture of **IBG4-2** (38 mg, 0.0876 mmol), (3-amino-4-fluoro-phenyl)boronic acid (**IBG4-3**) (41 mg, 0.263 mmol, 3.0 eq.), potassium fluoride (25 mg, 0.438 mmol, 5.0 eq.), and [2-(2-aminophenyl)phenyl]-chloro-palladium;dicyclohexyl-[2-(2,4,6-triisopropylphenyl)phenyl]phosphane (21 mg, 0.0263 mmol, 0.3 eq.) was dissolved in DMF (1.0 mL) and water (0.15 mL). After being stirred at 130 °C for 20 hours, the reaction was cooled to room temperature, diluted with EtOAc and then quenched with saturated aqueous NaHCO_3_. The mixture was extracted with EtOAc, and the organic layer was washed with saturated aqueous NaCl, dried over Na_2_SO_4_, filtered, and concentrated. Column chromatography (ODS, H_2_O-MeCN with 0.1% HCOOH) of the residue gave the product as a salt of formic acid. The product was dissolved in EtOAc. and washed with saturated aqueous NaHCO_3_, dried over Na_2_SO_4_, filtered, and concentrated to give **IBG4-4** (20 mg, 0.0398 mmol, 45% yield). ^1^H NMR (500 MHz, CDCl_3_) δ 7.48 (brs, 4H), 7.05-6.95 (m, 2H), 6.89 (m, 1H), 4.82 (dd, *J* = 6.1, 1.2 Hz, 1H), 3.82 (brs, 2H), 3.67 (dd, *J* = 9.9, 6.1 Hz, 1H), 3.60 (dd, *J* = 8.6, 7.4 Hz, 1H), 3.26 (s, 3H), 3.01 (s, 3H), 2.66 (s, 3H), 2.69 (s, 3H), 1.70 (s, 3H). ^13^C NMR (126 MHz, CDCl_3_) δ 170.73, 164.48, 156.22, 151.73 (d, *J* = 240.2 Hz), 149.83, 142.70, 137.30, 136.95 (d, *J* = 3.1 Hz), 134.87 (d, *J* = 13.3 Hz), 132.13, 131.30, 131.09, 130.39, 128.96, 126.98 117.47 (d, *J* = 6.7 Hz), 115.66 (d, *J* = 14.9 Hz), 115.54 (d, *J* = 7.9 Hz), 54.64, 37.69, 35.71, 35.68, 14.51, 13.18, 11.97. LC-MS, ESI^-^, *m/z* 503.1 [M+H]^+^.

##### (S)-N-(4’-(6-(2-(Dimethylamino)-2-oxoethyl)-2,3,9-trimethyl-6H-thieno[3,2-f][1,2,4]triazolo[4,3-a][1,4]diazepin-4-yl)-4-fluoro-[1,1’-biphenyl]-3-yl)pyrazolo[1,5-a]pyrimidine-3-carboxamide (IBG4)

To a solution of **IBG4-4** (20 mg, 0.0398 mmol), pyrazolo[1,5-a]pyrimidine-3-carboxylic acid (**IBG4-5**) (13 mg, 0.0796 mmol, 2.0 eq.), and HATU (30 mg, 0.0796 mmol, 2.0 eq.) in DMF (0.5 mL) was added *N,N*-diisopropylethylamine (0.035 mL, 0.199 mmol, 5.0 eq.) and 4-(dimethylamino)pyridin (24 mg, 0.199 mmol, 5.0 eq.). After being stirred at room temperature for 2 hours, another pyrazolo[1,5-a]pyrimidine-3-carboxylic acid (13 mg, 0.0796 mmol, 2.0 eq.), HATU (30 mg, 0.0796 mmol, 2.0 eq.), and *N,N*-diisopropylethylamine (0.035 mL, 0.199 mmol, 5.0 eq.) were added. The reaction was stirred for additional 90 hours at room temperature. The reaction was diluted with EtOAc and quenched with saturated aqueous NaHCO_3_. The organic layer was dried over Na_2_SO_4_, filtered, and concentrated. Preparative HPLC purification (ODS, H_2_O-MeCN with 0.1% HCOOH) of the residue gave **IBG4** (2.5 mg, 0.00389 mmol, 10% yield). ^1^H NMR (500 MHz, CD_3_OD) δ 9.15 (dd, *J* = 5.4, 1.6 Hz, 1H), 8.89 (dd, *J* = 2.6, 1.6 Hz, 1H), 8.80 (dd, *J* = 5.1, 2.3 Hz, 1H), 8.70 (s, 1H), 7.72 (d, *J* = 8.6 Hz, 2H), 7.57 (d, *J* = 8.3 Hz, 2H), 7.45 (m, 1H), 7.33 (dd, *J* = 10.7, 8.6 Hz, 1H), 7.30 (dd, *J* = 4.2, 2.8 Hz, 1H), 4.74 (dd, *J* = 7.4, 6.5 Hz, 1H), 3.69 (dd, *J* = 16.4, 7.5 Hz, 1H), 3.63 (dd, *J* = 16.4, 6.4 Hz, 1H), 3.33 (s, 3H), 3.06 (s, 3H), 2.76 (s, 3H), 2.50 (s, 3H), 1.78 (s. 3H). ^13^C NMR (126 MHz, CD_3_OD) δ 173.27, 167.89, 163.19, 158.24, 155.06, 154.82 (d, *J* = 245 Hz), 152.97, 152.50, 148.38, 148.27, 144.69, 139.55, 139.36, 138.84, 134.20, 133.94, 133.27 (d, *J* = 77.8 Hz), 131.25, 129.20 (d, *J* = 10.6 Hz), 128.94, 124.95 (d, *J* = 7.7 Hz), 122.44, 117.38 (d, *J* = 19.7 Hz), 112.32, 107.02, 56.33, 38.68, 37.05, 36.70, 15.26, 13.80, 12.47. HRMS, ESI+, m/z calcd for C34H31N9O2SF [M+H]+, 648.2305; found 648.2334.

#### Synthesis of DAT389

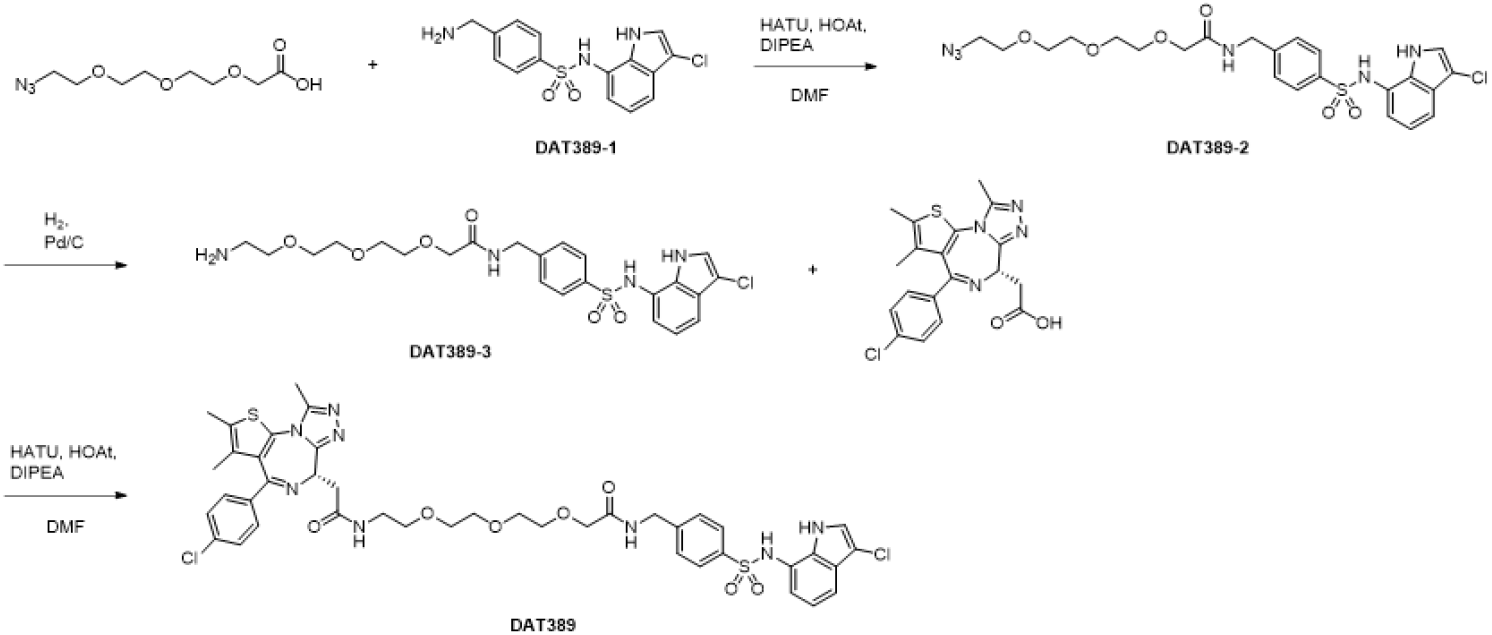

##### 2-(2-(2-(2-Azidoethoxy)ethoxy)ethoxy)-N-(4-(N-(3-chloro-1H-indol-7-yl)sulfamoyl)benzyl)acetamide (DAT389-2)

To a solution of 4-(aminomethyl)-*N*-(3-chloro-1H-indol-7-yl)benzenesulfonamide (19.7 mg, 0.059 mmol), 2-[2-[2-(2-azidoethoxy)ethoxy]ethoxy]acetic acid (16.4 mg, 0.070 mmol, 1.2 eq.) (**DAT389-1**)^10^ *N,N*-diisopropylethylamine (50 µl, 0.29 mmol, 5.0 eq.) and 1-hydroxy-7-azabenzotriazole (6.3 mg, 0.046 mmol, 0.8 eq.) in DMF (0.5 mL) was added HATU (24.5 mg, 0.065 mmol, 1.1 eq.) at room temperature. The reaction was stirred for 1 hour. The crude mixture was then purified with reverse-phase column chromatography (0.1% formic acid in acetonitrile-water) to give **DAT389-2** (19.0 mg, 0.035 mmol, 59% yield). ^1^H NMR (400 MHz, CD_3_OD) δ 7.70-7.65 (m, 2H), 7.43-7.37 (m, 2H), 7.34 (dd, *J* = 8.0, 1.0 Hz, 1H), 7.27 (s, 1H), 6.92 (dd, *J* = 7.6, 7.6 Hz, 1H), 6.71 (dd, *J* = 7.6, 0.8 Hz, 1H), 4.50 (s, 2H), 4.08 (s, 2H), 3.74-3.66 (m, 4H), 3.63-3.57 (m, 2H), 3.57-3.52 (m, 4H), 3.30 (t, *J* = 5.1 Hz, 2H). ^13^C NMR (101 MHz, CD_3_OD) δ 173.09, 145.37, 139.50, 131.96, 128.78, 128.63, 1128.25, 123.24, 123.18, 120.95, 119.10, 116.78, 106.29, 72.05, 71.44, 71.42, 71.36, 71.23, 70.97, 51.68, 42.94. LC-MS, ESI^+^, *m/z* 551.2 [M+H]^+^.

##### 2-(2-(2-(2-Aminoethoxy)ethoxy)ethoxy)-N-(4-(N-(3-chloro-1H-indol-7-yl)sulfamoyl)benzyl)acetamide (DAT389-3)

To a solution of **DAT-389-2** (19.0 mg, 0.035 mmol) in MeOH (3.0 mL) was added 10% Pd/C (2.0 mg). The reaction mixture was degassed under reduced pressure and filled with H_2_ gas. After being stirred at room temperature for 2 hours, the mixture was filtered, and concentrated. The product was used in the next reaction without further purifications.

##### (S)-N-(4-(N-(3-Chloro-1H-indol-7-yl)sulfamoyl)benzyl)-2-(2-(2-(2-(2-(4-(4-chlorophenyl)-2,3,9-trimethyl-6H-thieno[3,2-f][1,2,4]triazolo[4,3-a][1,4]diazepin-6-yl)acetamido)ethoxy)ethoxy)ethoxy)acetamide (DAT389)

To a solution of the obtained crude **DAT389-3** (0.035 mmol), (S)-2-(4-(4-chlorophenyl)-2,3,9-trimethyl-6H-thieno[3,2-f][1,2,4]triazolo[4,3-a][1,4]diazepin-6-yl)acetic acid (14.0 mg, 0.035 mmol, 1.0 eq.), *N,N-*diisopropylethylamine (13.5 µl, 0.10 mmol, 3.0 eq.), and 1-hydroxy-7-azabenzotriazole (4.6 mg, 0.035 mmol, 1.0 eq.) in DMF (0.5 mL) was added HATU (13 mg, 0.035 mmol, 1.0 eq.) at room temperature. The reaction was allowed to stir for 1 hour. Then the mixture was purified with reverse-phase preparative HPLC chromatography (0.1% formic acid in acetonitrile-water) to give **DAT389** (10.1 mg, 0.011 mmol, 32% yield). ^1^H NMR (400 MHz, CD_3_OD) δ 7.58-7.57 (m, 2H), 7.46-7.38 (m, 4H), 7.33-7.29 (m, 3H), 7.25 (s, 1H), 6.89 (t, *J* = 7.7 Hz, 1H), 6.70 (dd, *J* = 7.53, 0.77 Hz, 1H), 4.66 (dd, *J* = 8.8, 5.5 Hz, 1H), 4.45 (s, 2H), 4.08 (s, 2H), 3.74-3.67 (m, 4H), 3.64-3.61 (m, 2H), 3.60-3.53 (m, 4H), 3.49-3.30 (m, 4H), 2.69 (s, 3H), 2.46 (s, 3H), 1.69 (s, 3H). ^13^C NMR (106 MHz, CD_3_OD) δ 173.9, 167.1, 157.9, 153.1, 146.2, 140.5, 139.0, 138.9, 134.4, 134.1, 132.92, 132.89, 132.8, 132.2, 130.7, 129.7, 129.4, 129.1, 124.3, 124.1, 121.9, 119.9, 117.5, 107.2, 72.9, 72.4, 72.3, 72.2, 72.1, 71.5, 56.1, 43.9, 41.4, 39.7, 15.3, 13.8, 12.5. LC-MS, ESI^+^, *m/z* 906.9 [M+H]^+^.

### 2 Biology

#### Plasmids and oligonucleotides

The design and construction of the human CRL-focused sgRNA library used for BRD4 stability screens, lentiviral sgRNA expression vectors used for single gene knockouts, as well as viral vectors used for the engineering of inducible Cas9 cell lines have been described previously^14,15^. For the engineering of the fluorescent protein stability reporters, BRD4(S) (Twist Bioscience), BRD2 (Addgene plasmid # 65376) or BRD3 (Addgene plasmid # 65377; both gifts from Kyle Miller^51^) were cloned into a pRRL lentiviral vector, fused to a 3xV5 tag and mTagBFP, and coupled to mCherry for normalization. For knockout/rescue studies, DCAF16 open reading frame cDNA (Twist Bioscience) was synonymously mutated to remove the sgRNA protospacer adjacent motif and seed sequence, coupled to a FLAG-Tag and cloned into a pRRL lentiviral vector expressing iRFP670 for flow cytometric detection. All plasmids and sgRNAs used in this study are shown in Extended Data Table 1, and the CRL-focused sgRNA libraries used for FACS-based and viability-based CRISPR/Cas9 screens, are shown in Supplementary Table 1 and Supplementary Table 3, respectively.

#### Cell culture

HEK293(T) and HCT-116 cell lines, originally sourced from ATCC, were provided by the MRC PPU reagents facility at the University of Dundee. HEK293, Lenti-X 293T lentiviral packaging cells (Clontech) and HCT-116 were cultured in DMEM (Gibco) supplemented with 10% fetal bovine serum (FBS; Thermo Fisher), 100 U/mL penicillin/streptomycin (Thermo Fisher) and 2 mM L-glutamine (Thermo Fisher). MV4;11 and KBM7 cells were cultured in IMDM (Gibco), supplemented with the same additives as above. All cell lines were grown in a humidified incubator at 37⁰C and 5% CO_2_ and routinely tested for mycoplasma contamination.

#### Lentivirus production and transduction

Semiconfluent Lenti-X cells were co-transfected with lentiviral plasmids, the lentiviral pCMVR8.74 helper (Addgene plasmid # 22036) and pMD2.G envelope (Addgene plasmid # 12259; both gifts from Didier Trono) plasmids using polyethylenimine (PEI) transfection (PEI MAX® MW 40,000, Polysciences) as previously described. Virus containing supernatant was clarified by centrifugation. Target cells were infected at limiting dilutions in the presence of 4 μg/mL of polybrene (Santa Cruz Biotechnology).

#### CRISPR/Cas9 DCAF15 KO cell line generation

DCAF15 KO cell line was generated using HCT-116 cells via ribonuclear protein (RNP) transfection using gRNAs targeting both exon 2 and exon 4 (IDT) (Extended Data Table 1), spCas9 Nuclease V3 (IDT) and TransIT-X2® (Mirus Bio). Following initial transfection for 48 hours, cells were trypsinized and re-plated in 96-well plates at low density and allowed to grow for > 2 weeks. Single colonies were then isolated and expanded, and later verified for DCAF15 KO via western blotting using an optimised RBM39 degradation assay as well as via genomic DNA sequencing.

#### CRISPR/Cas9 HiBiT and BromoTag knock-in cell line generation

HiBiT BRD2, BRD3 and BRD4 cell lines were generated via RNP transfection of (IDT), single-stranded DNA oligonucleotides as the ssODN donor templates (IDT), spCas9 (Sigma Aldrich) and target-specific gRNA (IDT) (Extended Data Table 1). HEK293 cells were resuspended in buffer R (Thermo Fisher), along with the RNP complex and ssODN template, which were then subsequently electroporated using a 10 µL neon electroporation cuvette tip (Thermo Fisher). Immediately following electroporation, cells were added to pre-warmed DMEM supplemented with 10% (v/v) FBS (100 U/mL penicillin/streptomycin additionally added for BromoTag cell lines only). Edited pools were analysed for HiBiT insertion by assaying for luminescence on a PHERAstar spectrophotometer (BMG Labtech) 48–72 h post-electroporation. Successful knock-in of HiBiT three days post-electroporation was first established using Promega’s HiBiT lytic assay on the mixed cell population. Following identification of luminescent signal these cells underwent single cell sorting using an SH800 cell sorter (Sony Biotechnology). Single cells were sorted into 3 × 96 well plates per experiment in 200 μL of 50% filtered preconditioned media from healthy cells and 50% fresh DMEM. After two weeks, all visible colonies were expanded, validated using Promega’s HiBiT lytic assay and subsequently frozen down.

BromoTag cell lines were generated in HEK293 cells via simultaneous transfection of two vectors at a 4:1 reagent:DNA ratio with FuGENE 6 (Promega). The first vector was a pMK-RQ vector containing 500 bp homology arms on either side of either an eGFP-IRES-BromoTag or eGFP-IRES-HiBiT-BromoTag sequence for integration into MCM4 and BRD4, respectively (Extended Data Table 1). The second vector was a custom pBABED vector harbouring a U6-sgRNA, Cas9 and puromycin expression cassettes. MRC-PPU CRISPR services constructed these plasmids at the University of Dundee. Following transfection, cells were repeatedly washed with PBS and then treated with 1 µg/mL puromycin for one week before FACS sorting. Single cell clones were generated by FACS sorting of single GFP^+^ cells using an SH800 cell sorter and sorting between 2-10 × 96 well plate in 200 μL of 50% filtered preconditioned media from healthy cells mixed with 50% fresh media.

#### siRNA-mediated knockdown

Cells were transfected for 48 hours using ON-TARGETplus SMARTPool siRNAs for DCAF15, DCAF16, DDB1, RBX1, CUL4A, and CUL4B (all from Dharmacon) and RNAiMAX (Invitrogen) following the manufacturer’s instructions, with 35 pmol of siRNA per well in 6-well plates. When simultaneously targeting 2 genes, half the amount of siRNA was used for each gene.

#### Cell viability assay

MV4;11, HCT-116 or KBM7 cells were plated in 96-well plates at a density of 0.5×10^6^ (MV4;11 and HCT-116) or 0.1×10^6^ (KBM7) cells/mL in 50 µL cell suspension per well. The following day, 2x stocks of compounds were added for a final volume of 100 µL. Cells were treated for 24 (MV4;11), 72 (KBM7) or 96 (HCT-116) hours in a humidified incubator at 37⁰C and 5% CO_2_. CellTiter-Glo (G7570, Promega) or CellTiter-Glo® 2.0 reagent (G924A, Promega) was added to the plates per manufacturer instructions, before shaking the plate for 3-20 minutes at 300 rpm and measuring the luminescence using a PHERAstar (BMG Labtech) or VICTOR X3 (Perkin Elmer) multilabel plate reader. The results were normalised to DMSO controls and analysed using Graphpad Prism (v9.3.1 or v9.5.0) to derive EC_50_ values by 4-parameter non-linear regression curve fitting or interpolation of a sigmoidal standard curve.

#### Degradation assays and western blotting

HEK293 and HCT-116 cells were plated in 6-well plates at varying densities (0.2-0.6×10^6^ cells/mL) depending on experimental set up. In all experiments, media was changed prior to compound treatment. Stock solutions of compounds were prepared in DMSO at a concentration of 10 mM and stored at −20 ⁰C. Working dilutions were made fresh using DMEM media and added dropwise to 6-well plates. For competition assays, cells were plated in 6-well plates and pre-treated with 10 µM of the competition compounds, 3 µM MLN4924 or 50 µM MG132 for 1 hour, before treating with IBG1 at 10 nM for 2 hours.

For cell harvests, cells were washed once with ice-cold PBS before lysis for 15 minutes on ice with RIPA buffer supplemented with Benzonase (1:1000, Sigma or Millipore no. 70746,) and cOmplete™ EDTA-free Protease Inhibitor Cocktail (11873580001, Roche). Following clearance via centrifugation, protein concentration of lysates was determined using the Pierce™ BCA Protein Assay (23225, Fisher Scientific) and typically 20-30 µg of lysate was prepared using 4x LDS sample buffer (Thermo Fisher) and 10% 2-Mercaptoethanol or 50 mM dithiothreitol (DTT) and run on NuPAGE 4-12% bis-tris gels (Thermo Fisher). Proteins were transferred to nitrocellulose membranes, blocked for 1 hour in 5% milk TBS-T at room temperature, before incubating with primary antibodies overnight at 4 °C. The following primary antibodies were used: BRD2 (no. Ab139690, Abcam), BRD3 (no. Ab50818, Abcam), BRD4 (E2A7X, no. 13440, Cell Signaling Technology and no. Ab128874, Abcam), BromoTag (no. NBP3-17999, Novus Biologicals), CUL4A (no. A300-738A, Bethyl Laboratories), CUL4B (no. 12916-1-AP, Proteintech), DDB1 (no. A300-462A, Bethyl Laboratories), MCM4 (no. ab4459, Abcam) RBM39 (no. HPA001591, Atlas Antibodies), RBX1 (D3J5I, no. 11922, Cell Signalling Technology), DCAF11 (no. A15519, ABclonal), cleaved Caspase-3 (D3E9, no. 9579, Cell Signalling Technology), PARP1 (no. 9542, Cell Signalling Technology), MYC (D84C12, no. 5605, Cell Signalling Technology), β-Actin (AC-15, no. A5441, Sigma-Aldrich), α-Tubulin (DM1A, no. T9026, Sigma-Aldrich). Membranes were then washed in TBS-T and incubated with fluorescent or HRP-conjugated secondary antibodies for 1 hour at room temperature, before further washes and imaging on a ChemiDoc Touch imaging system (Bio-Rad). Secondary antibodies used were HRP anti-rabbit IgG (7074, Cell Signaling Technology), HRP anti-mouse IgG (7076, Cell Signaling Technology), IRDye® 680RD anti-mouse (no. 926-68070, Li-Cor), IRDye® 800CW anti-rabbit (no. 926-32211, Li-Cor), StarBright™ blue 520 goat anti-mouse (no. 12005866, Biorad) and hFABTM rhodamine anti-tubulin (no. 12004165, Biorad).

#### HiBiT degradation assays

Endogenously tagged HiBiT cells were plated in 96-well plates (PerkinElmer) at a density of 0.5×10^6^ cells/mL, with 50 µL of cell suspension per well. The following day, 2x stocks of compounds were added for a final volume of 100 µL. Cells were treated for 5, 6 or 24 hours as indicated in the respective figure legends before lysis using the HiBiT lytic assay buffer (Promega) per manufacturer instructions. Plates were then read on a BMG Pherastar® plate reader for luminescence detection. Treated wells were normalised to a DMSO-only control and analysed using GraphPad Prism 9 via fitting of non-linear regression curves for extraction of DC_50_ and D_MAX_ values.

#### Kinetic ubiquitination and degradation assays

For kinetic ubiquitination assays, HiBiT-tagged 293 cells were seeded in 6-well plates at a density of 8×10^6^ cells/mL in 2 mL volume. After 5 hours, LgBiT and Halo-Ub cDNA (Promega) were transfected using FuGENE HD (Promega) with 1 µg of each plasmid at a 3:1 transfection reagent:plasmid ratio. The following day, cells were trypsinized and re-suspended in phenol-red free OptiMEM (Gibco) supplemented with 4% FBS and seeded in 96-well plates at a density of 3.5×10^5^ cells/mL in the presence or absence of 0.1 mM HaloTag NanoBRET™ ligand (Promega). Following overnight incubation, media was removed from the wells and replaced with 90 µL OptiMEM (4% FBS) with a 1:100 dilution of Vivazine substrate. The plates were incubated at 37 °C for 1 hour before 10X stocks of experimental compounds were added and the plates were analyzed on a GloMAX® Discover microplate reader (Promega) in kinetic mode for NanoBRET ratio metric (460 nm donor and 618 nm acceptor emissions) signal detection for 6 hours, with measurements taken every 3-5 minutes. Data was processed by subtracting NanoBRET ligand-free controls before plotting NanoBRET signal versus time in GraphPad Prism 9.

Kinetic degradation assays were performed as previously described^36^, using the HiBiT-tagged cells with exogenous LgBiT transfection as described above for the kinetic ubiquitination assays. Cells were incubated in Endurazine substrate (1:100) for 2.5 hours at 37 °C prior to 10X compound addition, with luminescence measurements taken on a GloMAX® Discover microplate reader (Promega) every 15 minutes for 24 hours. Data were normalised to DMSO-only controls and plotted for luminescence signal versus time in GraphPad Prism 9.

#### NanoBRET bromodomain confirmational sensor assay

Transient transfection of the dual NanoLuc and Halo-Tagged tagged BRD4^Tandem^ plasmid (Promega) was performed as described previously^52^. Briefly, 0.02 µg of plasmid and 2 µg of carrier DNA were combined with FuGENE HD (Promega) at a 3:1 ratio and added per well of a 6-well plate seeded with 70% confluent HEK293 cells. The following day, cells were trypsinized and re-suspended in phenol-red free OptiMEM (Gibco) supplemented with 4% FBS and 100 µL were seeded per well in 96-well plates at a density of 2×10^5^ cells/mL in the presence or absence of 0.1 mM HaloTag NanoBRET™ ligand (Promega). The following morning, the media was aspirated and replaced with phenol red-free media containing MG132 (10 µM final concentration) for 1 hour, before cells were incubated with test compounds for 3 hours. For cell lysis and detection, 100 µL of 2x NanoBRET substrate solution was added per well, the plate was incubated in darkness while shaking at 400 RPM for 3 minutes, before reading on a BMG Pherastar® plate reader equipped with a NanoBRET filter (618/460 nm). Wells lacking Halo ligand were subtracted from wells containing Halo ligand, and the fold increase in signal compared to DMSO was plotted using GraphPad Prism 9.

#### FACS-based CRISPR/Cas9 BRD4 stability screens

For pooled FACS-based CRISPR–Cas9 BRD4 protein stability screens, a CRL-focused sgRNA library^19^ was lentivirally packaged using polyethylenimine (PEI MAX® MW 40,000, Polysciences) transfection of Lenti-X cells and the lentiviral pCMVR8.74 helper (Addgene plasmid # 22036) and pMD2.G envelope (Addgene plasmid # 12259; both gifts from Didier Trono) plasmids. The virus containing supernatant was cleared of cellular debris by filtration through a 0.45-µm PES filter and used to transduce KBM7 BRD4-BFP reporter cells harbouring a doxycycline inducible Cas9 allele (KBM7 iCas9) at a multiplicity of infection (MOI) of 0.05 and 1,000-fold library representation. Library-transduced cells were selected with G418 (1 mg/mL, Gibco) for 14 days, expanded and Cas9 expression was induced with DOX (0.4 µg/mL, PanReac AppliChem).

3 days after Cas9 induction, 25 million cells per condition were treated with DMSO (1:1000), MZ1 (10 nM), IBG1 (1 nM), GNE-0011 (1 µM), IBG3 (0.1 nM) or IBG4 (100 nM) for 6 hours in two biological replicates. Cells were washed with PBS, stained with Zombie NIR™ Fixable Viability Dye (1:1000, BioLegend) and APC anti-mouse CD90.1/Thy-1.1 antibody (1:400, BioLegend) in the presence of Human TruStain FcX™ Fc Receptor Blocking Solution (1:400, BioLegend), and fixed with 0.5 mL methanol-free paraformaldehyde 4% (Thermo Scientific™ Pierce™) for 30 min at 4 °C, while protected from light. Cells were washed with and stored in FACS buffer (PBS containing 5% FBS and 1 mM EDTA) at 4 °C over night. The next day, cells were strained trough a 35 µm nylon mesh and sorted on a BD FACSAria™ Fusion (BD Biosciences) using a 70 µm nozzle. Aggregates, dead (ZombieNIR positive), Cas9-negative (GFP) and sgRNA library-negative (Thy1.1-APC) cells were excluded, and the remaining cells were sorted based on their BRD4-BFP and mCherry levels into BRD4^HIGH^ (5-10% of cells), BRD4^MID^ (25-30%) and BRD4^LOW^ (5-10%) fractions. For each sample, cells corresponding to at least 1,500-fold library representation were sorted per replicate.

Next-generation sequencing (NGS) libraries of sorted cell fractions were prepared as previously described^14^. In brief, genomic DNA was isolated by cell lysis (10 mM Tris-HCl, 150 mM NaCl, 10 mM EDTA, 0.1% SDS), proteinase K treatment (New England Biolabs) and DNAse-free RNAse digest (Thermo Fisher Scientific), followed by two rounds of phenol extraction and 2-propanol precipitation. Isolated genomic DNA was subjected to several freeze–thaw cycles before nested polymerase chain reaction (PCR) amplification of the sgRNA cassette.

Barcoded NGS libraries for each sorted population were generated using a two-step PCR protocol using AmpliTaq Gold (Invitrogen. The resulting PCR products were purified using Mag-Bind® TotalPure NGS beads (Omega Bio-tek) and amplified in a second PCR introducing the standard Illumina adapters. The final Illumina libraries were bead-purified, pooled and sequenced on HiSeq 3500 or NovaSeq 6000 platforms (Illumina).

Screen analysis was performed as previously described^14^. Briefly, sequencing reads were quantified using the crispr-process-nf Nextflow workflow, available at https://github.com/ZuberLab/crispr-process-nf/tree/566f6d46bbcc2a3f49f51bbc96b9820f408ec4a3. For statistical analysis, we used the crispr-mageck-nf Nextflow workflow, available at https://github.com/ZuberLab/crispr-mageck-nf/tree/c75a90f670698bfa78bfd8be786d6e5d6d4fc455. To calculate gene-level enrichment, the sorted populations (BRD4^HIGH^ or BRD4^LOW^) were compared to the BRD4^MID^ populations in MAGeCK (0.5.9)^53^, using median normalized read counts.

#### Viability-based CRISPR/Cas9 screen

The ubiquitin/Nedd8 system CRISPR-KO library (Supplementary Table 3) was generated using the covalently-closed-circular-synthesized (3Cs) technology, as previously described^54,55^. The library contained 3,347 gRNAs cloned under the U6 promoter in a modified pLentiCRISPRv2-puromycin vector containing a modified gRNA scaffold sequence starting with GTTTG. Each gene was represented by 4 gRNAs selected with the Broad Institute CRISPick tool^56–58^. Additionally, the library included a set of essential genes, non-targeting as well as AAVS1-targeting control sgRNAs.

HCT-116 cells were transduced with the ubiquitin/Nedd8 system lentiviral CRISPR-Cas9 library at an MOI 0.5 and a coverage of 500. Cells were selected with 1 μg/mL puromycin for 12 days. Eight million selected cells per condition were then plated in T175 flasks. Cells were treated with DMSO or IBG1 (58 nM), corresponding to 4 times the IC_50_ value for 3 days, followed by replating and treatment for additional 3 days. After a total of 6 days of treatment, cells were trypsinized, washed three times with PBS, followed by genomic DNA isolation. Sequencing libraries were prepared via PCR as previously described^55^ and purified via GeneJET Gel Extraction Kit (Thermo Fisher Scientific).

Raw sequencing data were demultiplexed with bcl2fastq v2.20.0.422 (Illumina) to generate raw fastq files. To determine the abundance of individual gRNAs per samples, the fastq files were trimmed using cutadapt 2.8 to retain only the putative gRNA sequences. These sequences were then aligned to the original gRNA library with Bowtie 2.3.0 and only perfect matches were counted. Statistical analysis was performed via MAGeCK^53^, using median or total read count normalization and removal of gRNAs with zero counts in the control samples. Genes with a LFC > 1 or < −1 and a p-value < 0.01 were labelled as significantly depleted or enriched hits.

#### Flow-cytometric BRD4 reporter assay

KBM7 iCas9 cells were transduced with lentivirus expressing WT, mutated or truncated versions of SFFV-BRD4(S)-mTagBFP-P2A-mCherry to generate stable reporter cell lines. For evaluation of reporter degradation, cells were treated with DMSO (1:1000), IBG1 (1 nM), dBET6 (10 nM), IBG3 (0.1 nM) or IBG4 (100 nM) for 6 hours before flow cytometry analysis on an LSRFortessa (BD Biosciences).

To quantify the influence of genetic perturbation on compound-induced reporter degradation, stable BRD4(S) or BRD4^Tandem^ reporter cell lines were transduced with a lentiviral sgRNA (pLenti-U6-sgRNA-IT-EF1αs-Thy1.1-P2A-NeoR) and/or transgene expression vector (pRRL-SFFV-3xFLAG-DCAF16-EF1αs-iRFP670) to 30-50% transduction efficiency. Cas9 expression was induced with doxycycline (0.4 µg/mL) for 3 days, followed by 6 hours of degrader treatment. Cells were stained for sgRNA expression with an APC conjugated anti-mouse CD90.1/Thy1.1 antibody (no. 202526, BioLegend; 1:400) and Human TruStain FcX Fc receptor blocking solution (no. 422302, BioLegend; 1:400) for 5 minutes in FACS buffer (PBS containing 5% FBS and 1 mM EDTA) at 4 °C. Cells were washed and resuspended in FACS buffer and analysed on an LSRFortessa (BD Biosciences).

Flow cytometric data analysis was performed in FlowJo v10.8.1. BFP and mCherry mean fluorescence intensity (MFI) values for were normalized by background subtraction of the respective values from reporter-negative KBM7 cells. BRD4 abundance was calculated as the ratio of background subtracted BFP to mCherry MFI, and is displayed normalized to DMSO treated, sgRNA/cDNA double negative cells.

#### Quantitative proteomics

For unbiased identification of degrader target proteins, 50×10^6^ KBM7 iCas9 cells per condition were treated with DMSO (1:1000), IBG1 (1 nM) or dBET6 (10 nM) for 6 hours in biological triplicates. Cells were harvested via centrifugation, washed three times in ice-cold PBS and snap-frozen in liquid nitrogen. Cell pellet were lysed in 500 µL of freshly prepared lysis buffer (50 mM HEPES pH 8.0, 2% SDS, 1 mM PMSF and protease inhibitor cocktail (Sigma-Aldrich)). Samples incubated at RT for 20 minutes before heating to 99 °C for 5 min. DNA was sheared by sonication using a Covaris S2 high performance ultrasonicator. Cell debris was removed by centrifugation at 16,000 × g for 15 min at 20 °C. Supernatant was transferred to fresh tubes and protein concentration determined using the BCA protein assay kit (Pierce Biotechnology). Filter-aided sample preparation (FASP) was performed using a 30 kDa molecular weight cutoff centrifugal filters (Microcon 30, Ultracel YM-30, Merck Millipore) as previously described^59^. In brief, 200 µg of total protein per sample was reduced by the addition of DTT to a final concentration of 83.3 mM, followed by incubation at 99 °C for 5 minutes. Samples were mixed with 200 μL freshly prepared 8 M urea in 100 mM Tris-HCl (pH 8.5) (UA-solution) in the filter unit and centrifuged at 14.000 × g for 15 min at 20 °C to remove SDS. Residual SDS was washed out by a second wash step with 200 μL UA. Proteins were alkylated with 100 µL of 50 mM iodoacetamide in the dark for 30 min at RT. Thereafter, three washes were performed with 100 μL of UA solution, followed by three washes with 100 μL of 50 mM TEAB buffer (Sigma-Aldrich). Proteolytic digestion was performed using trypsin (1:50) overnight at 37 °C. Peptides were recovered using 40 μL of 50 mM TEAB buffer followed by 50 μL of 0.5 M NaCl. Peptides were desalted using the Pierce™ Peptide Desalting Spin Columns (Thermo Scientific). TMTpro 16plex Label Reagent Set was used for labeling according to the manufacturer (Pierce,). After the labeling reaction was quenched, the samples were pooled, the organic solvent removed in a vacuum concentrator, and the labeled peptides purified by C18 solid phase extraction (SPE).

For offline fractionation via reverse phase (RP) HPLC at high pH as previously described, tryptic peptides were re-buffered in 10 mM ammonium formate buffer (pH 10). Peptides were separated into 96 time-based fractions on a Phenomenex C18 RP column (150 × 2.0 mm Gemini-NX, 3 µm C18 110Å, Phenomenex) using an Agilent 1200 series HPLC system fitted with a binary pump delivering solvent at 50 µL/min. Acidified fractions were consolidated into 36 fractions via a concatenated strategy as previously described^60^. After removal of solvent in a vacuum concentrator, samples were reconstituted in 0.1% TFA prior to LC-MS/MS analysis.

Mass spectrometry analysis was performed on an Orbitrap Fusion Lumos Tribrid mass spectrometer coupled to a Dionex Ultimate 3000 RSLCnano system (via a Nanospray Flex Ion Source (all Thermo Fisher Scientific) interface and operated via Xcalibur Version 4.3.73.11 and Tune 3.4.3072.18. Peptides were loaded onto a trap column (PepMap 100 C18, 5 μm, 5 × 0.3 mm, Thermo Fisher Scientific) at a flow rate of 10 μL/min using 0.1% TFA as loading buffer. After loading, the trap column was switched in-line with an Acclaim PepMap nanoHPLC C18 analytical column (2.0 µm particle size, 75µm IDx500mm, # 164942, Thermo Fisher Scientific). The column temperature was maintained at 50 °C. Mobile phase A consisted of 0.4% formic acid (FA) in water, and mobile phase B consisted of 0.4% FA in a mixture of 90% acetonitrile and 10% water. Separation was achieved using a four-step gradient over 90 min at a flow rate of 230 nL/min. In the liquid junction setup, electrospray ionization was enabled by applying a voltage of 1.8 kV directly to the liquid being sprayed, and non-coated silica emitter was used. The mass spectrometer was operated in a data dependent acquisition (DDA) mode using a maximum of 20 dependent scans per cycle. Full MS1 scans were acquired in the Orbitrap with a scan range of 400 - 1600 m/z and a resolution of 120,000 at 200m/z. Automatic gain control (AGC) was set to ‘standard’ and a maximum injection time (IT) of 50 ms was applied. MS2 spectra were acquired in the Orbitrap at a resolution of 50,000 at 200 m/z with a fixed first mass of 100 m/z. To achieve maximum proteome coverage, a classical tandem MS approach was chosen instead of the available synchronous precursor selection (SPS)-MS3 approach. To minimize TMT ratio compression effects by interference of contaminating co-eluting isobaric peptide ion species, precursor isolation width in the quadrupole was set to 0.5 Da and an extended fractionation scheme applied. Monoisotopic peak determination was set to ‘peptides’ with inclusion of charge states between 2 and 5. Intensity threshold for MS2 selection was set to 2.5×10^4^. Higher energy collision induced dissociation (HCD) was applied with a normalized collision energy (NCE) of 34%. Normalized AGC was set to 200% with a maximum injection time of 86 ms. Dynamic exclusion for selected ions was 90 s.

The acquired raw data files were processed using Proteome Discoverer v.2.4.1.15, via TMT16plex quantification method. Sequest HT database search engine and the Percolator validation software node were used to remove false positives with FDR 1% at the peptide and protein level. All MS/MS spectra were searched against the human proteome (Canonical, reviewed, 20 304 sequences) and appended known contaminants and streptavidin, with a maximum of two allowable miscleavage sites. The search was performed with full tryptic digestion with or without deamidation on amino acids asparagine, glutamine, and arginine. Methionine oxidation and protein N-terminal acetylation, as well as methionine loss and protein N-terminal acetylation with methionine loss were set as variable modifications, while carbamidomethylation of cysteine residues and tandem mass tag (TMT) 16-plex labeling of peptide N termini and lysine residues were set as fixed modifications. Data were searched with mass tolerances of ±10 ppm and ±0.025 Da for the precursor and fragment ions, respectively. Results were filtered to include peptide spectrum matches with Sequest HT cross-correlation factor (Xcorr) scores of ≥1 and high peptide confidence assigned by Percolator. MS2 signal-to-noise (S/N) values of TMTpro reporter ions were used to calculate peptide/protein abundance values. Peptide spectrum matches with precursor isolation interference values of ≥70% and average TMTpro reporter ion S/N ≤10 were excluded from quantification. Both unique and razor peptides were used for TMT quantification. Correction of isotopic impurities was applied.

Data were normalized to total peptide abundance and scaled “to all average”. Abundances were compared to DMSO treated cells and protein ratios were calculated from the grouped protein abundances using an ANOVA hypothesis test. Adjusted p-values were calculated using the Benjamini-Hochberg method. Proteins with less than three unique peptides detected were excluded from downstream analysis.

#### Protein construction, expression and purification

His_6_-TEV-BRD4 bromodomain 1 (BRD4^BD^^1^) (amino acids 44-178) and His_6_-TEV-BRD4 bromodomain 2 (BRD4^BD2^) (amino acids 333-460) were expressed in *Escherichia coli* (*E. Coli)* BL21(DE3) and purified as described previously^61^. Briefly, proteins were purified by nickel affinity chromatography and SEC. His_6_ tag cleavage and reverse nickel affinity was performed prior to SEC for some applications, for others the tag was left on. Purified proteins in 20 mM HEPES, 150 mM sodium chloride, 1 mM DTT, pH 7.5 were aliquoted and flash frozen in liquid nitrogen and stored at −80 °C.

His_6_-SUMO-TEV-BRD4^Tandem^ (residues 1-463) was prepared as previously described^36^. Briefly, protein was expressed in *E. coli* BL21(DE3) and purified sequentially by nickel affinity on a HisTrap HP 5 mL column (Cytiva), His_6_ tag cleavage by SENP1 followed by reverse nickel affinity, cation exchange on a HiTrap SP HP 5 mL column (Cytiva), and size exclusion on a HiLoad 16/600 Superdex 200 pg column (Cytiva). Purified protein in 20 mM HEPES, 100 mM sodium chloride, 1 mM TCEP, pH 7.5 was aliquoted and flash frozen in liquid nitrogen then stored at −80 °C.

BRD4^Tandem^ (residues 43-459) was cloned into pRSF-DUET or a modified pGEX4T1 with an N-terminal His_10_ tag and HRV3C cleavage site or a His_12_-GST tag and TEV cleavage site, respectively.

His_10_-3C-BRD4^Tandem^ (residues 43-459) was transformed into *E. coli* BL21(DE3) and overnight expression at 18 °C was induced with 0.35 mM IPTG at OD_600_ ∼0.8-1. Cells were harvested by centrifugation and pellets were resuspended in ice cold PBS then spun down again. Supernatant was removed and pellets were flash frozen in liquid nitrogen and stored at −80 °C. Cells were thawed and resuspended in lysis buffer (50 mM HEPES, 500 mM NaCl, 0.5 mM TCEP, pH 7.5) supplemented with 2 mM magnesium chloride, DNAse and cOmplete EDTA-free Protease Inhibitor Cocktail (Roche, 1 tablet/L initial culture volume) and lysed at 35,000 psi using a CF1 Cell Disruptor (Constant Systems). The lysate was cleared by centrifugation at 20,000 rpm for 30 min at 4 °C then syringe filtered using a 0.45 μm filter. The lysate was supplemented with 40 mM imidazole and loaded on to a 5 mL HisTrap HP column (Cytiva) equilibrated in lysis buffer with 40 mM imidazole, washed at 60 mM imidazole and eluted with a gradient up to 100% elution buffer (50 mM HEPES, 500 mM NaCl, 0.5 mM TCEP, 500 mM imidazole, pH 7.5). The prep was split as required for tag cleavage or for purification of the His_10_-3C-taged form. For tag cleavage, the sample was buffer exchanged into lysis buffer on a HiPrep 26/10 Desalting column and HRV3C protease was added to cleave the tag overnight at 4°C. Imidazole was added to 20 mM to the cleaved BRD4^Tandem^ and the sample was run on a 5 mL HisTrap HP column equilibrated in lysis buffer with 20 mM imidazole and washed with the same imidazole concentration. The flowthrough and wash containing BRD4^Tandem^ were pooled and, along with uncleaved His_10_-3C-BRD4^Tandem^, were concentrated in 10,000 MWCO Amicon centrifugal filter units (Merck Millipore). The proteins were each loaded separately onto a HiLoad 26/600 Superdex 200 pg column (GE LifeSciences) equilibrated in 20 mM HEPES, 150 mM NaCl, 0.5 mM TCEP, pH 7.5. Fractions containing either pure BRD4^Tandem^ or His_10_-3C-BRD4^Tandem^ were confirmed by SDS-PAGE, then pooled, concentrated and aliquoted for storage at −80 °C until use.

For use in Cryo-EM with DCAF16 and **1**, His_1_-GST-TEV-BRD4^Tandem^ (residues 43-459) expression in *E. coli* BL21(DE3) cells was induced at OD_600_ = 2 with 0.5 mM IPTG at 20 °C for 16 hours. Cells were harvested by centrifugation and resuspended in lysis buffer (50 mM HEPES, 500 mM NaCl, 20 mM Imidazole, 0.5 mM TCEP, pH 7.5) (10 mL/g pellet weight) supplemented with DNAse and 1 cOmplete EDTA-free Protease Inhibitor Cocktail tablet (Roche) per 2 L of culture. Cells were lysed at 30 kpsi using a CF1 Cell Disruptor (Constant Systems Ltd) and lysate was clarified by centrifugation. Lysate was filtered through a BioPrepNylon Matrix Filter (BioDesign) then incubated with 1 mL Ni-NTA resin per litre culture for 1 hour. The lysate-resin slurry was poured into a Bio-Rad Econo-column and resin was washed with >10 CV lysis buffer. Bound protein was eluted with elution buffer (50 mM HEPES pH 7.5, 150 mM NaCl, 500 mM Imidazole, 0.5 mM TCEP) then incubated with 1 mL glutathione agarose resin per litre culture for 30 mins. The mixture was poured into an Econo-column and resin was washed with 20 mM HEPES, 150 mM NaCl, 0.5 mM TCEP, pH 7.5. TEV protease was added to the resin slurry for on-bead cleavage and the column was incubated overnight on a roller at 4 °C. Protein was eluted from the column then concentrated and run on a HiLoad 16/600 Superdex 75 pg column equilibrated in 20 mM HEPES, 150 mM NaCl, 0.5 mM TCEP, pH 7.5. Fractions containing protein were pooled, concentrated and aliquoted then flash frozen in liquid nitrogen then stored at −80 °C until use.

DCAF15Δpro with N-terminal His_6_-TEV-Avi tag, DDB1ΔBPB (residues 396-705 replaced with a GNGNSG linker), and full-length DDA1 coding sequences were cloned into a pFastBacDual vector. Bacmid was generated using the Bac-to-Bac baculovirus expression system (Thermo Fisher Scientific). Baculovirus was generated via an adapted single step protocol^62,63^. Briefly, bacmid (1 µg/mL culture volume) was mixed with 2 µg PEI 25K (Polysciences, Inc.) per µg bacmid in 200 µL warm PBS and incubated at RT for 30 mins. The mixture was added to a suspension culture of Sf9 cells at 1 x 10^6^ cells/mL in Sf-900™ II SFM (Gibco) and incubated at 27 °C with shaking at 110 rpm. Viral supernatant (P0) was harvested after 4-6 days. For expression, *Spodoptera* frugiperda cells (Sf9) were grown to densities between 1.9-3.0 x 10^6^ cells/mL in Sf-900™ II SFM (Gibco) and infected with a total virus volume of 1% per 1 x 10^6^ cells/mL. Cells were incubated at 27 °C in 2 L Erlenmeyer flasks (∼500 mL culture/flask) with shaking at 110 rpm for 48 hours. Cells were spun at 1,000 x g for 10 mins and supernatant was discarded. Pellets were resuspended in lysis buffer (50 mM HEPES, 200 mM NaCl, 2 mM TCEP, pH 7.5) with magnesium chloride (to 2 mM), benzonase (to 1 µg/mL) and cOmplete EDTA-free Protease Inhibitor Cocktail (Roche, 2 tablets/L initial culture volume). The suspension was frozen and stored at −80 °C, and then thawed. Cell suspensions were sonicated and lysates were centrifuged at 40,000 rpm for 30 mins. The supernatant was incubated with 1.5 mL Ni-NTA agarose resin (Qiagen) on a roller at 4 °C for 1.5 hour. The lysate-resin slurry was loaded into a glass bench top column. Supernatant was allowed to flow through then the resin was washed with wash buffer (50 mM HEPES, 200 mM NaCl, 2 mM TCEP, 20 mM imidazole, pH 7.5). Bound protein was eluted with elution buffer (50 mM HEPES pH 7.5, 200 mM NaCl, 2 mM TCEP, 500 mM imidazole). TEV protease was added to protein and dialyzed with buffer (50 mM HEPES, 200 mM NaCl, 2 mM TCEP, pH 7.5). Cleaved protein was run over 1.5 mL Ni-NTA agarose resin and the flow through and washes with binding buffer were collected and pooled. Protein was diluted with buffer (25 mM HEPES, 2 mM TCEP, pH 7.5) to adjust the NaCl concentration to 50 mM, then loaded onto a HiTrap Q HP 5 mL column (Cytiva). The column was washed with IEX buffer A and bound protein was eluted with a 0-100% IEX buffer B (25 mM HEPES, 1 M NaCl, 2 mM TCEP, pH 7.5) gradient. Fractions containing protein were pooled and concentrated to ∼1-2 mL then run on 16/600 Superdex 200 pg column in GF buffer (25 mM HEPES, 300 mM NaCl, 1 mM TCEP, pH 7.5). Fractions containing the purified protein complex were pooled, concentrated and aliquoted then flash frozen in liquid nitrogen for storage at −80 °C.

The coding sequence for full-length DCAF16 or DCAF11 with TEV-cleavable N-terminal His_6_-tags were cloned into a pFastBacDual vector under the control of the polh promoter. Coding sequences for full-length DDB1 or DDB1ΔBPB (residues 396-705 replaced with a GNGNSG linker) and full-length DDA1 were cloned into a pFastBacDual vector under the control of polh and p10 promoters, respectively. Bacmid was generated using the Bac-to-Bac baculovirus expression system (Thermo Fisher Scientific). Baculovirus was generated as described above and viral supernatant (P0) was harvested after 5-7 days. For expression, *Trichoplusia ni* High Five cells were grown to densities between 1.5-2 x 10^6^ cells/mL in Express Five™ SFM (Gibco) supplemented with 18 mM L-glutamine and infected with a total virus volume of 1% per 1 x 10^6^ cells/mL, consisting of equal volumes of DCAF16/DCAF11 and DDB1+DDA1 baculoviruses. Cells were incubated at 27 °C in 2 L Erlenmeyer flasks (∼600-650 mL culture/flask) with shaking at 110 rpm for 72 hours. Cells were spun at 1,000 x g for 20 mins and supernatant was discarded. Pellets were resuspended in 25 mL Binding Buffer (50 mM HEPES, 500 mM NaCl, 1 mM TCEP, pH 7.5), flash frozen in liquid nitrogen and stored at −80 °C. Pellets were thawed and diluted with Binding Buffer to ∼100 mL/L original culture volume. Tween-20 (to 1% (v/v)), magnesium chloride (to 2 mM), benzonase (to 1 µg/mL) and cOmplete EDTA-free Protease Inhibitor Cocktail (Roche, 2 tablets/L initial culture volume) were added to the cell suspension and stirred at RT for 30 mins. Cell suspensions were sonicated, and lysates were centrifuged at 23,000 rpm for 60 mins. Supernatants were filtered through 0.45 µm filters and supplemented with 10 mM imidazole then incubated with 2 mL cobalt agarose resin/L culture on a roller at 4 °C for 1 hour. The lysate-resin slurry was loaded into a glass bench top column. Supernatant was allowed to flow through then the resin was washed with wash buffer (50 mM HEPES, 500 mM NaCl, 1 mM TCEP, 15 mM imidazole, pH 7.5). Bound protein was eluted with elution buffer (50 mM HEPES, 500 mM NaCl, 1 mM TCEP, 250 mM imidazole, pH 7.5) and buffer exchanged on a 26/10 HiPrep Desalting column (Cytiva) into Binding Buffer. TEV protease was added to protein and incubated for 2 hours at RT then 4 °C overnight. Imidazole was added to the cleaved protein to a concentration of 10 mM and the sample was run over cobalt agarose resin. Flowthrough and washes with binding buffer supplemented with 10 mM imidazole were collected and pooled. Protein was buffer exchanged into ion exchange (IEX) buffer A (50 mM HEPES, 50 mM NaCl, 1 mM TCEP, pH 7.5) on a 26/10 HiPrep Desalting column then loaded onto a HiTrap Q HP 5 mL column (Cytiva). The column was washed with IEX buffer A and bound protein was eluted with a 0-100% IEX buffer B (50 mM HEPES, 1 M NaCl, 1 mM TCEP, pH 7.5) gradient. Fractions containing protein were pooled and concentrated then run on 16/600 Superdex 200 pg column in equilibrated in 20 mM HEPES, 150 mM NaCl, 1 mM TCEP, pH 7.5. Fractions containing the purified protein complex were pooled and concentrated then aliquoted and flash frozen in liquid nitrogen for storage at −80 °C.

#### Sulfo-Cy5 NHS ester labelling

For DCAF16 labelling, sulfo-Cy5 NHS ester (Lumiprobe) in DMF was prepared to a final concentration of 800 µM with DCAF16:DDB1ΔBPB:DDA1 (100 µM) and sodium bicarbonate (100 mM). For DCAF11 labelling, sulfo-Cy5 NHS ester (Lumiprobe) in DMF was prepared to a final concentration of 1 mg/mL with DCAF11:DDB1ΔBPB:DDA1 (1 mg/mL) and sodium bicarbonate (100 mM). The solutions were protected from light and shaken for 1 hour at RT. The solutions were spun down at 15,000 x g for 5 mins then run on a Superdex 200 10/300 GL column (Cytiva) to remove free dye and aggregated protein. Fractions containing the sulfo-Cy5-labelled protein were pooled and concentrated, the degree of labelling (DOL) was calculated to be greater than 100% for each batch of labelled protein. Labelled protein was aliquoted then flash frozen in liquid nitrogen and stored at −80 °C.

#### Fluorescence polarisation (FP) assay

Stock solutions of reaction components including DCAF15Δpro:DDB1ΔBPB:DDA1, DCAF16:-DB1ΔBPB:DDA1, His_6_-BRD4^BD^^1^, His_6_-BRD4^BD^^2^, BRD4^Tandem^ (residues 43-459), and FITC-sulfonamide probe^7^ were prepared in FP assay buffer (25 mM HEPES pH 7.5, 300 mM NaCl, 1.0 mM TCEP). DCAF15Δpro:DDB1ΔBPB:DDA1, DCAF16:DDB1ΔBPB:DDA1, BRD4^BD1^, BRD4^BD2^, and BRD4^Tandem^ were titrated 1/3 in FP assay buffer. Components were added to Corning™ 384-Well Solid Black Polystyrene Microplates to a final volume of 15 µL. Final concentration of 20 nM for FITC-sulfonamide probe was used while DCAF15Δpro:DDB1ΔBPB:DDA1, DCAF16:DDB1ΔBPB:DDA1, BRD4^BD^^1^, His_6_-BRD4^BD^^2^, and BRD4^Tandem^ were titrated from 4 µM to 5.5 nM. Background subtraction was performed with 20 nM FITC-sulfonamide probe and no protein constructs. Components were mixed by spinning down plates at 50 x g for 1 min and the plate was covered and incubated at room temperature for 1 hour, before analysis on a PHERAstar FS (BMG LABTECH) with fluorescence excitation and emission wavelengths (λ) of 485 and 520 nm, respectively, with a settling time of 0.3 s.

#### AlphaLISA displacement assay

The alphaLISA assays were performed as described previously^36^ using His_6_-BRD4^BD1^, His_6_-BRD4^BD2^ or 10xHis BRD4^Tandem^ and the biotinylated JQ1 probe. Assay conditions in the present work used were as follows: 100 nM bromodomain protein, 10 nM Bio-JQ1 probe, 25 µg/mL acceptor (nickel chelate) and donor (anti-his europium; both Perkin Elmer). All components were diluted to working concentrations in alphaLISA buffer (50 mM HEPES, 100 mM NaCl, 0.1% BSA, 0.02% CHAPS, pH 7.5). Bromodomain protein was co-incubated with test compounds using 384-well in AlphaPlates (PerkinElmer) in the absence or presence of DCAF16 (1 µM) for 1 hour, before adding the acceptor and donor beads simultaneously in a low light environment and incubating the pate at RT for a further hour. The plate was then read on a BMG Pherastar® equipped with an alphaLISA module. Data were normalised to a DMSO control and expressed as % bound vs log[concentration] of compound and analysed by non-linear regression, with extraction of binding affinity values (IC_50_) from the curves. Where applicable, K_D_ values were calculated from a titration of bromodomain protein on the same assay plate alone into the probe, as described previously^64^.

#### TR-FRET proximity assay

Stock solutions of reaction components including sulfo-Cy5-labelled DCAF16:DDB1ΔBPB:DDA1, sulfo-Cy5-labelled DCAF11:DDB1ΔBPB:DDA1 His_6_-BRD4^BD1^, His_10_-BRD4^BD2^, His_10_-BRD4^Tandem^, experimental compounds and LANCE Eu-W1024 Anti-His_6_ donor (PerkinElmer) were prepared in TR-FRET assay buffer (50 mM HEPES pH 7.5, 100 mM NaCl, 1 mM TCEP, 0.05% Tween-20). Two types of TR-FRET assay were performed: titration of compound into protein (‘complex formation assay’) and titration of sulfo-Cy5-labelled DCAF into BRD4 vs BRD4:compound (‘complex stabilization assay’). For the former, compounds were titrated 1:4 into 100 nM BRD4 and 100 nM Cy5-DCAF to a PerkinElmer OptiPlate-384 (white) to a final well volume of 16 μL. For the Complex stabilization assay, sulfo-Cy5-labelled DCAF16:DDB1ΔBPB:DDA1 or DCAF11:DDB1ΔBPB:DDA1 were titrated 1:4 and 1:3 respectively in TR-FRET assay buffer. Components were added to PerkinElmer OptiPlate-384 (white) to a final well volume of 16 μL. Final concentrations of 100 or 200 nM for BRD4 constructs and 0.5 µM or 1 µM for IBG1 respectively were used. LANCE Eu-W1024 anti-His_6_ donor and DMSO concentrations were kept constant across the plate for both assay formats at 2 nM and 0.5%, respectively. Background subtraction was performed with using concentration matched samples containing sulfo-Cy5-labelled DCAF complexes but not BRD4. Components were mixed by spinning down plates at 50 x g for 1 min and plates were covered and incubated at RT for 30 minutes. Plates were read on a PHERAstar FS (BMG LABTECH) with fluorescence excitation and dual emission wavelengths (λ) of 337 and 620/665 nm, respectively with an integration time between 70 – 400 μs. Data were processed in GraphPad Prism, curve fitting for the IBG1 curve was performed by setting the maximum as DMSO-only 5 µM sulfo-Cy5-labelled DCAF16:DDB1ΔBPB:DDA1 datapoint.

#### Analytical size exclusion chromatography

For DCAF16 experiments, DCAF16:DDB1ΔBPB:DDA1, BRD4^Tandem^ (residues 1-463), BRD4^BD^^1^ (His_6_ tag removed), BRD4^BD^^2^ (His_6_ tag removed), and IBG1 were incubated alone and in various combinations in buffer (20 mM HEPES, 150 mM NaCl, 1 mM TCEP, 2% DMSO, pH 7) on ice for 50 minutes. Final concentrations used for Fig. 4a and Extended Fig. 4a were 10 µM DCAF16:DDB1ΔBPB:DDA1, 5 µM BRD4^Tandem^, 25 µM IBG1 in 250 µL reaction volumes. Final concentrations used for Fig. 4b were 5 µM DCAF16:DDB1ΔBPB:DDA1, 5 µM BRD4^Tandem^, 5 µM BRD4^BD1^, 5 µM BRD4^BD2^, 12.5 µM IBG1 in 200 µL reaction volumes. Samples were run on a Superdex 200 Increase 10/300 gl column in 20 mM HEPES, 150 mM NaCl, 1 mM TCEP, pH 7.

For DCAF11 experiments, DCAF11:DDB1ΔBPB:DDA1, BRD4^Tandem^ (residues 43-463) and IBG4 were incubated alone and in various combinations in buffer (20 mM HEPES, 150 mM NaCl, 0.5 mM TCEP, 2% DMSO, pH 7.5) at final concentrations of 5 µM, 5 µM and 10 µM, respectively. Samples were run on a Superdex 200 Increase 10/300 gl column in 20 mM HEPES, 150 mM NaCl, 0.5 mM TCEP, pH 7.5.

For BRD4 intramolecular dimerization experiments, BRD4^Tandem^ (residues 43-463) and compounds were incubated in buffer (20 mM HEPES, 150 mM NaCl, 0.5 mM TCEP, 2% DMSO, pH 7.5) at final concentrations of 5 µM and 10 µM, respectively. Samples were run on a Superdex 200 Increase 10/300 gl column in 20 mM HEPES, 150 mM NaCl, 0.5 mM TCEP, pH 7.5.

#### Isothermal Titration Calorimetry

Titration experiments were performed with ITC200 instrument (Malvern) in 100 mM Bis-tris propane, 50 mM NaCl, 0.5 mM TCEP, pH 7.5 at 298 K. Protein samples were prepared by dialysing in buffer in D-Tube™ Dialyzer Midi, MWCO 6-8 kDa (Millipore). BRD4^Tandem^ (residues 43-459) was pre-incubated with either IBG1 or IBG3 at a 1:1.1 molar ratio for 30 mins at RT prior to titrations at a DMSO concentration of 2% (v/v). DCAF16:DDB1ΔBPB:DDA1 at 2% DMSO (v/v) was titrated into either pre-complexed BRD4^Tandem^:IBG1 or BRD4^Tandem^:IBG3. The titration consisted of 0.4 μL initial injection (discarded during data analysis) followed by 19 injections of 2 μL at 180 s intervals between injections. Data were fitted using a one-set-of-site binding model to obtain dissociation constant (K_D_), binding enthalpy (ΔH), and stoichiometry (N) using MicroCal PEAQ-ITC Analysis Software1.1.0.1262.

#### Cryo-EM sample and grid preparation

Protein complexes for cryo-EM were prepared by first co-incubating BRD4^Tandem^ (res 43-459) with IBG1 in 20 mM HEPES, 50 mM NaCl, 0.5 mM TCEP-HCl, 2% (v/v) DMSO, pH 7.5 for 10 mins at RT. DCAF16:DDB1ΔBPB:DDA1 was added to the mixture to give final concentrations of 14 µM BRD4^Tandem^, 14 µM DCAF16:DDB1ΔBPB:DDA1 and 35 µM IBG1 in a final reaction volume of 200 µL and incubated on ice for 50 mins. The sample was loaded onto a Superdex 200 Increase 10/300 GL column in 20 mM HEPES, 50 mM NaCl, 0.5 mM TCEP-HCl, pH 7.5. Due to incomplete complex formation and to avoid monomeric proteins, only the earliest eluting fraction containing the ternary complex was taken and concentrated to 4.8 µM. The complex was vitrified in liquid ethane using Vitrobot Mark IV (Thermo Fisher Scientific). Quantifoil R1.2/1.3 Holey Carbon 400 mesh gold grids (Electron Microscopy Sciences) were glow discharged for 60 s with a current of 35 mA under vacuum using a Quorum SC7620. The complex (3.5 µL) was dispensed onto the grid, allowed to disperse for 10 s, blotted for 3.5 s using blot force 3, then plunged into liquid ethane using a Vitrobot Mark IV (Thermo Fisher Scientific) with the chamber at 4 °C and 100% humidity.

#### Cryo-EM data acquisition

Cryo-EM data were collected on a Glacios transmission electron microscope (Thermo-Fisher) operating at 200 keV. Micrographs were acquired using a Falcon4i direct electron detector, operated in electron counting mode. Movies were collected at 190,000 x magnification with the calibrated pixel size of 0.74 Å/pixel on the camera. Images were taken over a defocus range of –3.2 µm to −1.7 µm with a total accumulated dose of 12.7e−/Å^2^ using single particle EPU (Thermo Fisher Scientific, version 3.0) automated data software. A total of 2,075 movies were collected in EER format and after cleaning up for large motion and poor CTF a total of 1,896 movies were used for further processing. A summary of imaging conditions and image processing is presented in Extended Data Table 2.

#### Cryo-EM image processing

Movies were imported into cryosparc^65^ (version 4.1.2) and the EER movie data was fractionated into 8 fractions to give a dose of 1.59 e−/Å^2^ per fraction. Movies were processed using patch motion correction and CTF correction then manually curated to remove suboptimal movies. Manual picking of 153 particles was performed on 20 micrographs, which were used for blob tuner with minimum and maximum diameters of 70 and 130 Å, respectively. 12,579 particles were picked by blob tuner, extracted with a box size of 324 pix (240 Å) and run through initial 2D classification. Good classes with diverse views were selected and used as templates for template picking on 1,895 movies. Picks were inspected and curated, and 1.35 million particles were extracted with box size 324 pix and used for 2D classification. Particles from the well-resolved, diverse classes were used for ab-initio reconstruction with 3 classes. One class contained primarily empty DDB1ΔBPB and a second class contained biased views upon testing of the particle set with 2D re-classification, leading to smeared maps. The third class unambiguously contained density corresponding to DDB1ΔBPB, two bromodomains, and density likely corresponding to DCAF16 between them. Particles belonging to the second and third class were run through heterogenous refinement. The best class yielded a map into which DDB1ΔBPB and 2 bromodomains could be placed with confidence. To improve the resolution, movies were re-imported in cryosparc and fractionated into 18 fractions to give a lower dose of ∼0.7 e/Å^2^ per fraction. Fifty templates for particle picking were generated using the create templates job with the input map from the previous heterogeneous refinement. The templates were used in the template picker to pick particles from 1,132 curated movies with a minimum CTF fit resolution cut-off of 3.5. Picks were curated with thresholds of NCC score > 0.4, local power >368 and <789, resulting in 564,575 particles that were extracted with a box size of 324 pixels and used for ab-initio reconstruction with 4 classes. Resulting classes were subjected to a heterogeneous refinement, with one class clearly containing all components of the complex and the others either junk, DDB1ΔBPB alone or biased views. The map and particles (192,014) from the best class were used for homogenous refinement with the dynamic mask threshold set to 0.5. Local refinement with a dynamic map threshold of 0.5 produced a map with a gold-standard Fourier shell correlation (GSFSC) resolution of 3.77 Å at cut-off 0.143. The workflow, GSFSC curve, local resolution estimation, angular distribution plot, and posterior position directional distribution plot are presented in Extended Data Fig. 4.

#### Cryo-EM model building

DDB1ΔBPB, BRD4^BD1^ and BRD4^BD2^ extracted from PDB entries 5FQD^28^, 3MXF^66^ and 6DUV, respectively, were manually placed into the map in Coot^67^ (version 0.9.8.1) by rigid body fitting. Despite co-purifying with DCAF16 and DDB1ΔBPB, we did not see density for DDA1, as was observed in another DDB1-substrate receptor structure from a recent publication^27^. Correct placement of each bromodomain was aided by manual inspection of residues Asn93 and Gly386 in equivalent positions in the ZA loops of BD1 and BD2, respectively. In one bromodomain, this position was facing solvent while in the other it was at a protein-protein interface with density corresponding to DCAF16. Given that mutation of Gly386 to Glu prevents degradation of BRD4 by IBG1 (Fig. 3i), BD2 was placed in the position where Gly386 was adjacent to the DCAF16 density. The BD2 ZA loop is 3 residues longer than the BD1 ZA loop, further confirming the correct positioning of each domain based on the map around these positions. Both bromodomains were joined onto a single chain designation. Initial restraints for IBG1 were generated using a SMILES string with eLBOW^68^, then run through the GRADE webserver (Grade2 v1.3.0). IBG1 was fit into density by overlaying the JQ1 moiety with its known binding mode in either the BRD4^BD1^ or BRD4^BD2^. Positioning the ligand in BD2 was compatible with electron density, whereas positioning in BD1 caused a clash with DCAF16 due to the rigid linker. DCAF16 was built using a combination of models from ColabFold^69,70^ (version 1.3), ModelAngelo^71^ (version 0.2.2) and manual building in Coot. ColabFold correctly predicted the α5 and α6 helices that bind the DDB1 central cavity while ModelAngelo correctly built the 4-helical bundle of α3, 4, 7 and 8, as well as α6 in the DDB1 cavity. Correctly built parts of the models were combined, and the structure was refined with rounds of model building in Coot, fitting with adaptive distance restraints in ISOLDE^72^ (version 1.6) and refinement with Phenix (version 1.20.1-4487) real-space refinement^73,74^. Figures were generated in ChimeraX^75^ (version 1.6) and The PyMOL Molecular Graphics System^76^ (version 2.5.2), Schrödinger, LLC.

